# Discovery of an orally available potent ER aminopeptidase 1 (ERAP1) inhibitor that enhances anti-tumor responses and limits inflammatory autoimmunity in vivo

**DOI:** 10.1101/2025.11.17.686761

**Authors:** Christopher P. Tinworth, Justyna Wojno-Picon, Michael Adam, Stephan Gade, Ashley P. Hancock, David J. Hirst, Jonathan P. Hutchinson, Semra J. Kitchen, Despoina Koumantou, Jonathan D. Lea, Stephanie Lehmann, John Liddle, Richard Lonsdale, Margarete Neu, Leng Nickels, Alex Phillipou, James E. Rowedder, Paul Rowland, Jessica L. Schneck, Paul T. Scott-Stevens, Hester Sheehan, Michael Steidel, Chloe L. Tayler, Ioannis Temponeras, Kevin Thang, David F. Tough, Giovanni Vitulli, Ian D. Wall, Robert J. Young, Nico Zinn, Simon Peace, Efstratios Stratikos

## Abstract

Endoplasmic reticulum aminopeptidase 1 (ERAP1) is an intracellular enzyme that can regulate immune responses primarily by proteolytically processing peptides before loading and presentation on the cell surface by major histocompatibility class I molecules (MHC-I). ERAP1 activity can either reduce the immunogenicity of cancer cells by over-trimming cancer-associated antigenic peptides or contribute to autoimmunity by generating self-antigenic peptides. As a result, ERAP1 inhibition has emerged as a tractable approach for cancer immunotherapy and specific classes of autoimmunity. Here, we describe the discovery, after hit-to-lead optimization, of a potent and selective ERAP1 inhibitor based on the pyrrolidine 3-carboxylic acid scaffold that targets the regulatory allosteric site. The compound has favourable *in vivo* pharmacokinetics, including oral bioavailability, and can regulate the immunopeptidome of cancer cells and enhance cancer cell antigenicity *in vivo* in a dose-dependent manner, controlling tumor growth. In addition, when administered in the murine collagen-induced arthritis model, it does not induce any exacerbation of autoimmune responses but rather results in a dose-dependent therapeutic benefit. Our results demonstrate that ERAP1 inhibition can constitute a tractable approach to modulating immune responses for therapeutic applications, providing mechanistic insight and a valuable lead and *in vivo* tool for further drug development efforts and for interrogating ERAP1 biology.

## Introduction

T cells of the adaptive immune system recognize, via the T cell receptor (TCR), infected or aberrant cells on the basis of small peptides bound onto major histocompatibility complex (MHC) molecules on the cell surface. MHC class I molecules (MHC-I) fold inside the endoplasmic reticulum (ER) by binding peptides that originate from intracellular proteins and then translocate to the cell surface.^1^ TCRs recognize these MHC-I/peptide complexes thus gaining insight on the proteomic content of the cell.^2^ The presence of peptides that belong to pathogens or aberrantly expressed proteins elicits recognition by T cells and signifies infection or cancerous transformation, inducing molecular responses that lead to the killing of the target cell. The MHC-presented peptides, collectively named the immunopeptidome, are generated by complex proteolytic cascades that often include the proteasome or its immune system counterpart, the immunoproteasome.^3^ Peptide fragments from intracellular proteins are then translocated into the ER by a specialized transporter called transporter associated with antigen presentation (TAP).^4^ However, since MHC-I molecules bind peptides with strong selectivity for length, many ER imported peptides require additional processing to enable binding. To achieve this, ER-resident aminopeptidases trim those peptides, optimizing their length for binding onto nascent MHC-I molecules (on average 9 amino acids length).^5^ One such aminopeptidase, ER aminopeptidase 1 (ERAP1) is highly specialized for this task and can efficiently prepare correct-length peptides for MHC-I binding, although it can also over-trim some peptides to lengths that preclude MHC-I binding, essentially destroying their antigenic potential.^6,7^ Although ERAP1 is typically retained into the ER, it can be recruited to endosomes in dendritic cells where it can process peptides from internalized proteins for cross-presentation, a process whereby peptides from extracellular proteins are presented in association with MHC-I molecules.^8^ Other functions have also been proposed for ERAP1, although these are less explored. For instance, under certain inflammatory conditions, ERAP1 can be secreted by immune cells, enhancing innate immune responses through the activation of the inflammasome complex.^9,10^

Several studies have shown that genetic down-regulation of ERAP1 in cells, or its mouse orthologue endoplasmic reticulum aminopeptidase associated with antigen processing (ERAAP), can strongly enhance the antigenicity of cells.^7^ These effects have been primarily attributed to changes in the cellular immunopeptidome, the sum of presented peptides by MHC-I molecules. Several molecular and cellular mechanisms have been proposed that may synergize depending on the inflammatory and cellular context: i) presentation of longer length, unstable peptides that are recognized as foreign by TCRs^7^ or fail to productively engage inhibitory receptors on natural killer (NK) cells,^11,12^ ii) presentation of a different repertoire of good-binding peptides that is recognized by existing T cells or can lead to the maturation of new effector T cells^13^ and iii) reduced presentation of peptides derived from the MHC-I signal sequence by HLA-E.^14,15^ ERAP1 expression levels regulated by expression quantitative trait loci have been reported to regulate cancer immunotherapy responses,^16^. Additionally, ERAP1 activity has been correlated with CD8^+^ T cell tumor infiltration^17^ and ERAP1 knockdown has been shown to synergize with other agents in promoting the efficacy of cancer immunotherapy.^12,15^ Thus, an increasing amount of evidence suggests that ERAP1 may be a tractable target for cancer immunotherapy approaches.

MHC (HLA in humans) are amongst the most polymorphic genes in humans with over 20,000 different alleles identified to date.^18^ This polymorphic variation is primarily focused on the antigenic peptide binding groove and determines the breadth of possible peptides that can be presented by each individual, defining the context for variable immune responses within human populations. A subset of inflammatory diseases of autoimmune aetiology, called MHC-I-opathies, is very strongly correlated with specific HLA class I alleles, likely due to the presentation of self-antigenic peptides or to changes in folding of MHC-I/peptide complexes.^19^ ERAP1 is also polymorphic, and ERAP1 coding single nucleotide polymorphisms (SNPs) have been correlated, often in epistasis with MHC-I haplotypes, with predisposition to MHC-I-opathies, most notably Ankylosing Spondylitis.^20^ ERAP1 coding SNPs affect enzymatic activity^21^ and are found in human populations in allotypes with variable penetrance and highly variable enzymatic properties.^22,23^

Several inhibitors of ERAP1 have been reported, discovered either by rational design or screening campaigns.^24,25^ A group of sub-micromolar ERAP1 inhibitors reported target the Zn(II)-containing active site of the enzyme and utilize phosphinic pseudopeptide, diaminobenzoic acid, or α-hydroxy-β-amino acid scaffolds, but present limitations in selectivity possibly due to the highly conserved active site among M1 aminopeptidases.^26–28^ Allosteric sites that allow for superior selectivity have been also explored, leading to nanomolar potency inhibitors targeting one of two allosteric sites found to be occupied by buffer components in a high-resolution structure of ERAP1.^29–32^ The malate allosteric site has attracted significant interest due to its functional significance in recognizing the *C*-terminus of ERAP1’s peptidic substrates and regulating length selectivity.^33^ Inhibitors targeting the active site or allosteric sites have also been shown to be able to modulate the immunopeptidome of cancer cell lines, albeit in distinct patterns.^34–36^ Development of selective small molecules targeting ERAP1 has significantly accelerated, with the first clinical candidate GRWD5769 entering clinical trials in 2023.^37^

Here, we report the development of an orally available, potent ERAP1 inhibitor targeting the regulatory site of the enzyme^33^ optimized from a series of 3-pyrrolidine carboxylic acids originating from a previously described high-throughput screen (HTS).^31^ This compound can regulate the immunopeptidome of cancer cells and regulate cellular immunogenicity in two distinct *in vivo* models with favourable therapeutic outcomes. Our results provide proof-of-concept that targeting the regulatory site of ERAP1 has significant therapeutic potential and report the first potent *in vivo* tool for interrogating ERAP1 biology and guide further drug development efforts.

## Results

Following a previously described HTS campaign, (3*S*,4*S*)-1-(3-cyano-6-methylpyridin-2-yl)-4-isopropylpyrrolidine-3-carboxylic acid **1** was identified as a ligand efficient hit molecule with micromolar inhibition of ERAP1, as measured using a previously described assay monitoring the cleavage of 9-mer antigenic peptide YTAFTIPSI to 8-mer TAFTIPSI (**Table 1**).^31,38^ The corresponding enantiomer acid **2** was prepared, which was observed to have approximately equivalent binding potency. Both compounds were found to have high kinetic solubility, consistent with the carboxylic acid moiety and low lipophilicity as measured by ChromlogD_7.4_.

**Table 1.**
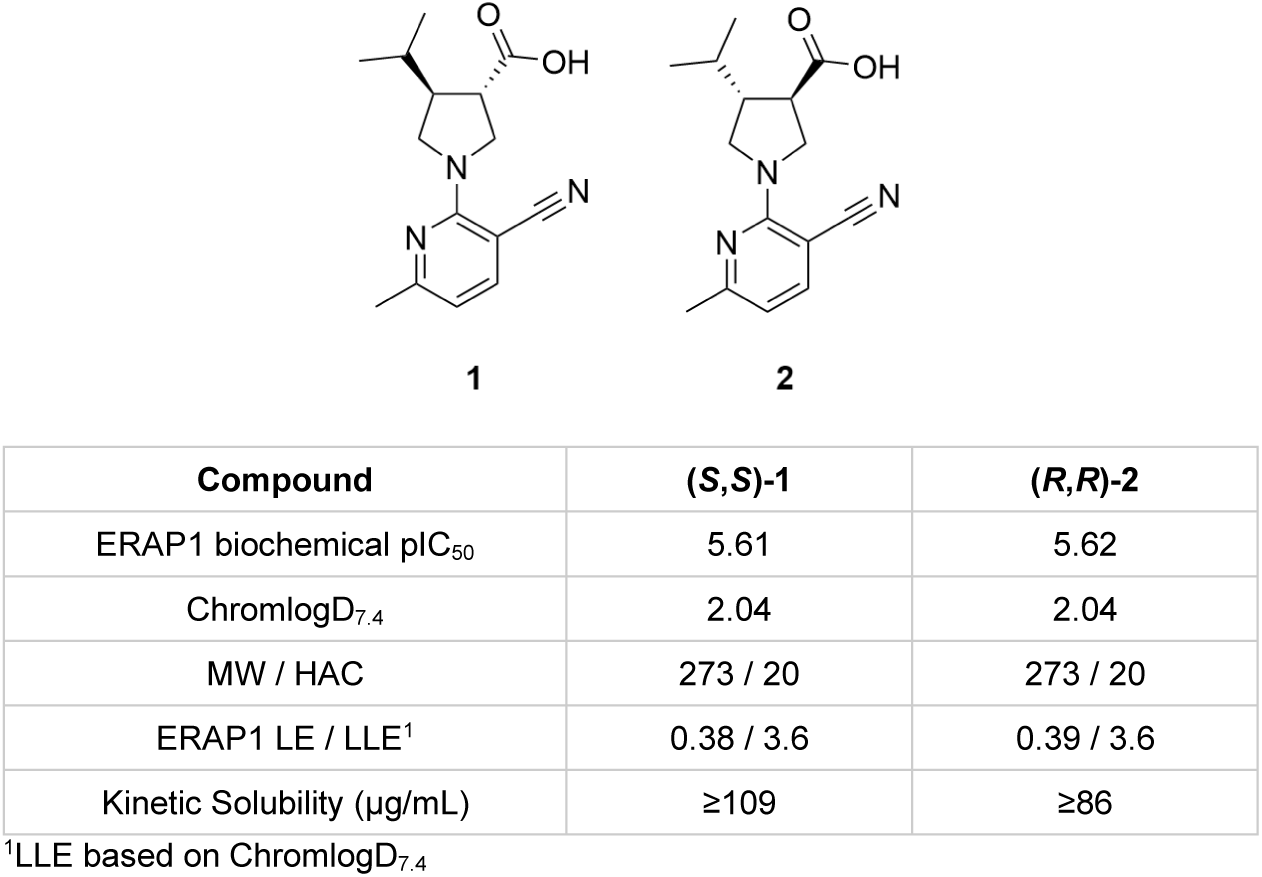
Profiling of hit enantiomers **1** and **2**

| Compound | (*S,S*)-**1** | (*R,R*)-**2** |
| --- | --- | --- |
| ERAP1 biochemical pIC<sub>50</sub> | 5.61 | 5.62 |
| ChromlogD<sub>7.4</sub> | 2.04 | 2.04 |
| MW / HAC | 273 / 20 | 273 / 20 |
| ERAP1 LE / LLE<sup>1</sup> | 0.38 / 3.6 | 0.39 / 3.6 |
| Kinetic Solubility (μg/mL) | ≥109 | ≥86 |

<sup>1</sup>LLE based on ChromlogD<sub>7.4</sub>

A co-crystal structure for both compounds **1** and **2** with ERAP1 was generated and is presented in **Figure 1**. This confirmed that the hit molecules target the ERAP1 regulatory site, as previously observed for a natural product modulator identified within our laboratories.^31^ In both co-crystal structures, the carboxylic acid makes charge interactions with Tyr684/Lys685/Arg807, while the isopropyl substituent occupies a proximal hydrophobic pocket. The relative configuration of the two pyrrolidine substituents results in similar spatial arrangement of the acid and isopropyl groups, accounting for the similar potency observed between the two enantiomers. The nitrile group from the aminopyridine forms a hydrogen bonding interaction with Gln881.

**Figure 1.**
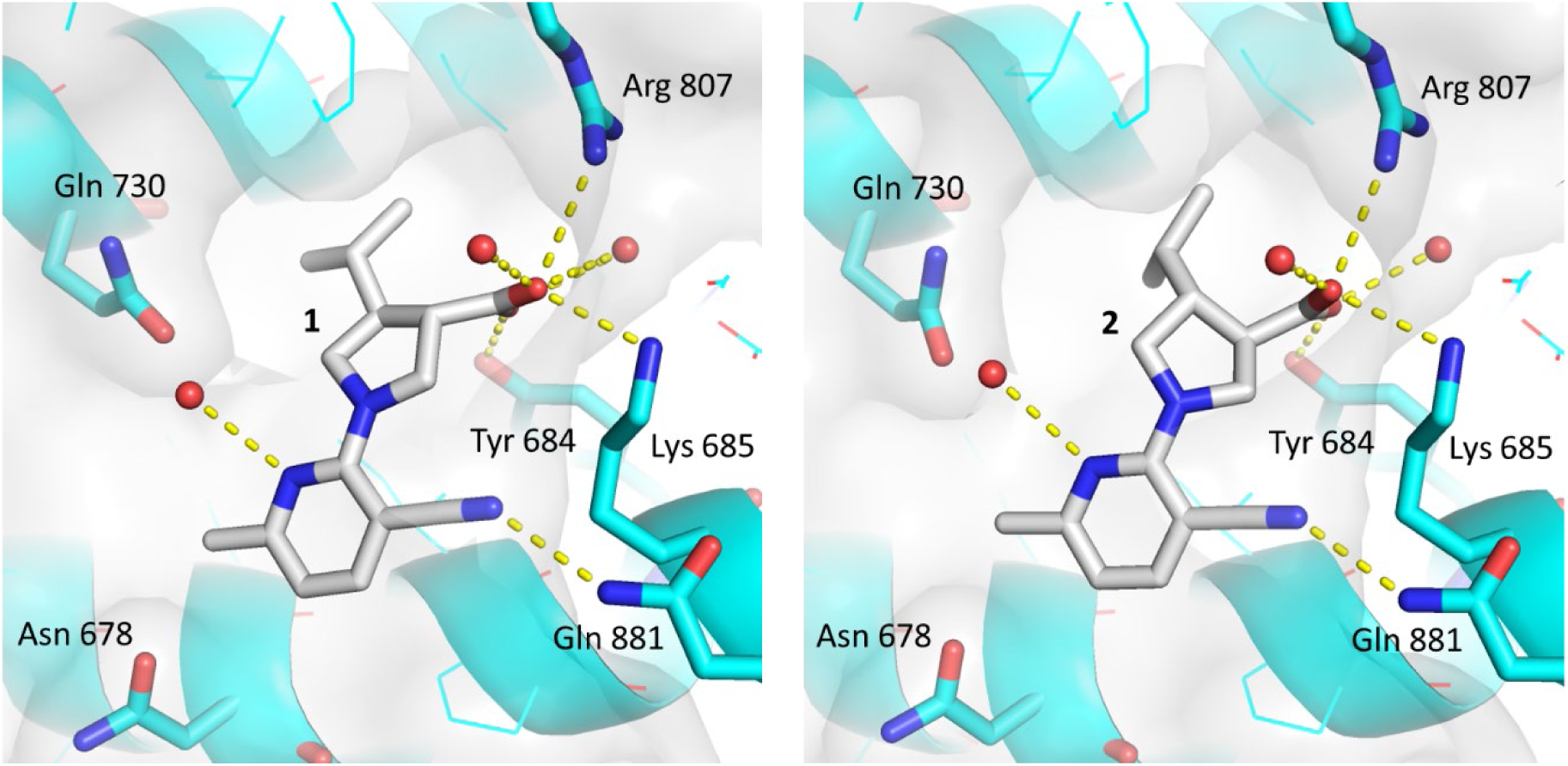
Co-crystal structure of hit pyrrolidine 3-carboxylic acid enantiomers **1** (left) and **2** (right) with ERAP1

The co-crystal structures of compounds **1** and **2** indicated that the 6-methylpyridine was in close proximity to both and Asn678/Gln730. It was thought that inclusion of hydrogen bond donors or acceptors in this region could gain productive hydrogen bonding interactions with either of these sidechains to further increase binding potency, as outlined in **Table 2**. Based on the similar binding potency of the two enantiomers, compounds were prepared as racemates, with enantiomeric separation or chiral synthesis carried out for key analogues.^39^ For in-line comparison, compound **3** was prepared, a racemic mixture of compounds **1** and **2**. Introduction of a hydrogen bond acceptor in 2-methoxypyridine **4** led to no improvement in potency compared to compound **3**. However, introduction of a hydrogen bond donor in 2-aminopyridine **5** led to a significant improvement in biochemical potency, ligand efficiency (LE) and lipophilic ligand efficiency (LLE). Cellular activity of compound **5** was profiled using a previously described assay based on detection of ERAP1-dependent antigenic epitope SIINFEKL in HeLa cells,^31^ and was found to inhibit antigen presentation at micromolar level (**Table 2**).

**Table 2.** SAR of 6-pyridyl substituents to probe proximal hydrogen bonding interactions

| Compound | R | ERAP1 pIC<sub>50</sub> | HeLa antigen presentation pIC<sub>50</sub> | ChromlogD<sub>7.4</sub> | LE | LLE | Kinetic Solubility (μg/mL) |
| --- | --- | --- | --- | --- | --- | --- | --- |
| (±)-3 | Me | 5.67<sup>1</sup> | nt | 2.05 | 0.39 | 3.6 | ≥102 |
| (±)-4 | OMe | 5.53 | nt | 2.39 | 0.36 | 3.1 | ≥118 |
| (±)-5 | NH<sub>2</sub> | 6.31 | 5.48<sup>1</sup> | 0.94 | 0.43 | 5.4 | ≥118 |
| (±)-6 | NH<sup>i</sup>Pr | 7.45 | 6.68 | 2.68 | 0.44 | 4.8 | ≥109 |
| (±)-7 | NH<sup>c</sup>Pr | 7.59 | 6.86 | 2.38 | 0.45 | 5.2 | ≥145 |
| (*R,R*)-8 | NH<sup>c</sup>Pr | 7.43 | 6.99 | 2.52 | 0.44 | 4.9 | ≥106 |
| (*S,S*)-9 | NH<sup>c</sup>Pr | 7.13 | 6.80 | 2.44 | 0.43 | 4.7 | ≥96 |

<sup>1</sup>n=2. nt = not tested**Figure 2.** Co-crystal structure of cyclopropylamine derivative **8** with ERAP1.

The co-crystal structures of compounds **1** and **2** indicated potential close proximity of the appended amine to the proximal hydrophobic pocket. Isopropyl amine **6** afforded the desired biochemical potency improvements, while also improving cellular activity (**Table 2**). Cyclisation of the isopropyl to cyclopropyl in compound **7** retained binding activity of isopropylamine **6** with improved LLE. Cyclopropyl amine **7** was separated to give both (*R,R*)-enantiomer **8** and (*S,S*)-enantiomer **9**, which revealed a minor biochemical preference of the (*R,R*)-enantiomer. A co-crystal structure of compound **8** with ERAP1 supports that the 6-aminopyridyl group forms the desired hydrogen bonding interaction with Asn678 (**Figure 2**).

**Figure 2.**
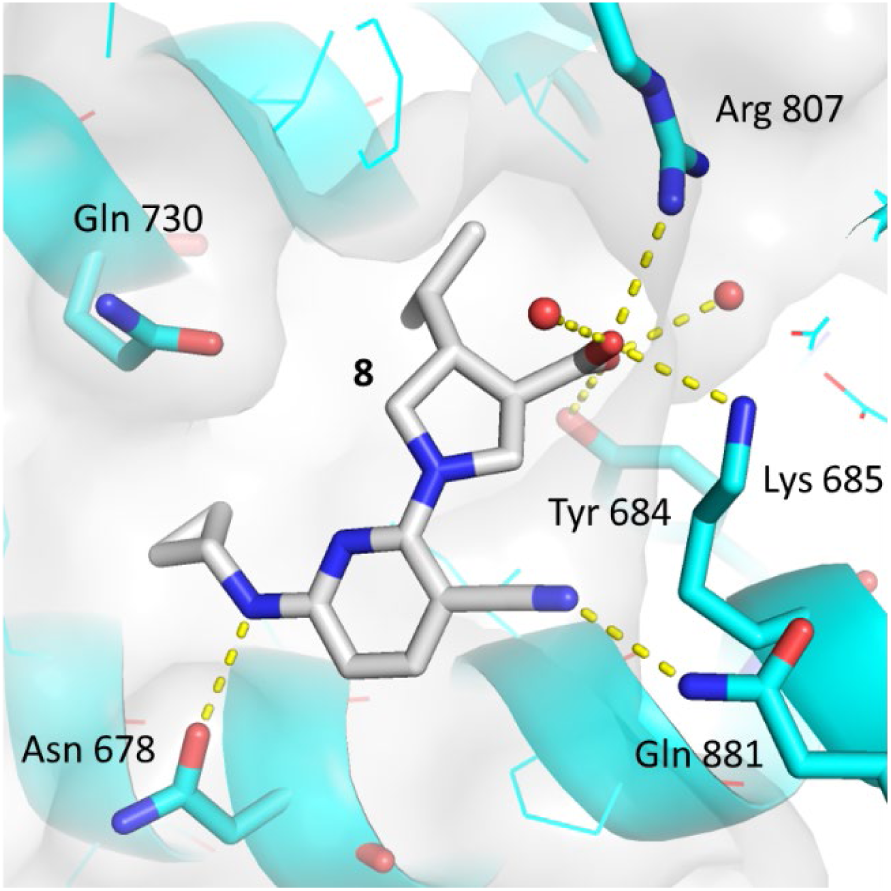
Co-crystal structure of cyclopropylamine derivative **8** with ERAP1.

Further lipophilic analogues occupying the hydrophobic pocket were explored, outlined in **Table 3**. Cyclopentyl and cyclohexyl analogues **10** and **11** resulted in further increases in binding potency and cellular pIC_50_>7, though with no LLE increase compared to cyclopropyl analogue **9**. Introduction of polarity into the more potent cyclopentyl analogues through 3-aminotetrahydrofurans **12** and **13** demonstrated a significant LLE benefit for the preferred (*S*)-isomer **13** however cellular activity was reduced. Cyclohexyl analogues such as difluorocyclohexyl compound **14** increased LLE with retained cellular activity, while the low lipophilicity of tetrahydropyran **15** proved detrimental to cellular potency. A compromise in biochemical LLE and cellular potency was observed with aminomethyl cycloalkanes, where cyclopropylmethylamine **16** demonstrated high LE/LLE values and retained cellular potency, while reduced LE/LLE was observed with cyclobutylmethylamine **17** .

**Table 3.** SAR of aminopyridine substituents to occupy proximal hydrophobic pocket

| Compound | R | ERAP1<br>pIC<sub>50</sub> | HeLa antigen<br>presentation pIC<sub>50</sub> | ChromlogD<sub>7.4</sub> | LE | LLE | Kinetic<br>Solubility<br>(μg/mL) |
| --- | --- | --- | --- | --- | --- | --- | --- |
| (±)-10 |  | 8.27 | 7.44 | 3.42 | 0.45 | 4.9 | ≥129 |
| (±)-11 |  | 8.52 | 7.63 | 3.93 | 0.45 | 4.6 | ≥112 |
| (±)-12 |  | 6.57 | 5.65 | 1.50 | 0.36 | 5.1 | ≥96 |
| (±)-13 |  | 8.17 | 6.66 | 1.67 | 0.45 | 6.5 | ≥136 |
| (±)-14 |  | 8.34 | 7.37 | 3.42 | 0.41 | 4.9 | ≥167 |
| (±)-15 |  | 7.93 | 5.86 | 1.86 | 0.42 | 6.1 | ≥140 |
| (±)-16 |  | 8.05 | 7.54 | 2.90 | 0.50 | 5.2 | ≥101 |
| (±)-17 |  | 8.15 | 7.27 | 3.52 | 0.45 | 4.6 | ≥116 |

Based on the crystal structure of compound **9** (**Figure 2**) a further hydrophobic region at the *ortho-*position to the nitrile was identified for growth and potential to further balance physicochemical properties. While introduction of a methyl group in this region with compound **18** gave little benefit to biochemical potency when compared to compound **14**, cellular potency increased to pIC_50_ >8 (**Table 4**). Separation of the enantiomers of compound **18** again demonstrated a binding preference for (*R*,*R*)-enantiomer **19** over (*S*,*S*)-enantiomer **20**. Further variation of the amine group identified 4-methyltetrahydropyran **21** as a potent and cell active binder with reduced lipophilicity compared to compound **19**. Profiling *in vitro* clearance from human hepatocytes for compounds **19** and **21** indicated improved unbound intrinsic clearance (CL_int,u_) for compound **21** (**Supplementary Table 1**). Crystallography of compound **21** bound to ERAP1 indicated that the key interactions from compound **9** were retained, while an additional interaction was observed between the THP oxygen atom and Gln730 (**Figure 3**).

**Table 4.** SAR following nitrile *ortho-*methylation with variable amine substituents

| Compound | R | ERAP1<br>pIC<sub>50</sub> | HeLa antigen<br>presentation pIC<sub>50</sub> | ChromlogD<sub>7.4</sub> | LE | LLE | Kinetic<br>Solubility<br>(μg/mL) | Human<br>hepatocyte<br>CL<sub>int,u</sub><br>(mL/min/kg) |
| --- | --- | --- | --- | --- | --- | --- | --- | --- |
| (±)-**18** |  | 8.42 | 8.11 | 3.64 | 0.41 | 4.8 | ≥108 | nt |
| (*R,R*)-**19** |  | 8.55<sup>2</sup> | 8.09 | 3.67 | 0.40 | 4.9 | ≥126 | 68 |
| (*S,S*)-**20** |  | 8.16 | 7.39 | 3.72 | 0.39 | 4.4 | ≥112 | nt |
| (*R,R*)-**21**<br>(GSK235) |  | 8.45 | 7.25<sup>1</sup> | 2.44 | 0.41 | 6.0 | ≥201 | 39 |

<sup>1</sup>n=2. nt = not tested

**Figure 3.**
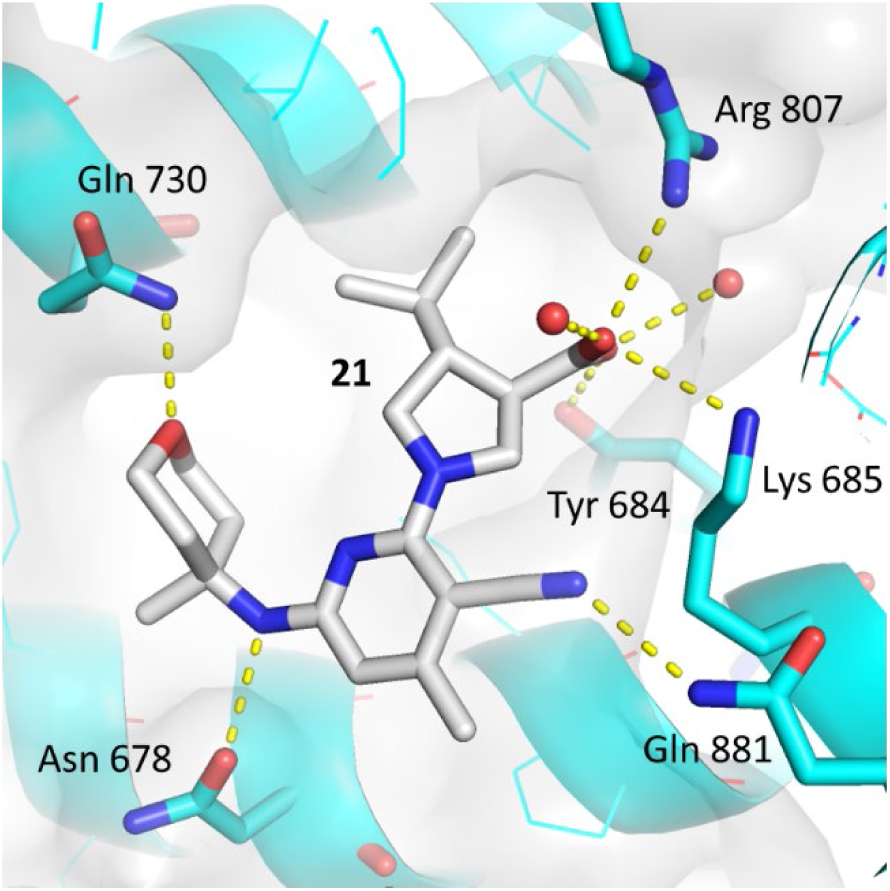
Co-crystal structure of ERAP1 and compound **21**.

Due to their favourable cellular potency and physicochemical properties, compounds **19** and **21** were progressed to mouse oral pharmacokinetic (PK) studies to assess their potential as *in vivo* tool molecules. Despite lower cellular activity, compound **21** was found to have improved unbound oral exposure in mouse compared to compound **19** (**Table 5**). Therefore compound **21**, otherwise known as GSK235, was progressed to further *in vivo* studies.

**Table 5.** Oral PK of compounds **19** and **21** following administration to naïve BALB/c mice (n=3).

| Compound | Dose (mg/kg) | C<sub>max</sub> (ng/mL) | AUC<sub>0-t<sub>l</sub></sub> (ng.h/mL) | f<sub>u,blood</sub> | DNAUC<sub>0-t<sub>l</sub></sub> (ng.h/mL/mg/kg) |
| --- | --- | --- | --- | --- | --- |
| **19** | 10 | 950 | 2989 | 0.002 | 0.602 |
| **21** (GSK235) | 10 | 8516 | 15440 | 0.005 | 7.43 |

AUC = Area under the curve. DNAUC = Dose-normalised AUC

Extended profiling and developability screening of compound **21** is described in **Table 6**. Compound **21** is a potent ERAP1 inhibitor with high LE/LLE which retains inhibitory activity against mouse ERAAP. Selectivity of over 1000x for the two most homologous enzymes ERAP2 and IRAP was observed for compound **21** based on previously reported assays.^31^ During the course of optimisation, the original HeLa antigen presentation assay protocol was adjusted such that the data was normalized against a pharmacological tool compound (see **Materials and Methods**).^38^ Based on this the cellular potency of compound **21** was adjusted to pIC_50_ = 7.05, with unbound cellular pIC_50_ = 7.26, accounting for binding to the cell assay media. Consistent with the low lipophilicity of compound **21**, solubility was high based on amorphous solubility and crystalline FaSSIF solubility, with moderate *in vitro* permeability. Compound **21** was found to be inactive in Ames and MLA assays; while extended cross-screening across a panel of known liability targets indicated no major flags, other than weak inhibitory activity against OATPB1 and BSEP transporters. No activity against the hERG channel was observed.

**Table 6.** Profile of compound 21 (GSK235)

| Compound | **21** (GSK235) |
| --- | --- |
| ERAP1 biochemical pIC<sub>50</sub> (human, YTAFTIPSI) | 8.45 |
| ERAAP biochemical pIC<sub>50</sub> (mouse, YTAFTIPSI) | 7.59 |
| ERAP2 / IRAP biochemical pIC<sub>50</sub> | 4.02 / 4.28 |
| HeLa antigen presentation pIC<sub>50</sub> total / unbound | 7.05 / 7.26 |
| MW / ChromlogD<sub>7.4</sub> / Acidic pK<sub>a</sub> | 387 / 2.44 / 4.01 |
| Solubility (µg/mL): CAD (amorphous) / FaSSiF (crystalline) | ≥201 / 675 |
| AMP / MDCK P<sub>app</sub> (nm/s) | 28 / 46 |
| Hepatocyte CL<sub>int</sub> m/r/h (mL/min/g tissue) | 0.86 / 2.20 / 1.43 |
| Fraction unbound: media; blood (m/r/h) | 0.61; 0.005 / 0.002 / 0.006 |
| hERG pIC<sub>50</sub> / Ames / MLA | <4 / Negative / Negative |
| Secondary pharmacology cross-screening panel | OATP1B1 pIC<sub>50</sub> = 5.0, BSEP pIC<sub>50</sub> = 4.9 |

## Profiling of Compound 21 (GSK235)

Compound **21** was progressed to profiling in two *in vivo* mechanistic models with the purpose of investigating the linkage between target engagement and downstream pharmacology whilst also providing insight into potential liabilities. Firstly, the impact of modified tumor immuno-visibility resulting from modulation of MHC proteins was evaluated with compound **21** in an oncology setting through (i) evaluation of tumor-relevant peptide substrates, such as MART1^40^ (ii) comparison of inhibition with knock-out in the CT26 mouse tumor cell line (iii) inhibition of *in vivo* tumor growth with the syngeneic CT26 model.

Secondly, compound **21** was evaluated in an inflammatory autoimmune setting, based on the strong genetic associations between ERAP1 polymorphisms and immune-mediated diseases such as ankylosing spondylitis and psoriasis. These associations imply: (1) ERAP1 may participate in the generation of antigenic peptides recognised by pathogenic T cells in these diseases and (2) inhibition of ERAP1 may interfere with the generation of these pathogenic peptides and provide benefit in disease. However, alteration of the immunopeptidome by ERAP1 inhibition could have the unintended effect of generating new self-antigen epitopes with the potential of generating or exacerbating an autoimmune response. Such a risk has been implied by published studies of ERAAP knock-out.^7,41^

Mouse collagen-induced arthritis (CIA) was selected as an *in vivo* model for this purpose due to the presence of both an autoimmune T and B cell response and inflammation. Such a setting would be expected to provide an environment rich in co-stimulatory signals capable of enabling a T cell response that might be triggered by ERAP1-inhibition driven new self-peptides. Standard pathology and scoring endpoints for CIA were supplemented with several measures of immune activation to investigate how ERAP1 inhibition impacted autoimmunity and inflammation.

Doses of 30, 90 and 270mg/kg BID were selected based on preliminary mouse PK studies (**Table 5**), targeting maximal inhibition and a dynamic concentration range. Herein we describe the PK, estimation of target engagement based on *in vitro* findings, *in vivo* pharmacology and impact on pathological endpoints.

### a) Estimation of unbound drug concentration *in vivo*

The most comprehensive and representative longitudinal PK data were measured in the CIA study and are presented in **Figure 4**. PK sampling was undertaken on days 18, 26 and 35 at 1h, 3h and 12h, immediately before the second daily dose, and at 24h, immediately before the first dose on the following day. The unbound concentration of compound **21** was calculated from the total blood concentration and fraction unbound data in **Table 6**. The concentrations observed were largely dose-proportional, with some variation across the study, for example lower concentrations were noted in some groups on day 26. The ‘peak’ and ‘trough’ concentrations were assessed to be at 1h and 24h and varied from approximately 3-100nM for the 30mg/kg group, 20-700nM for the 90mg/kg group and 200-2000nM for the 270mg/kg group.

**Figure 4.**
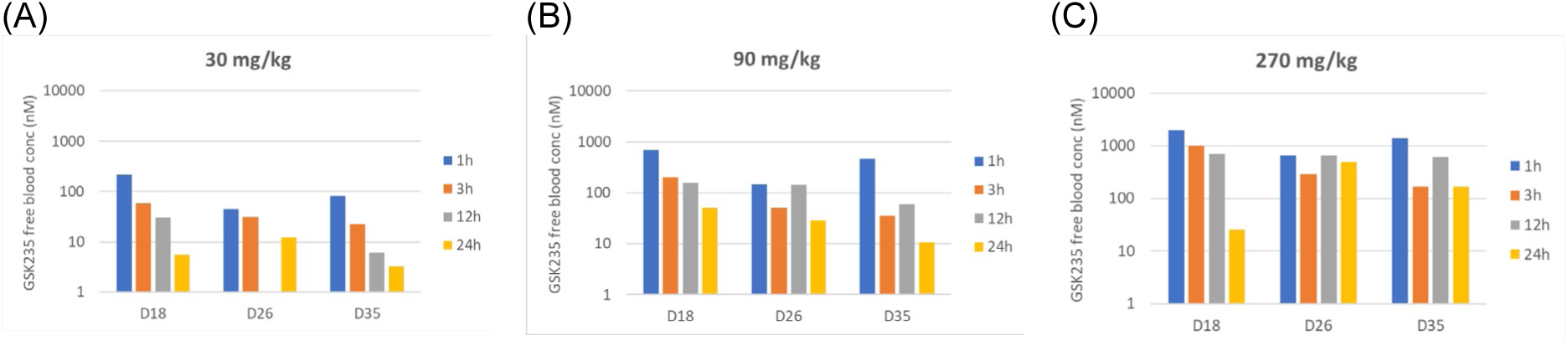
Unbound concentration of GSK235 based on the CIA study PK on days 18, 26 and 35 following oral BID dosing at (A) 30mg/kg, (B) 90mg/kg and (C) 270mg/kg.

### b) In vitro target engagement and estimation of in vivo target engagement

The murine ERAAP potency of compound **21** against YTAFTIPSI and tumor-relevant MART1 EAAGIGILTV^40^ substrates was measured and is presented in **Table 7**.^42^ It is noteworthy that the depth of response and IC_50_ are substrate dependent, illustrating the challenge and complexity of assessing the degree of target engagement.

**Table 7.**
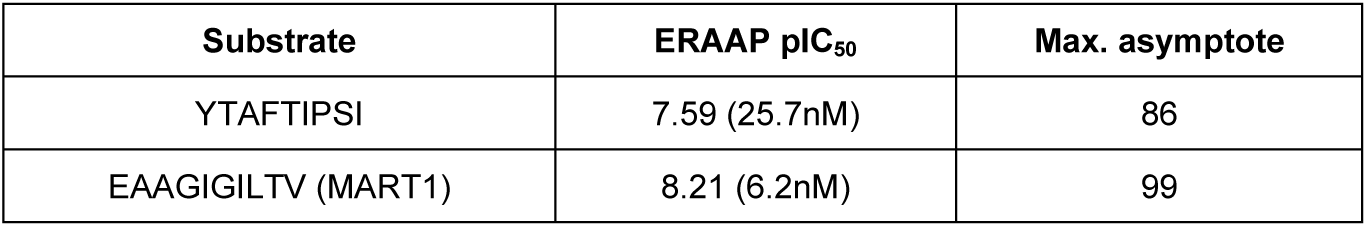
Mouse ERAAP potencies of compound **21** for distinct peptide substrates.

| Substrate | ERAAP pIC<sub>50</sub> | Max. asymptote |
| --- | --- | --- |
| YTAFTIPSI | 7.59 (25.7nM) | 86 |
| EAAGIGILTV (MART1) | 8.21 (6.2nM) | 99 |

The ERAAP potency and maximum asymptote (25.7nM, 86%; YTAFTIPSI substrate) were combined with unbound concentrations at 1 and 24 hours from the CIA study to provide preliminary estimations of peak and trough target engagement (**Table 8**). The predicted ERAAP inhibition varied from 53-73% (peak) declining to 14-31% (trough) at the 30mg/kg dose group escalating to 80-83% (peak, C_max_) to 43-79% (trough, C_min_) for the 270mg/kg group. It is important to note that the YTAFTIPSI is the most conservative choice of substrate potency and the maximum inhibition achievable in this prediction would be 86%.

**Table 8.** Peak and trough target engagement (TE) estimations derived from CIA PK data and ERAAP potency (YTAFTIPSI substrate)

| Estimated TE | Mouse oral dose |  |  |
| --- | --- | --- | --- |
|  | 30mg/kg BID | 90mg/kg BID | 270mg/kg BID |
| C<sub>max</sub> | 53-73% | 69-80% | 80-83% |
| C<sub>min</sub> | 14-31% | 29-55% | 43-79% |

The impact of compound **21** against a wider variety of immunopeptides was assessed by 30-day treatment of CT26 tumor cells with 1µM compound **21** and compared with CT26 ERAAP knock-out cells (**Figure 5**). The unbound concentration was estimated to be 610nM in the study based on the fraction unbound measured in cell assay media (**Table 6**). Substantial evidence of peptide lengthening was observed; for example, significant increases in 10-mer and 11-mer peptides were accompanied by decreases in 9-mer and 8-mer peptides. Furthermore, the 10-mer and 11-mer peptides were typically extensions of 9-mer peptides (blue circles) indicating a clear effect of ERAAP inhibition or knock-out. The number of peptides lengthened by inhibitor treatment is apparently 80-90% of the total affected by knock-out, indicating a substantial pharmacological effect at an unbound concentration of 610nM, 23 and 100-fold higher than the biochemical inhibition measured using YTAFTIPSI and MART substrates respectively. **Figure 6** compares the magnitude of effect for compound **21** (y-axis) versus knock-out (x-axis) and the variance of the correlation line (blue) from the line of unity implies substantial but incomplete inhibition compared with the knock-out condition. Taken together these data provide substantial evidence that compound **21** is likely to induce significant modulation of the immunopeptidome in the 270mg/kg BID dose group given the unbound concentrations of 200-2000nM achieved (see **Figure 4**).

**Figure 5.**
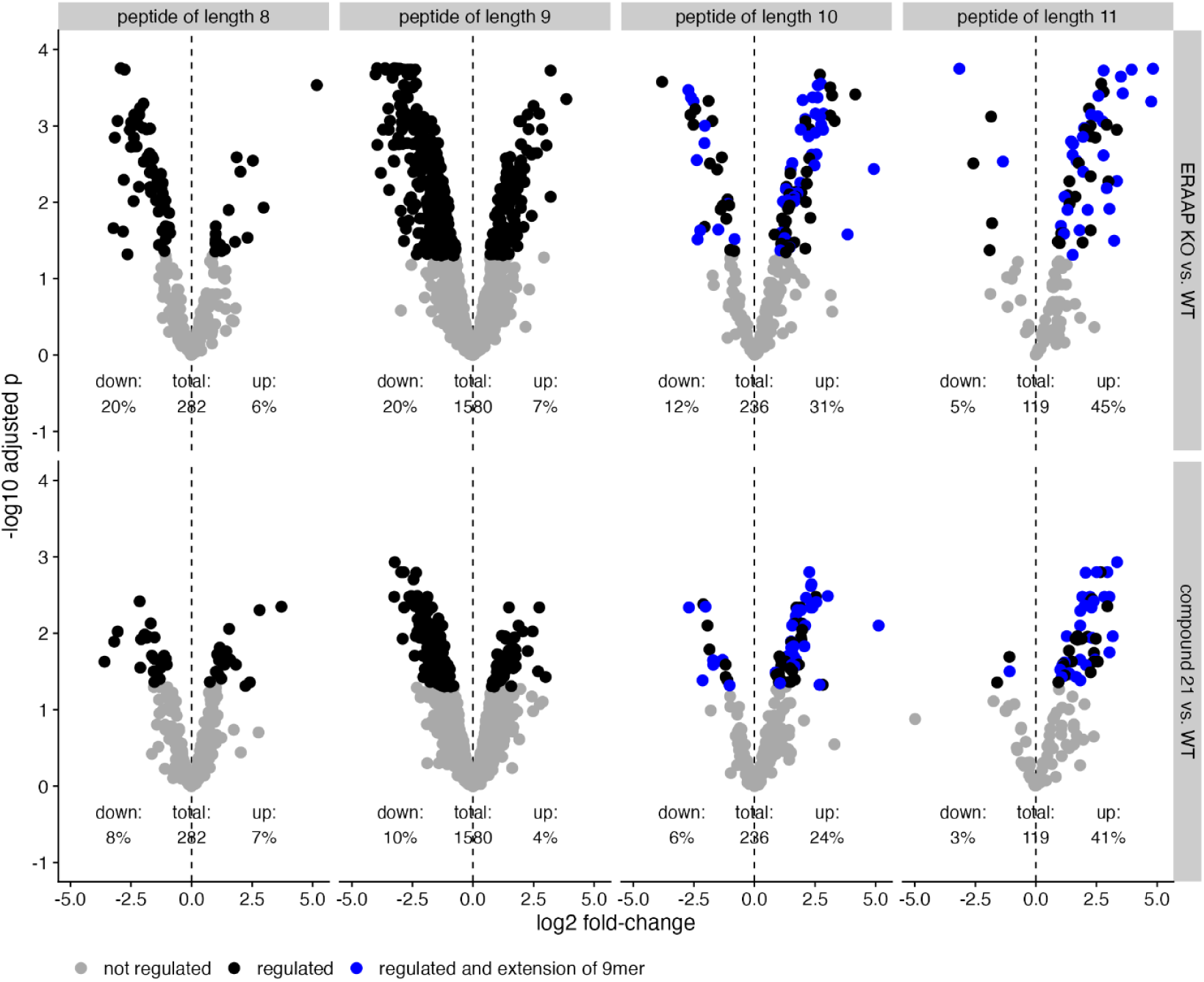
Impact on the CT26 tumor cell immunopeptidome *in vitro* by compound **21** (1µM total, 0.61µM unbound) and ERAAP knock-out compared to wild-type.

**Figure 6.**
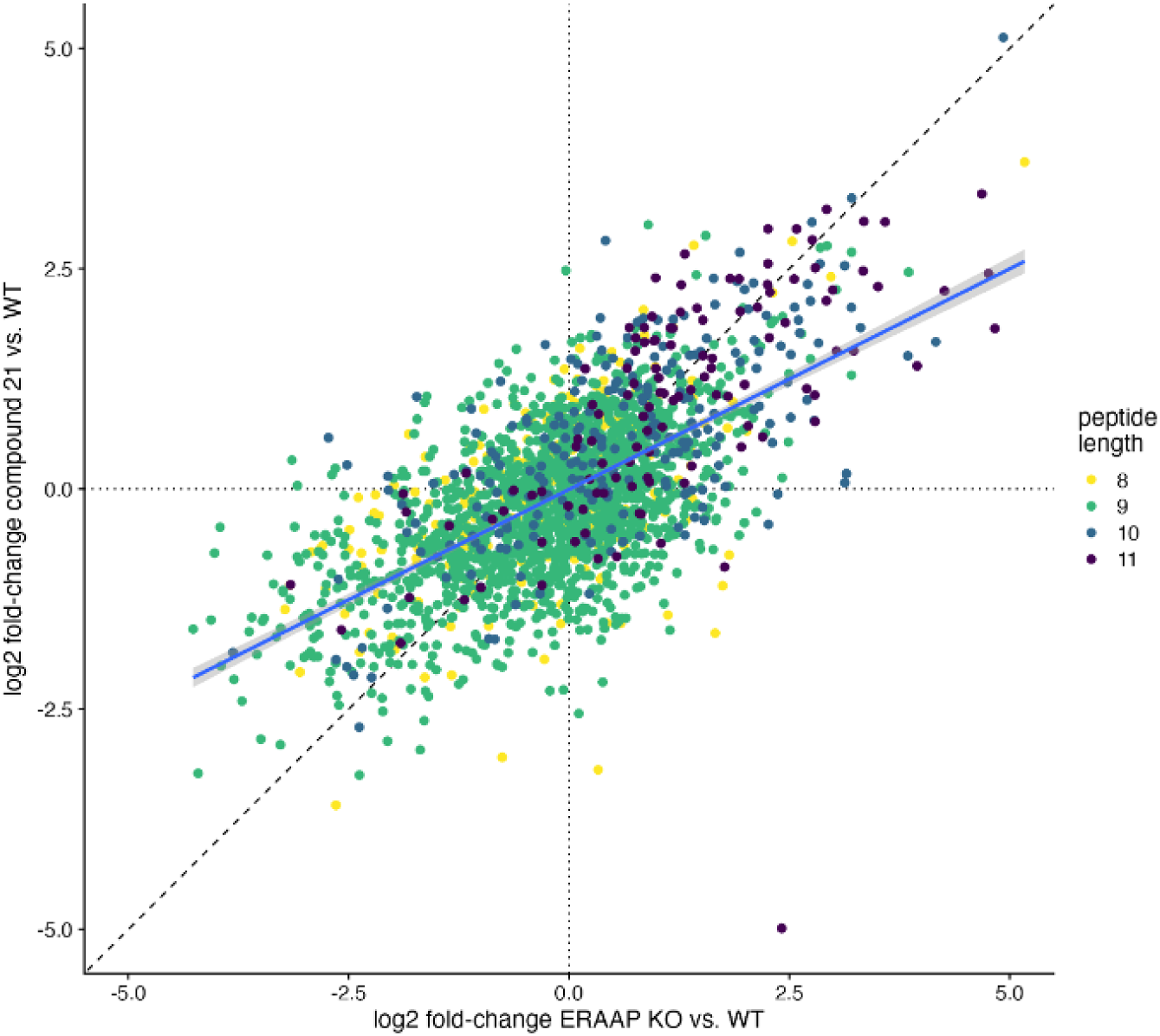
Comparison of compound **21** treatment and ERAAP knock-out on the immunopeptidome (CT26 cells)

### c) Impact of compound 21 on pathological endpoints *in vivo*

Tumor growth inhibition was evaluated in mice using the CT26 syngeneic model. Dosing of compound **21** was initiated concurrently with CT26 tumor cell inoculation (day 1) and treatment continued at 30, 90 and 270mg/kg BID with periodic monitoring of tumor growth. A clear dose response (270 > 90 > 30mg/kg) was observed with significant reduction in tumor volume (<50 mm^3^) by termination of the study, day 17, at the highest dose (**Figure 7**, pane A) and no significant dose-dependent body-weight changes were observed (**Figure 7**, pane B). The findings are not considered directly clinically translatable given the prophylactic nature of the experiment; however, the result motivated further tumor-bearing studies, the results of which will be reported in due course. Nevertheless, the data provide evidence for biologically meaningful modulation of tumor cell immunogenicity by compound **21**. Furthermore, the impact of 610nM compound **21** (unbound) on the immunopeptidome *in vitro* (*vide supra*) provides robust mechanistic support given that the unbound concentration in the 270mg/kg group was expected to be 200-2000nM.

**Figure 7.**
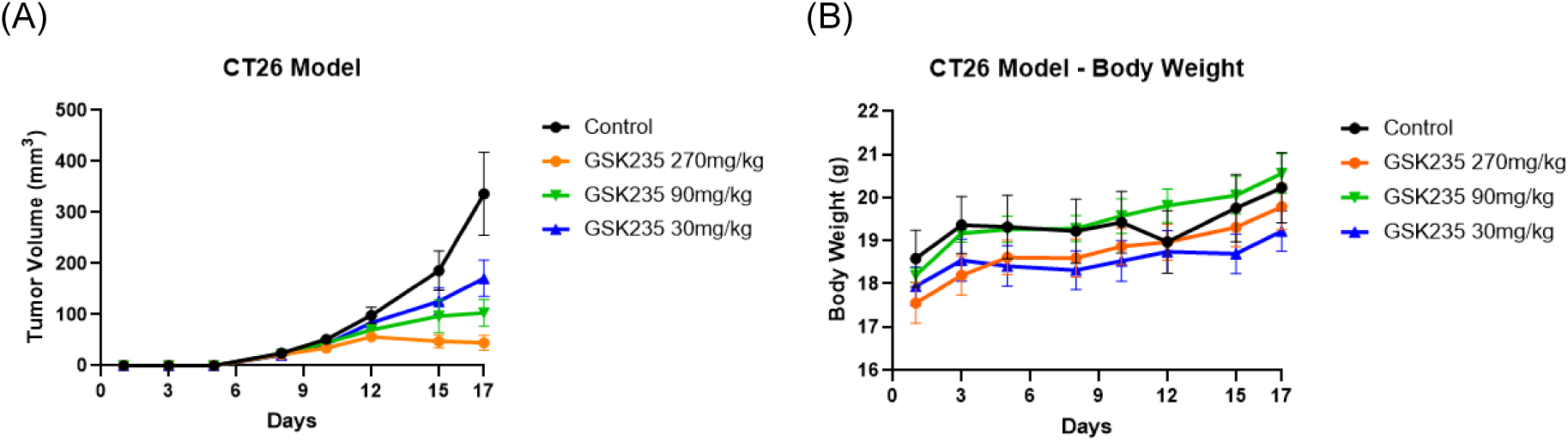
Tumor growth (A) and body weight change (B) with escalating oral doses of **21.**

**Figure 8.**
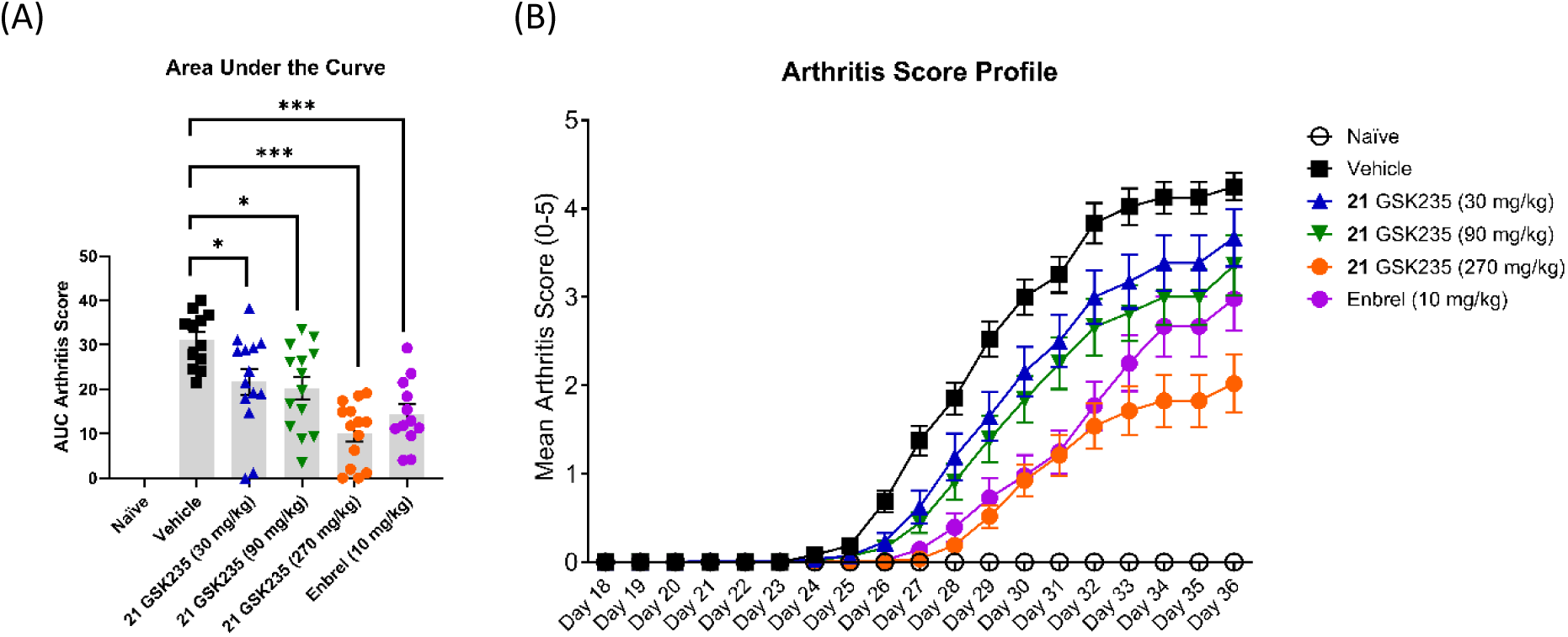
(A) AUC Arthritis Score and (B) Arthritis Score Profile comparing naïve and positive control (Enbrel) compared to escalating dose of compound **21**

Having established that compound **21** could drive a substantial response in tumor-bearing mice, we sought to explore potential risks by investigating the effects of ERAP1 inhibition in an inflammatory context. The pathology results from the CIA study are presented in **Figure 9**, with compound **21** eliciting a dose-proportional reduction of arthritis score that was at least as strong as the positive control, the TNF inhibitor Enbrel, used clinically to treat rheumatoid arthritis, in the 270mg/kg group. Minimal changes in body weight were observed compared to vehicle or positive control at the lowest and highest doses respectively (**Supplementary Figure 1**) and the findings clearly contradicted the hypothesis that MHC-presented peptide modulation could exacerbate an autoimmune response in an inflammatory setting.

**Figure 9.**
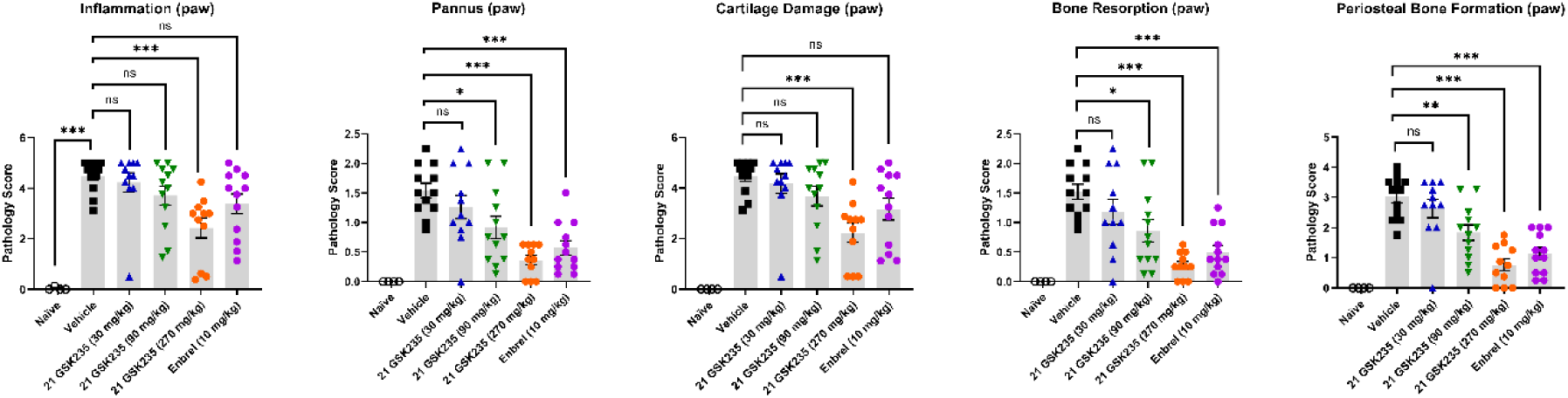
Paw histopathology with escalating dose of compound **21**

Multiple measures of paw pathology, including inflammation, pannus formation, cartilage damage, bone resorption and periosteal bone formation (**Figure 9**) closely reflected the arthritis score and confirmed a dose-dependent beneficial effect of ERAAP inhibition, again achieving comparable protection to the positive control at the higher dose. Knee pathology also reflected these findings (**Supplementary Figure 2**).

### d) In vivo pharmacology measurements

A suite of immunological endpoints was measured in the CIA study and indicated that the beneficial effects of compound **21** on arthritis readouts were linked to reductions in autoimmunity and inflammation. This included lower levels of anti-type II collagen antibodies in **21**-treated mice (**Figure 10**, pane A) as well as dose-dependent inhibition of IL-12p40 and IL-6 (**Figure 10**, pane B) whereas several other cytokines including KC, MCP-1 and TNF were unaffected by compound **21**. Notably, dose-dependent reductions in the cell surface expression of MHC-I protein were observed on B cell and conventional dendritic cell (cDC) populations in lymph nodes and spleen (**Figure 10**, pane C), consistent with modulation of ERAAP function by treatment with compound **21**.

**Figure 10.**
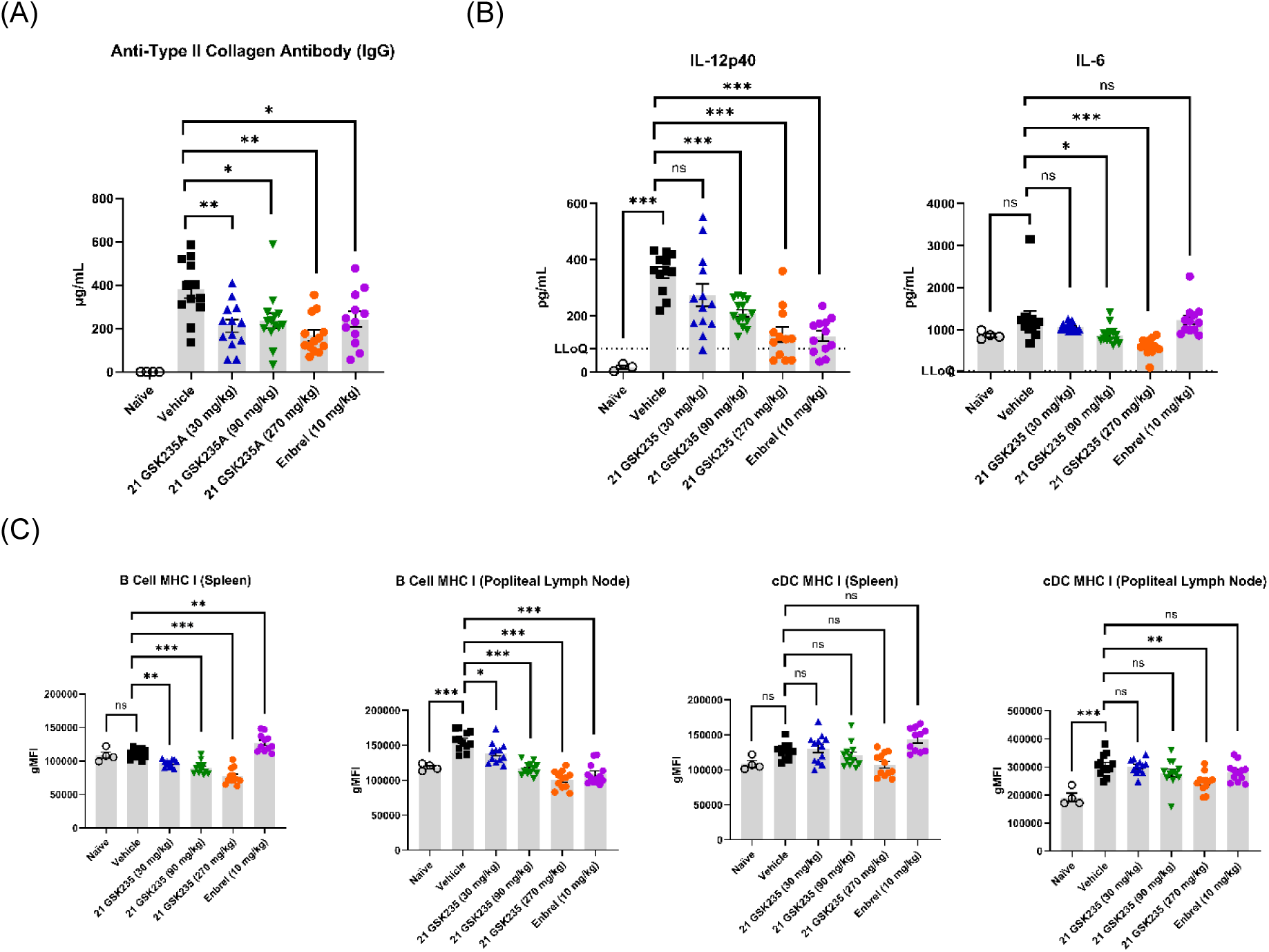
(A) Collagen antibody levels (B) IL-12p40 and IL-6 levels (C) MHC levels in B cells and cDC cells resulting from dose-escalation of compound **21**.

In addition to assessing CIA-specific pathology, broader tissue histopathology was evaluated to determine whether ERAAP inhibition might lead to non-specific inflammation and to investigate potential additional compound-related effects such as direct toxicity, metabolites or engagement of additional pharmacology. No significant effects attributable to the compound were noted in any of the tissues assessed, including numerous endpoints in the skin, muscle, liver inflammation or cellular infiltrate, gall bladder and small intestine.

In summary, the integrated data provides robust correlation of unbound drug levels with pharmacology across an oral dose range of 30mg/kg to 270mg/kg. Unbound drug levels of 200-2000nM were maintained for several weeks at 270mg/kg and clear linkage between proximal pharmacodynamic effects and downstream pharmacology is evident in both systems tested (1) *In vitro* treatment of CT26 cells with 610nM unbound concentration of compound **21** resulted in significant impact on the immunopeptide repertoire and CT26 tumor-growth regression was observed *in vivo* at the 270mg/kg dose, expected to achieve or exceed this concentration (2) Dose-dependent down-regulation of MHC proteins translated to significant impact on immunological and pathological end-points in the CIA study across the same dose range, achieving effects comparable to the positive control at 270mg/kg. Undesirable effects were not observed in either study, yet all available data indicates that substantial, chronic inhibition of ERAAP was achieved in both studies.

## Materials and Methods

### Synthetic Procedures

Chemical synthesis and characterisation are provided in **Supplementary Information**

### Crystallography Methods

The ERAP1 (1-941) Δ486-513 GSG insert, N70Q, N154Q, N414Q, N760Q protein was generated as described previously^29^ with a final protein buffer 50 mM Hepes pH 7, 150 mM NaCl. Crystallisation was carried out using sitting-drop vapour diffusion at 20°C, with 100nL + 100nL and 100nL + 50nL (protein + well) drops. Co-crystallisations were carried out for each ligand using 100 mM DMSO stock solutions with final ligand concentrations of ∼2 mM, using protein at ∼8 mg/ml. The crystallisation conditions were initially identified from PACT screen (Qiagen) and optimised in each case, with the conditions used to grow the harvested crystals as follows. Compounds **1** and **2**: 25% w/v PEG1500, 0.1M SPG buffer pH=5.5. Compound **8**: 25% w/v PEG1500, 0.1M SPG buffer pH=5.0. Compound **21**: 20% w/v PEG1500, 0.1M SPG buffer pH=5.5. Crystals were harvested into a 20% ethylene glycol cryoprotectant for a few seconds and then flash cooled in liquid nitrogen.

X-ray diffraction data was collected at 100K at Diamond Light Source beamline I03 (Compounds **1** and **2**), APS beamline 22-ID (Compound **8**) and ESRF beamline ID30B (Compound **21**). The datasets were processed and scaled using AUTOPROC (Compounds **1** and **21**),^43^ DIALS (Compound **2**),^44^ or HKL2000 (Compound **8**),^45^ utilising XDS,^46^ AIMLESS^47^ and the CCP4 suite of programs.^48^ The structures were determined using the coordinates of an isomorphous unliganded model of ERAP1 (unpublished). Preliminary refinement was carried out using BUSTER.^49^ The primary ligands were clearly visible in the resulting Fo-Fc electron density maps (**Supplementary Figure 3**). Model building was carried out with COOT,^50^ using a ligand dictionary generated from GRADE.^51^ Final refinement was carried out using BUSTER. Data collection statistics and refinement details for the final models are given in **Supplementary Table 2**. The coordinates and structure factors have been deposited in the Protein Data Bank under the accession codes XXX, XXX, XXX, and XXX.

### Biochemical assays

Biochemical activity monitoring the cleavage of 9-mer antigenic peptide YTAFTIPSI to 8-mer TAFTIPSI was performed as described previously,^31^ all data n≥3 unless otherwise specified. An identical method was used to determine ERAAP biochemical activity with YFATIPSI and EAAGIGILTV (MART1) peptide substrates. ERAP2 and IRAP biochemical assays were conducted as previously reported.^31^

### Cellular antigen presentation assay

The cellular antigen presentation assay data described in Tables 1-4 for compounds **1**-**21** was performed as described previously,^31^ all data n≥3 unless otherwise specified. Adjustments were made to the data normalisation method for data presented for compound **21** in Table 6, as described previously.^38^

### Physicochemical property data

Data was acquired using published protocols for ChromLogD_7.4,_^52^ kinetic solubility measured by CAD^53^ and FaSSIF^54^ solubility.

### Fraction Unbound in Blood, Microsomes, and Cellular Assay Media

Blood was obtained on the day of experimentation from in house GSK stock animals. Human liver microsomes were obtained from BioIVT (Westbury, NY) and diluted to 0.5mg protein/mL in potassium phosphate buffer. Cellular assay media was provided by GSK Biology. The fraction unbound in blood, human liver microsomes, and cellular assay media (DMEM/F-12 media with 10% FBS, 2 mM Glutamax, and 50 U/mL Pen-Strep) was determined using rapid equilibrium dialysis technology (RED plate (Linden Bioscience, Woburn, MA) at nominal concentrations of 0.5 to 5 µM. Matrices were dialyzed against phosphate buffered saline solution by incubating the dialysis units at 37°C for 4 h with constant shaking at 200 rpm. Following incubation aliquots of blood and buffer were matrix matched prior to analysis by LC−MS/MS. The unbound fraction was determined using the peak area ratios in buffer and in blood.

### Immunopeptidomics

#### i) MHC Class I Peptide Enrichment

Immunoprecipitation (IP) of MHC-I complexes was performed from CT26 cell lysates using a mouse monoclonal antibody (US Biological, M3885-73R) specific for H2-Kd/Dd. The protocol was adapted from Bassani Sternberg,^55^ with modifications to accommodate experimental conditions. Three biological replicates were processed per condition: ERAAP knock-out, compound **21** treatment, and vehicle control.

All steps involving antibody binding and elution were conducted at 4 °C to preserve complex integrity. CT26 cells were lysed according to the referenced protocol, and lysates were applied to affinity columns containing 2 mg of antibody per 400 µL bead matrix. Sequential washes were performed using buffers of increasing ionic strength (150 mM and 400 mM NaCl in 20 mM Tris-HCl, pH 8.0). Bound complexes were eluted with 0.1 M acetic acid. The presence of MHC complexes was confirmed by SDS-PAGE and silver staining. Elution fractions were pooled, dried via SpeedVac, and subjected to C18-based peptide cleanup prior to mass spectrometry analysis.

#### ii) Mass Spectrometry Analysis (MHC-I Peptide Enrichment)

Lyophilized peptide samples were resuspended in 0.05% trifluoroacetic acid and analysed using an Ultimate3000 nanoRLSC system (Dionex) coupled to an Orbitrap Exploris 480 mass spectrometer (Thermo Fisher Scientific). Peptides were separated on custom-packed 50 cm × 100 µm ID reversed-phase columns (C18, 1.9 µm, Reprosil-Pur, Dr. Maisch) maintained at 55 °C. Gradient elution was performed from 2% to 40% acetonitrile in 0.1% formic acid and 3.5% DMSO over 120 minutes.^56^

Raw data were processed using FragPipe v22.0. MSFragger^57^ was configured with the default “Nonspecific-HLA” workflow. Spectra were searched against a SwissProt-based Mus musculus protein database (downloaded January 11, 2018) supplemented with a custom contaminant list. Search parameters included: Precursor and fragment mass tolerance: ±20 ppm, Isotopic error: 0/1, Cleavage specificity: nonspecific, Peptide length: 7–25 amino acids, Variable modifications (max 3): Oxidation (M), Acetylation (Protein *N*-term), Pyro-glutamate formation from Q (nQ) and E (nE), Peptide-spectrum matches (PSMs) were re-scored using MSBooster with DIA-NN (v1.8.1) deep learning-based fragment.^58^ Validation was performed using Percolator.

Quantification was conducted using IonQuant with MaxLFQ-based label-free quantification, including match-between-runs (MBR).^59^ Ion-level false discovery rate (FDR) was controlled at 1%, followed by intensity-based normalization.

#### iii) Statistical Analysis

Statistical analysis was performed in R (http://www.r-project.org) using peptide intensities as a proxy for relative abundance. Variance stabilization normalization (VSN) was applied to derive log2-transformed intensities suitable for differential analysis. Only peptides quantified across all conditions were considered for further analysis. Differential abundance was assessed using the LIMMA modified t-test.^60^ P-values were adjusted for multiple testing using the Benjamini-Hochberg method. Peptides were considered significantly regulated if they exhibited an adjusted p-value < 0.05 and a fold-change ≥ 1.5.

### In Vivo Studies

All animal studies were ethically reviewed and carried out in accordance with Animals Scientific Procedures Act 1986 and the GSK Policy on the Care, Welfare and Treatment of Animals.

#### i) Mouse oral PK

Female BALB/c mice were orally dosed with compound at a target dose of 10mg/kg, formulated as suspensions in 1% aq. methylcellulose (w/v) at 1 mg/mL. Serial blood samples were collected up to 24h post oral administration. Following collection, aliquots of blood were diluted with an equal volume of water, mixed, prior to freezing on cardice and storage at -80°C until bioanalysis. Aliquots of diluted whole blood (50:50) were analysed using quantitative high-performance liquid chromatography with tandem mass spectrometric detection (LC-MS/MS) following protein precipitation. PK parameters were estimated from the blood concentration–time profiles using noncompartmental analysis with WinNonlin (Pharsight, Mountain View, CA).

#### ii) Mouse CT26 Tumor Model

Female BALB/c mice (6-8 weeks old) were acquired from Envigo and housed under standard laboratory conditions with ad libitum access to food and water. On study day one, animals were randomized by body weight into study groups and inoculated subcutaneously in the right hind flank with 5×10^4 CT26 cells. Concurrently, treatment administration began with either compound **21** (GSK235) or vehicle control and dosing was performed twice daily (BID) for the duration of the study.

#### iii) Mouse Collagen-Induced Arthritis (CIA) Model

A total of 70 male DBA/1OlaHsd mice (6–7 weeks old) were obtained from Envigo. Animals were housed under standard conditions with food and water provided ad libitum. On Day 0 (D0), mice received an intradermal (i.d.) injection of 0.2 mg bovine collagen type II emulsified in 100 µL Freund’s Complete Adjuvant (FCA). From Day 18 to Day 36, mice were administered either vehicle (1% methyl cellulose), compound **21** (GSK235) at doses of 30, 90, or 270mg/kg, or the positive control Enbrel at 10mg/kg. Vehicle and compound **21** (GSK235) were administered twice daily via oral gavage, while Enbrel was administered intraperitoneally every other day. On Day 21, a booster i.d. injection of 0.2 mg bovine collagen type II in 100 µL FCA was given.

Clinical arthritis scores were assessed in a blinded manner for all four paws (**Supplementary Table 3**). PK blood samples were collected on Days 18, 19, 27, 35, and 36. On Day 36, animals were euthanised, and serum was collected for quantification of anti-collagen type II IgG and pro-inflammatory cytokines. Tissues including joints, liver, small intestine, and skin were harvested for histological analysis. Spleen and lymph nodes (popliteal and inguinal) were collected for flow cytometry.

### Anti-Collagen Type II IgG Quantification

Serum samples were analysed in duplicate using a Mouse Anti-Collagen Type II IgG ELISA kit (Chondrex, Inc., Cat #2036T), following the manufacturer’s instructions.

### Multiplex Cytokine Assays

Levels of IL-12p40 and IL-6 were quantified using the U-PLEX multiplex platform (Meso Scale Discovery), according to the manufacturer’s protocol.

### Flow Cytometry

Single cell suspensions from spleen and pooled lymph nodes were stained using antibody panels detailed in **Supplementary Table 4**. Samples were processed with the True-Nuclear™ Transcription Factor Buffer Set (BioLegend, Cat #424401) and analysed on a CytoFLEX Flow Cytometry Analyzer (Beckman Coulter, Inc.).

## Discussion

Accumulating evidence supports the tractability of inhibiting ERAP1 for regulating human adaptive immune responses. Although ERAP1 has been reported to participate in several biological functions, the most established one is a highly specialized role in regulating which antigenic peptides can be presented by MHC-I molecules.^61^ A secondary and possibly related role in specific cellular and inflammatory context includes an autoinflammatory function when secreted and operates in the interface of innate and adaptive immunity.^10^ Both functions have been shown to be regulated by small molecule inhibitors and may synergize in regulating immune responses.^34,62,63^ As a result, down-regulation of ERAP1 activity may induce synergistic pleiotropic effects on T cell and NK cell responses as well as inflammatory signals. Such effects show promise in reprogramming the immune system to fight tumors since the upregulation of ERAP1 may act as an immune evasion mechanism. In addition, since antigen presentation is central to tumor responses, ERAP1 mediated immune effects may synergize with other immune modulating approaches in cancer immunotherapy efforts including immune checkpoint blockade, cancer vaccines, adoptive cell therapies, oncolytic viruses, epigenetic modulators, proteasome inhibitors, radiotherapy, NK cell therapies, Stimulator of interferon genes (STING) agonists, and cytokine therapies.^12,15,64–68^

Despite significant effort towards the development of ERAP1 inhibitors, suitable *in vivo* tools have been lacking and reported inhibitors suffer from limitations such as low potency, limited selectivity or low cellular potency.^24,25^ To address these issues, we pursued a library hit that targets the unique regulatory site of the enzyme, which was optimised to an *in vivo* tool ERAP1 inhibitor, compound **21** (GSK235). This compound has excellent potency and selectivity both *in vitro* and in cells, with favourable oral PK and target engagement profile. Furthermore, it induces relevant *in vivo* effects in two murine models suggesting that it is suitable as a tool for further *in vivo* investigations. The mechanism of action of compound **21** focuses on the regulatory site of ERAP1 that normally accommodates the *C*-termini of peptidic substrates and self-activates the enzyme so that it can preferentially cleave elongated peptides that can concurrently occupy both the active and regulatory site.^33^ Thus, although this compound acts as a non-competitive activator of small substrates, it is a competitive inhibitor of longer peptides.^31^ Small molecules targeting this site have been proposed to induce a large conformational change in ERAP1, part of its normal catalytic cycle, that closes of external access to the active site and blocks further catalytic site.^69^ Thus, although the allosteric site is mechanistically distinct, the effect on binding of small molecules can be functionally equivalent to active site blocking. This is supported by both cell-based assays for a single antigen and immunopeptidomics analysis that demonstrates functional equivalence of the reported compound to genetic knock-out. Thus, it appears that GSK235 can combine high selectivity by targeting a non-conserved site while retaining the pan-substrate effect of an active site inhibitor. This comes as a contrast to a previously described natural-product that targets the same site, which presented distinct effects on the cellular immunopeptidome.^35^ However, the potency of that compound was approximately 3 orders of magnitude weaker and intra-molecular competition between the *C*-termini of substrate peptides and the weak inhibition may underlie such effects.

Testing the efficacy of GSK235 in a murine cancer model demonstrated effective and dose-dependent retardation of tumor growth, leading to tumor control at the higher dosages. The pharmacodynamic effect corresponds well to immunopeptidome shifts identified *in cellulo*, consistent with the main therapeutic hypothesis that ERAP1 inhibition leads to immunopeptidome shifts that induce more potent or novel cellular adaptive immune responses. Although we did not proceed to detailed immunological analysis using this tool, observed effects parallel the described efficacy of ERAP1 genetic knockdown in other murine models that underlie T cell and NK cell mechanisms.^11–13,15,17^ However, the interpretation of immune effects after genetic downregulation of ERAP1 may have important limitations for translating to therapeutic interventions due to the dynamic and spatial nature of immune responses. Specifically, murine knock-out models may adapt to lack of ERAP1 activity pre-tolerance and genetic down-regulation in implanted cell lines simulates perfect cellular targeting that is not easily achievable clinically. Thus, GSK235 constitutes a valuable tool for interrogating the *in vivo* role of ERAP1 in different cancer immunotherapy settings and more importantly in combinatorial treatments. Combinatorial cancer immunotherapy is actively being pursued due to its promise in avoiding cancer evasion mechanisms.^70^ Although GSK235 was found to be surprisingly effective as a monotherapy in our murine model, future investigations will require investigation into the interplay of ERAP1 inhibition with other immune system interventions in cancer.

The significant effects of ERAP1 inhibition on the cellular immunopeptidome have raised concerns for potential autoimmune effects due to the presentation of self-antigens that are normally repressed.^71^ Self-mimicry due to presentation of self-antigenic peptides has been proposed to be a major mechanism behind MHC-I-opathies a class of inflammatory diseases of autoimmune aetiology that correlate with specific HLA alleles and ERAP1 allotypes, often in epistasis.^19,72–74^ In addition, potential effects of ERAP1 on ER homeostasis, stress and the unfolded protein responses have been proposed as separate or complementary mechanisms.^75,76^ These mechanisms generate concerns of immune toxicity of ERAP1 inhibitors that could complicate therapeutic approaches. On the other hand, enhanced ERAP1 activity has been correlated mechanistically with predisposition to these diseases, suggesting that ERAP1 inhibition may hold therapeutic promise.^77^

To gain further insight into these concerns, we evaluated the *in vivo* effects of GSK235 in the mouse collagen induced arthritis model. ERAP1 inhibition did not exacerbate disease but had a strong, dose-dependent, therapeutic effect that equalled or surpassed the effect of Enbrel, a TNF inhibitor used clinically to treat rheumatoid arthritis. These effects closely correlated with reductions in anti-collagen IgG and B cell MHC-I levels as well as with IL-12p40 and IL-6 production. These results add to previous reports indicating that ERAP1 knockdown or inhibition can dampen T cell and NK cell activation and cytokine signalling, while also having effects on dendritic cell and macrophage function, limiting Th1 and Th17 polarization.^63,78^ The effects observed here on autoantibody production could be mediated by down-regulated generation of unknown self-antigenic epitopes and subsequent reduced antigen presentation to T helper cells, either during initial T cell priming on dendritic cells or in the context of T cell – B cell help. Since the T cell – antigen-presenting cell interactions driving antibody responses typically involve MHC class II peptide presentation, such a mechanism would need to invoke a (direct or indirect) role for ERAP1 in the MHC II pathway; a secondary role in shaping the peptide load of MHC class II molecules has been previously proposed.^79^ Conversely, reduced T and B cell responses may be secondary to a reduced overall inflammatory environment mediated by ERAP1 inhibition. Thus, the effects of ERAP1 function on innate immune responses may also play a role and could include inhibition of secreted ERAP1 by B cells or macrophages or the effect of ERAP1 in cellular homeostasis leading to reduction of innate immunity responses.^10,75,80^ The molecular details of these complex and potentially synergistic effects will have to be thoroughly analysed in future studies and GSK235 will undoubtably be a valuable tool since it can perturb ERAP1 function dynamically, circumventing limitations of pre-tolerance genetic manipulations. Still, our results suggest that ERAP1 inhibition may have a significant therapeutic potential not only for HLA-associated autoimmunity but also in other forms of inflammatory autoimmunity that should be explored.^81^

## Conclusion

In summary, we report the discovery of the first potent *in vivo* tool for interrogating ERAP1 biology that also constitutes a potential lead molecule for drug-discovery efforts. Hit compound **1** was identified *via* high-throughput screening and early generation of X-ray crystallography allowed efficient optimisation to lead compound **21** (GSK235) through introduction of productive interactions. Compound **21** was progressed firstly to a pilot prophylactic study in tumor-bearing mice, confirming dose-dependent tumor growth inhibition, then to a comprehensive CIA model to assess risks implied by literature studies with knock-out mice. Administration of compound **21** across a substantial dose range resulted in significant protection in this model with integration of *in* vitro and *in* vivo data strongly implying 50-80% inhibition of ERAAP through the study. The CIA model findings do not directly support a protective role for ERAP1 inhibition in human disease; however, the data substantially allay concerns of disease exacerbation resulting from ERAP1 inhibition on an inflammatory background and, encouragingly, no substantial pathological consequences were evident in the study. Our results help establish the tractability of ERAP1 inhibition for cancer immunotherapy and expand options to certain forms of inflammatory autoimmune disease, not limited to MHC-I-opathies. The public availability of GSK235 will empower mechanistic studies that will help progress the therapeutic value of ERAP1 inhibition and facilitate the exploration of combinatorial strategies aiming to reprogram the immune system.

## Acknowledgements

The authors thank Heather Barnett, Stephen Besley, Shenaz Bunally, Esme Clarke, Yanan He, Justyna Korczynska, Stefan Maehringer, Ferdausi Mazumder, Richard Upton and Bob Watson.

## Additional Information

**Supplementary Information** contains supplementary figures & tables, synthetic procedures and X-ray crystallography methods

## Corresponding Authors

Correspondence to Simon Peace and Efstratios Stratikos

## Competing interests

All authors except D. Koumantou, I. Temponeras and E. Stratikos were employees at GSK at time of this work.

