## Supplementary Information for "Discovery of an orally available potent ER aminopeptidase 1 (ERAP1) inhibitor that enhances anti-tumor responses and limits inflammatory autoimmunity in vivo"

**Supplementary Figure 1.** Bodyweight profile from CIA model

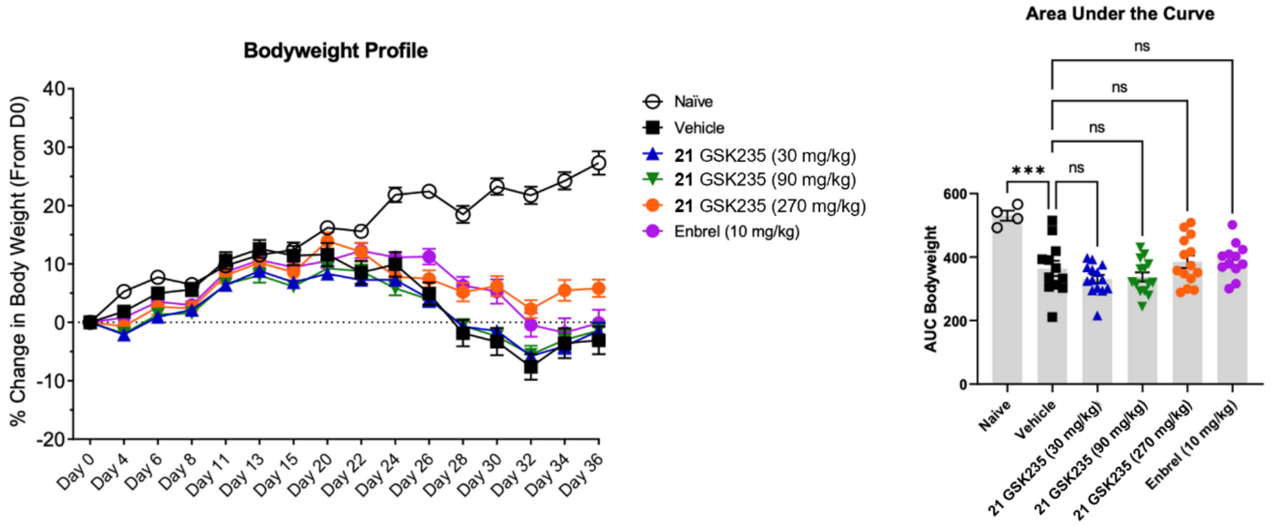

**Supplementary Figure 2.** Knee pathology from CIA model

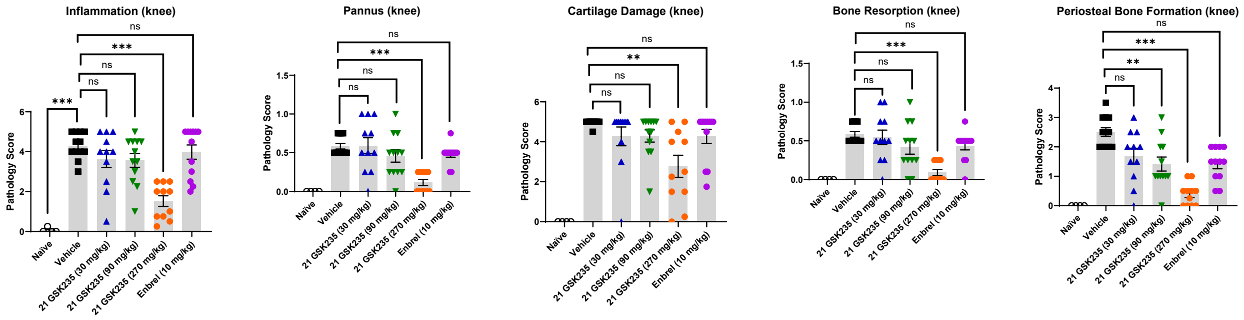

**Supplementary Figure 3.** ERAP1 electron density maps in the compound binding sites

Electron density $F_o$-$F_c$ difference maps contoured at 3σ (green) and -3σ (red) superposed on the refined protein-inhibitor complex structures. Maps calculated after refinement of the unliganded model.

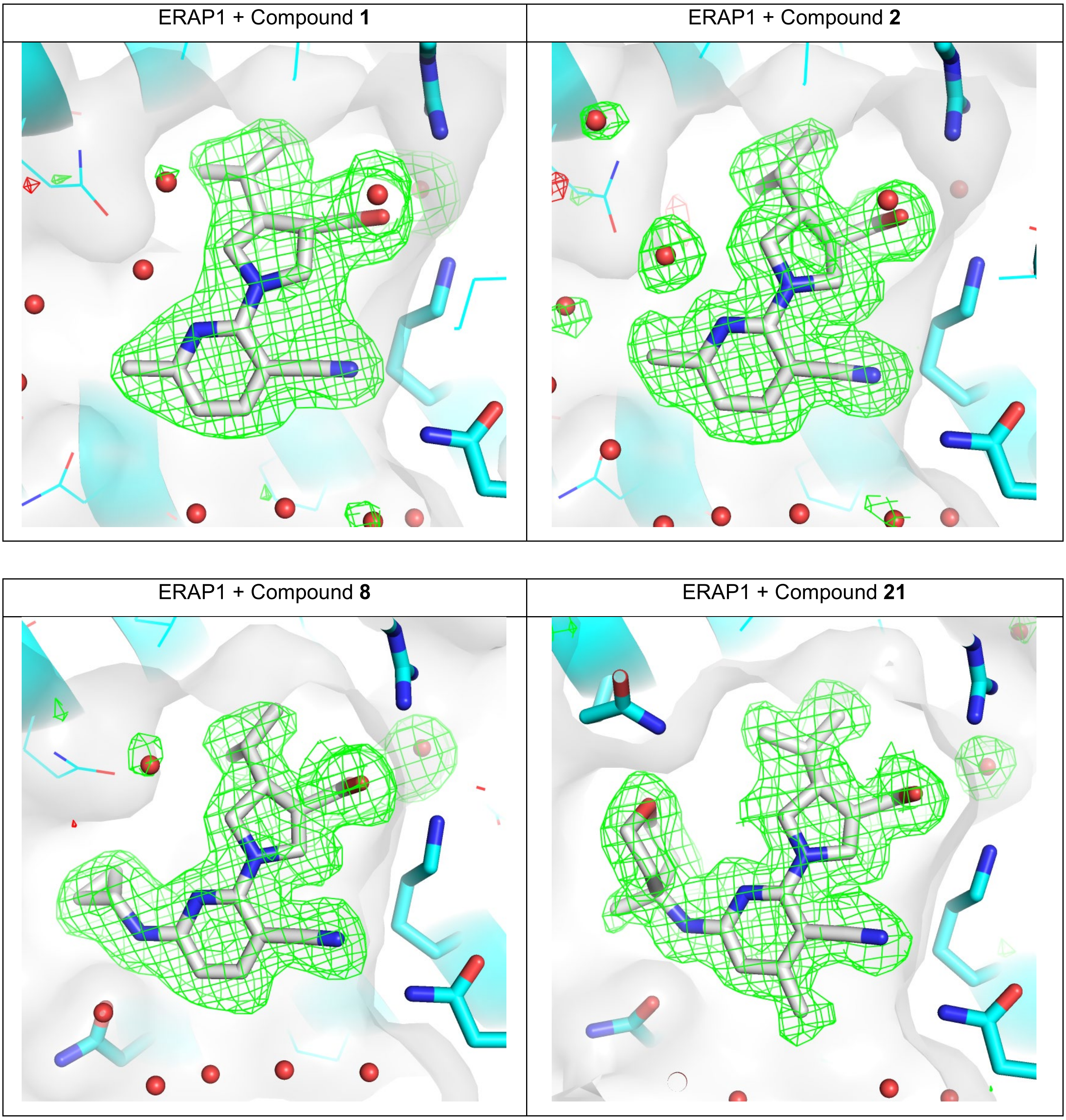

**Supplementary Table 1.** *In vitro* clearance from human hepatocytes

| Compound | Human hep $CL_{int}$ (mL/min/g tissue) | Human hep $CL_{int}$ (mL/min/kg, BWT) | $f_{u,mic}$ (human) | $f_{u,hep}$ (human)[1] | Human hep $CL_{int,u}$ (mL/min/kg) |
|---|---|---|---|---|---|
| **19** | 2.53 | 62.0 | 0.86 | 0.91 | 68 |
| **21** | 1.43 | 35.1 | 0.83 | 0.89 | 39 |

[1]Kilford equation used to predict fu,hep from fu,mic

**Supplementary Table 2.** ERAP1 data collection and refinement statistics

| | **Compound 1** | **Compound 2** | **Compound 8** | **Compound 21** |
|---|---|---|---|---|
| **Data collection** | | | | |
| Space group | $P2_12_12$ | $P2_12_12$ | $P2_12_12$ | $P2_12_12$ |
| Cell dimensions | | | | |
| *a*, *b*, *c* (Å) | 112.9, 140.3, 57.7 | 113.1, 140.4, 57.6 | 113.1, 140.7, 57.5 | 113.0, 140.6, 57.8 |
| $\alpha, \beta, \gamma$ (°) | 90, 90, 90 | 90, 90, 90 | 90, 90, 90 | 90, 90, 90 |
| Resolution (Å) | 88-1.87 (1.91-1.87) | 71-1.63 (1.67-1.63) | 32-1.73 (1.78-1.73) | 89-1.74 (1.80-1.74) |
| Observations | 406390 (23662) | 747410 (54634) | 438698 (-) | 352833 (19180) |
| Unique reflections | 76171 (3705) | 115183 (8412) | 95006 (4648) | 82810 (4141) |
| $R_{meas}$ | 0.242 (3.585) | 0.224 (3.228) | 0.113 (0.795) | 0.173 (1.148) |
| $CC_{1/2}$ | 0.995 (0.293) | 0.995 (0.320) | 0.993 (0.647) | 0.963 (0.540) |
| $I/\sigma I$ | 6.3 (0.5) | 6.8 (1.0) | 11.1 (1.1) | 6.3 (1.5) |
| Completeness | | | | |
| Spherical (%) | 100 (100) | 100 (100) | 99.0 (97.5) | 86.3 (45.1) |
| Ellipsoidal (%) | - | - | - | 89.3 (55.5) |
| Multiplicity | 6.6 (6.4) | 6.5 (6.5) | 4.6 (3.9) | 4.3 (4.6) |
| **Refinement** | | | | |
| Resolution (Å) | 71-1.87 (1.89-1.87) | 60-1.63 (1.64-1.63) | 32-1.73 (1.75-1.73) | 89-1.74 (1.77-1.74) |
| $R_{work}$ | 0.167 (0.282) | 0.159 (0.231) | 0.169 (0.247) | 0.208 (0.280) |
| $R_{free}$ | 0.211 (0.288) | 0.185 (0.240) | 0.203 (0.326) | 0.255 (0.310) |
| No. atoms | 7923 | 8273 | 8107 | 8254 |
| Protein | 6953 | 6994 | 6975 | 7049 |
| Ligands | 59 | 67 | 58 | 98 |
| Waters | 911 | 1212 | 1074 | 1107 |
| B-factors (Å$^2$) | | | | |
| Protein | 32.5 | 22.0 | 24.5 | 20.5 |
| Ligands | 40.4 | 28.3 | 30.7 | 26.9 |
| Waters | 45.8 | 41.0 | 39.8 | 35.2 |
| R.m.s deviations | | | | |
| Bond lengths (Å) | 0.010 | 0.010 | 0.010 | 0.008 |
| Bond angles (°) | 1.05 | 0.98 | 1.00 | 0.94 |

*Highest resolution shell shown in parentheses

**Supplementary Table 3.** Clinical scoring system

Clinical arthritis scores were recorded for all 4 paws daily from D18 through D36 using the following criteria:

| | |
|---|---|
| Grade 0 | Normal |
| Grade 1 | One hind or forepaw joint affected or minimal diffuse erythema and swelling |
| Grade 2 | Two hind or forepaw joints affected or mild diffuse erythema and swelling |
| Grade 3 | Three hind or forepaw joints affected or moderate diffuse erythema and swelling |
| Grade 4 | Four hind or forepaw joints affected or marked diffuse erythema and swelling |
| Grade 5 | Entire paw affected, severe diffuse erythema and severe swelling, unable to flex digits |

**Supplementary Table 4.** Antibody panels

T cell panel

| Fluorophore | Marker | Clone | Company | Cat # |
|---|---|---|---|---|
| FITC | CD3 | 145-2C11 | Tonbo Biosciences | 35-0031-U500 |
| BV786 | Ki-67 | B56 | BD | 563756 |
| PE | CD69 | H1.2F3 | BioLegend, Inc. | 104507 |
| PerCP/Cy5.5 | CD198/CCR8 | SA214G2 | BioLegend, Inc. | 150308 |
| PE-Cy7 | CD278 (ICOS) | C398.4A | BioLegend, Inc. | 313520 |
| Alexa 647 | CD25 | PC61 | BioLegend, Inc. | 102020 |
| BV605 | CD44 | IM7 | BioLegend, Inc. | 103047 |
| BV421 | FOXP3 | MF-14 | BioLegend, Inc. | 126419 |
| Red710 | CD4 | RM4-5 | Tonbo Biosciences | 80-0042-U100 |
| Red780 | Viability | N/A | Tonbo Biosciences | 13-0865-T500 |
| BV510 | CD8a | 53-6.7 | BD | 563068 |

*B Cell Panel*

| Fluorophore | Marker | Clone | Company | Cat # |
|---|---|---|---|---|
| FITC | Mouse IgG1 | A85-1 | BD | 562026 |
| FITC | Mouse IgG2a/b | R2-40 | BD | 553399 |
| BV786 | CD11b | MI/70 | BioLegend, Inc. | 101243 |
| PE | GL7 | GL7 | BioLegend, Inc. | 144608 |
| BV510 | CD138 | 281-2 | BioLegend, Inc. | 142521 |
| BB700 | IAIE | M5/114.15.2 | BD | 746197 |
| PE-Cy7 | Ki-67 | B56 | BD | 561283 |
| Alexa 647 | H-2Kq | H-2Kq | BioLegend, Inc. | 115106 |
| BV605 | CD19 | 6D5 | BioLegend, Inc. | 115540 |
| BV650 | CD3 | 17A2 | BD | 740530 |
| BV421 | Mouse IgD | 11-26c.2a | BioLegend, Inc. | 405725 |
| Alexa 700 | CD11c | N418 | BioLegend, Inc. | 117320 |
| Red780 | Viability | N/A | Tonbo Biosciences | 13-0865-T500 |

#### Statistical Analysis for CIA model

Statistical analysis was performed in R. A one-way ANOVA model was constructed with Treatment as a fixed effect. Contrast tests were carried out to test the difference in effects between the Vehicle/Naïve groups (untreated) and the different doses of compound **21** (GSK235) and Enbrel. P-values were adjusted for multiple testing using the Benjamini-Hochberg method and effects were considered significant if an adjusted p-value < 0.05 (*), <0.01 (**), <0.001 (***).

#### *In vitro* hepatocyte intrinsic clearance

Compounds were incubated at 0.5 µM in Williams Medium E with hepatocytes of the appropriate species at 0.5 million cells/mL. The incubation plate was kept in a 37°C incubator in the presence of 5% CO2 with constant shaking at 200 rpm. Aliquots were collected at time points between 0 and 240 minutes. The resultant samples were extracted by protein precipitation with acetonitrile containing an analytical internal standard. The samples were analysed by LC-MS/MS and elimination rate constant (k) for loss of compound from the incubation mixtures was calculated from the slope of the log-transformed analyte:internal standard peak area ratio versus time curve. Data was then scaled using in house scaling factors to mL/min/kg body weight.

#### Cross-screening panel

No activity of concern was observed against additional targets in the cross-screening liability panel beyond reported activity in **Table 6**. These assays are performed in various formats depending on target and risk directionality, and include:

5-HT1B Agonist (HTR1B) - pEC50
5-HT2A Agonist (HTR2A) - $pEC_{50}$
5-HT2A Antagonist (HTR2A) - $pIC_{50}$
5-HT2B Agonist (HTR2B) - $pEC_{50}$
5-HT2C Agonist (HTR2C) - $pEC_{50}$
5-HT2C Antagonist (HTR2C) - $pIC_{50}$
5-HT3 Agonist (HTR3A) - $pEC_{50}$
5-HT3 Antagonist (HTR3A) - $pIC_{50}$
nAChR - Agonist - $pEC_{50}$
nAChR - $pIC_{50}$
Acetylcholinesterase (AChE) - $pIC_{50}$
Adenosine A2a Human Agonist - $pEC_{50}$
Alpha 1 nAChR Human Agonist FLIPR - $pEC_{50}$
Alpha 1 nAChR Human Antagonist - $pIC_{50}$
Alpha 2C Adrenoceptor - $pEC_{50}$
Alpha1B (ADRA1B) Antagonist - $pIC_{50}$
AR Agonist - $pEC_{50}$
AR Antagonist - $pIC_{50}$
Aryl Hydrocarbon Receptor (AhR) - $pEC_{50}$
Aurora B Kinase - $pIC_{50}$
Beta2 Adrenoceptor Human Agonist $pEC_{50}$
Beta2 Adrenoceptor Human Antagonist $pIC_{50}$
Bile Salt Export Pump (BSEP) $pIC_{50}$
CB1 Agonist (CNR1) - $pEC_{50}$
Cyclooxygenase 2 (COX-2) Human FLINT - $pIC_{50}$
D1 Antagonist (DRD1) - $pIC_{50}$
Dopamine 1 Receptor (DRD1) - $pXC_{50}$
Dopamine D2 (D2L) Agonist - $pEC_{50}$
Dopamine D2 (D2L) Antagonist - $pIC_{50}$
ER Agonist (NHR) - $pEC_{50}$
ER Antagonist (NHR) - $pIC_{50}$
GABAA Human - Agonism - $pEC_{50}$
GABAA Human - Antagonism - $pIC_{50}$
GR Agonist (NHR) - $pEC_{50}$
hCav1.2 Blocker Electrophysiology Qube - $pIC_{50}$
hERG (human) Blocker Electrophysiology Qube - $pIC_{50}$
Histamine 1 (HRH1) Antagonist - $pIC_{50}$
Histamine 3 Receptor (HRH3) BRET - $pXC_{50}$
hKV1.5 Blocker Electrophysiology QUBE - $pIC_{50}$
hNav1.5 Blocker Electrophysiology Qube - $pIC_{50}$
KCNQ1 (human) Blocker Electrophysiology Qube - $pIC_{50}$
LCK - $pIC_{50}$
M1 Agonist (CHRM1) - $pEC_{50}$
M1 Antagonist (CHRM1) - $pIC_{50}$
M2 (CHRM2) Human Ag - $pEC_{50}$
M2 (CHRM2) Human Antag - $pIC_{50}$
MATE1 (Human) - $pIC_{50}$
Monoamine Oxidase A (MAOA) - $pIC_{50}$
Mu Opioid (OPRM1) Agonist - $pEC_{50}$
NET Blocker - Transporter Assay - $pIC_{50}$
NK1 Antagonist (TACR1) - $pIC_{50}$
NMDA 1A/2B (NMDAR) - $pIC_{50}$
NR1I2 Rat PXR - $pEC_{50}$
OATP1B1 - $pIC_{50}$
OCT2 (Human) - $pIC_{50}$
OPRK1 Agonist - $pEC_{50}$
Phosphodiesterase 3A (PDE3A) - $pIC_{50}$
Phosphodiesterase 4B (PDE4B) - $pIC_{50}$
PI3K-gamma Human Inhibition - $pIC_{50}$
PPARG Agonist (NHR) - $pEC_{50}$
PXR (NR1I2) Human Agonist - $pEC_{50}$
SERT Blocker - Transporter Assay - $pIC_{50}$
V1a Antagonist (AVPR1a) - $pIC_{50}$

#### General Chemistry Procedures

All compounds used for biological testing were >95% pure by HPLC, LCMS or NMR.

Unless otherwise stated, all reactions were carried using anhydrous solvents. Solvents and reagents were purchased from commercial suppliers and used as received. Reactions were monitored by thin layer chromatography (TLC) or LCMS. TLC was carried out on glass or aluminium-backed 60 silica plates coated with UV254 fluorescent indicator. Spots were visualised using UV light (254 or 365 nm) or common staining methods as appropriate. Silica flash chromatography was carried out using Teledyne Isco CombiFlash® apparatus using RediSep® pre-packed silica cartridges. Extracted organic mixtures were dried using Biotage PTFE hydrophobic phase separator frits unless otherwise stated. Microwave chemistry was performed in a Biotage Initator using initial high setting.

NMR spectra were recorded at rt (unless otherwise stated) using standard pulse methods on a Bruker AV-400 spectrometer ($^{1}H$ = 400 MHz, $^{13}C$ = 101 MHz). Chemical shifts are referenced to trimethylsilane (TMS) or the residual solvent peak, and are reported in ppm. Coupling constants are reported as observed in Hz and refer to $^{3}J_{H\text{-}H}$ couplings, unless otherwise stated. Coupling constants are quoted to the nearest 0.1 Hz and multiplicities are given the following abbreviations and combinations thereof: s (singlet), d (doublet), t (triplet), q (quartet), quin (quintet), sxt (sextet), m (multiplet), br. (broad). Pairs of coupling constants were averaged to the nearest 0.1 Hz.

Routine LCMS analysis was carried out on a Waters Acquity UPLC instrument equipped with a BEH or CSH column (50 mm x 2.1 mm, 1.7 µm packing diameter) and Waters micromass ZQ MS using alternate-scan positive and negative electrospray. Analytes were detected as a summed UV wavelength of 210 – 350 nm. Two liquid phase methods were used:

**Formic**: 40 °C, 1 mL/min flow rate. Gradient elution with the mobile phases as (A) water containing 0.1% volume/volume (v/v) formic acid and (B) acetonitrile containing 0.1% (v/v) formic acid. Gradient conditions were initially 1% B, increasing linearly to 97% B over 1.5 min, remaining at 97% B for 0.4 min then increasing to 100% B over 0.1 min.

**High pH**: 40 °C, 1 mL/min flow rate. Gradient elution with the mobile phases as (A) 10 mM aqueous ammonium bicarbonate solution, adjusted to pH 10 with 0.88 M aqueous ammonia and (B) acetonitrile. Gradient conditions were initially 1% B, increasing linearly to 97% B over 1.5 min, remaining at 97% B for 0.4 min then increasing to 100% B over 0.1 min.

Additional LCMS analysis was carried out on a Waters Acquity UPLC instrument equipped with a BEH C18 column (100 mm x 2.1 mm, 1.7 µm packing diameter) and Waters micromass ZQ MS using positive electrospray. Analytes were detected as a summed UV wavelength of 210 – 500 nm. The following liquid phase method was used:

**TFA**: 50 °C, 0.8 mL/min flow rate. Gradient elution with the mobile phases as (A) water containing 0.1% volume/volume (v/v) trifluoroacetic acid and (B) acetonitrile containing 0.1% (v/v) trifluoroacetic acid. Gradient conditions were initially 3% B, increasing linearly to 99.9% B over 8.5 min, remaining at 99.9% B for 0.5 min, then reducing to 97% B over 0.5 min then remaining at 97% B for 0.5 min.

**Mass directed automatic purification (MDAP)**

**Formic MDAP:** The HPLC separation was conducted on an Xselect CSH C18 column (150 mm x 30 mm i.d. 5 µm packing diameter) at ambient temperature, eluting with 0.1% formic acid in water (solvent A) and 0.1% formic acid in acetonitrile (solvent B) using an elution gradient of between 0 and 100% solvent B over 15 or 25 min. The UV detection was an averaged signal from wavelength of 210 nm to 350 nm. The mass spectra were recorded on a Waters ZQ Mass Spectrometer using alternate-scan positive and negative electrospray.

**High pH MDAP:** The HPLC analysis was conducted on an Xselect CSH C18 column (150 mm x 30 mm i.d. 5 µm packing diameter) at ambient temperature, eluting with 10 mM ammonium bicarbonate in water adjusted to pH 10 with ammonia solution (solvent A) and acetonitrile (solvent B) using an elution gradient of between 0 and 100% solvent B over 15 or 25 min. The UV detection was an averaged signal from wavelength of 210 nm to 350 nm. The mass spectra were recorded on a Waters ZQ Mass Spectrometer using alternate-scan positive and negative electrospray.

**Synthetic Procedures**

**Compounds 1 and 2**

**Methyl *trans*-1-(3-cyano-6-methylpyridin-2-yl)-4-isopropylpyrrolidine-3-carboxylate (22)**

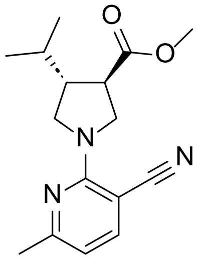

2-Chloro-6-methylnicotinonitrile (50.0 mg, 0.328 mmol) was added to a solution of *trans*-methyl 4-isopropylpyrrolidine-3-carboxylate hydrochloride (82.0 mg, 0.395 mmol) and DIPEA (0.200 mL, 1.15 mmol) in DMSO (0.8 mL). The mixture was heated in a microwave at 120 °C for 1.5 h. The reaction mixture was directly purified by MDAP (high pH), then the appropriate fractions were combined and concentrated under a stream of nitrogen to give the required product (60 mg, 64% yield) as an off-white solid.

**$^1$H NMR** (Methanol-d$_4$, 400 MHz) δ 7.68 (d, 1H, *J* = 7.9 Hz), 6.57 (d, 1H, *J* = 7.9 Hz), 4.08 (dd, 1H, *J* = 8.4, 10.8 Hz), 3.99 (dd, 1H, *J* = 7.9, 10.8 Hz), 3.84 (dd, 1H, *J* = 8.9, 10.8 Hz), 3.74 (s, 3H), 3.58 (dd, 1H, *J* = 8.9, 10.8 Hz), 3.0-3.1 (m, 1H), 2.4-2.5 (m, 1H), 2.39 (s, 3H), 1.78 (sxtd, 1H, *J* = 6.8, 13.8 Hz), 0.99 (m, 6H). **LCMS** (HpH, 2 min): Rt = 1.31 mins, MH+ = 288 (100% a/a).

**Methyl (3*S*,4*S*)-1-(3-cyano-6-methylpyridin-2-yl)-4-isopropylpyrrolidine-3-carboxylate (23) and methyl (3*R*,4*R*)-1-(3-cyano-6-methylpyridin-2-yl)-4-isopropylpyrrolidine-3-carboxylate (24)**

Methyl *trans*-1-(3-cyano-6-methylpyridin-2-yl)-4-isopropylpyrrolidine-3-carboxylate (60 mg) was purified by chiral HPLC using the following method:

**Sample preparation:** Total sample dissolved in ca. 10ml 50:50 Heptane/ Ethanol
**Column:** Chiralpak IF (250mmx30mm, 5micron, ambient temp)
**Temperature:** Ambient
**Flow Rate:** 42.5 mL/min
**Injection:** 0.35ml of the solution was injected onto the column via manual valve
**Detection:** UV Diode Array at 250nm (Band width 80nm, reference 400nm bandwidth 100nm)
**Mobile Phase A:** Heptane + 0.2% Isopropylamine
**Mobile Phase B:** Ethanol + 0.2% Isopropylamine
**Isocratic method:** 98:2 mobile phase A:B
**Runtime:** 20 min

The appropriate fractions were combined and evaporated in vacuo to give the required products:

**Methyl (3*S*,4*S*)-1-(3-cyano-6-methylpyridin-2-yl)-4-isopropylpyrrolidine-3-carboxylate (23)**

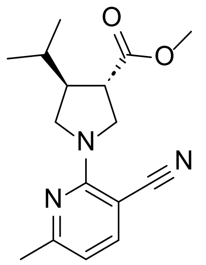

The required product (26.2 mg, 44% yield) was recovered as a white solid.

**$^1$H NMR** (400 MHz, Methanol-d$_4$) δ = 7.66 (d, *J* = 7.8 Hz, 1H), 6.57 (d, *J* = 8.1 Hz, 1H), 4.07 (dd, *J* = 8.2, 10.9 Hz, 1H), 3.98 (dd, *J* = 7.8, 10.8 Hz, 1H), 3.82 (dd, *J* = 8.8, 11.0 Hz, 1H), 3.74 (s, 3H), 3.57 (dd, *J* = 9.0, 10.8 Hz, 1H), 2.99 (q, *J* = 8.6 Hz, 1H), 2.47 - 2.35 (m, 4H), 1.85 - 1.69 (m, 1H), 0.98 (m, 6H). **LCMS** (High pH, 2 min) Rt = 1.31 min, MH+ = 288 (100% a/a). Stereochemistry retrospectively assigned based on subsequent chemistry and ERAP1 co-crystal structure.

**Methyl (3*R*,4*R*)-1-(3-cyano-6-methylpyridin-2-yl)-4-isopropylpyrrolidine-3-carboxylate (24)**

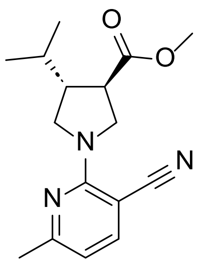

The required product (24.1 mg, 40% yield) was recovered as a white solid.

**$^1$H NMR** (400 MHz, Methanol-d$_4$) δ = 7.67 (d, *J* = 8.1 Hz, 1H), 6.56 (d, *J* = 7.8 Hz, 1H), 4.07 (dd, *J* = 8.1, 11.0 Hz, 1H), 3.98 (dd, *J* = 7.9, 10.9 Hz, 1H), 3.82 (dd, *J* = 8.7, 10.9 Hz, 1H), 3.74 (s, 3H), 3.57 (dd, *J* = 9.0, 10.8 Hz, 1H), 2.99 (q, *J* = 8.6 Hz, 1H), 2.47 - 2.34 (m, 4H), 1.85 - 1.69 (m, 1H), 0.98 (m, 6H). **LCMS** (High pH, 2 min) Rt = 1.30 min, MH+ = 288 (100% a/a). Stereochemistry retrospectively assigned based on subsequent chemistry and ERAP1 co-crystal structure.

**(3*S*,4*S*)-1-(3-Cyano-6-methylpyridin-2-yl)-4-isopropylpyrrolidine-3-carboxylic acid (1)**

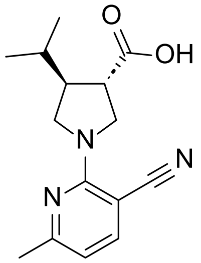

To a solution of methyl (*S,S*)-1-(3-cyano-6-methylpyridin-2-yl)-4-isopropylpyrrolidine-3-carboxylate (26.2 mg, 0.091 mmol) in THF (0.4 mL) was added a solution of lithium hydroxide monohydrate (8.0 mg, 0.191 mmol) in water (0.4 mL), then the mixture was stirred at rt for 18 h. The reaction mixture was directly purified by MDAP (formic), then the appropriate fractions were combined and dried under a stream of nitrogen to give the required product (7.5 mg, 30% yield) as a white solid.

**$^1$H NMR** (Methanol-d$_4$, 400 MHz) δ 7.67 (d, 1H, *J*=7.9 Hz), 6.56 (d, 1H, *J*=7.9 Hz), 4.09 (dd, 1H, *J*=8.1, 11.1 Hz), 3.98 (dd, 1H, *J*=7.9, 10.8 Hz), 3.85 (dd, 1H, *J*=8.6, 11.1 Hz), 3.57 (dd, 1H, *J*=9.1, 10.6 Hz), 2.9-3.0 (m, 1H), 2.4-2.5 (m, 1H), 2.39 (s, 3H), 1.80 (sxtd, 1H, *J*=7.0, 13.8 Hz), 1.02 (m, 6H). **LCMS** (Formic, 2 min): Rt = 1.08 min, MH+ = 274 (100% a/a). Absolute stereochemistry assigned based on ERAP1 co-crystal structure.

**(3*R*,4*R*)-1-(3-Cyano-6-methylpyridin-2-yl)-4-isopropylpyrrolidine-3-carboxylic acid (2)**

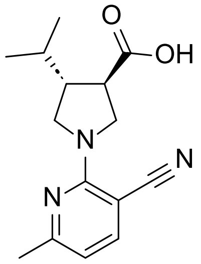

To a solution of methyl (*R,R*)-1-(3-cyano-6-methylpyridin-2-yl)-4-isopropylpyrrolidine-3-carboxylate (24.1 mg, 0.084 mmol) in THF (0.4 mL) was added a solution of lithium hydroxide monohydrate (7.00 mg, 0.167 mmol) in water (0.4 mL) then the mixture was stirred at rt for 18 h. The reaction mixture was directly purified by MDAP (formic), then the appropriate fractions were combined and concentrated under a stream of nitrogen to give the required product (13.7 mg, 60% yield) as a white solid.

**$^1$H NMR** (Methanol-d$_4$, 400 MHz) δ 7.67 (d, 1H, *J*=7.4 Hz), 6.56 (d, 1H, *J*=7.9 Hz), 4.09 (dd, 1H, *J*=7.9, 10.8 Hz), 3.98 (dd, 1H, *J*=7.9, 10.8 Hz), 3.85 (dd, 1H, *J*=8.6, 11.1 Hz), 3.57 (dd, 1H, *J*=8.9, 10.8 Hz), 2.9-3.0 (m, 1H), 2.4-2.5 (m, 1H), 2.39 (s, 3H), 1.79 (sxtd, 1H, *J*=6.8, 13.8 Hz), 1.01 (m, 6H). **LCMS** (Formic, 2 min): Rt = 1.08 min, MH+ = 274 (100% a/a). Absolute stereochemistry assigned based on ERAP1 co-crystal structure.

**Compound 3**

***trans*-1-(3-Cyano-6-methylpyridin-2-yl)-4-isopropylpyrrolidine-3-carboxylic acid (3)**

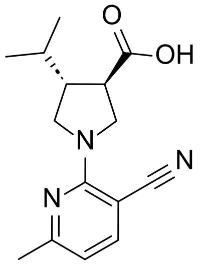

To a solution of methyl *trans*-1-(3-cyano-6-methylpyridin-2-yl)-4-isopropylpyrrolidine-3-carboxylate (24 mg, 0.084 mmol) in a THF:water mixture (1:1, 1.5 mL) was added lithium hydroxide monohydrate (15.0 mg, 0.357 mmol) then the mixture was stirred at rt for 2 h. The mixture was quenched with HCl (4M in 1,4-dioxane, 0.125 mL, 0.500 mmol), then the mixture was concentrated *in vacuo*. The residue was dissolved in minimal DMSO and filtered, then purified by MDAP (formic). The appropriate fractions were combined and concentrated under a stream of nitrogen to give the required product (11 mg, 48% yield) as a white solid.

**$^{1}$H NMR** (DMSO-$d_6$, 400 MHz) δ 12.63 (br s, 1H), 7.80 (d, 1H, *J*=7.8 Hz), 6.61 (d, 1H, *J*=7.8 Hz), 3.98 (dd, 1H, *J*=8.3, 10.8 Hz), 3.87 (dd, 1H, *J*=7.8, 10.8 Hz), 3.75 (dd, 1H, *J*=8.3, 10.8 Hz), 3.46 (dd, 1H, *J*=8.8, 10.8 Hz), 2.8-3.0 (m, 1H), 2.3-2.4 (m, 4H), 1.76 (sxtd, 1H, *J*=6.8, 13.7 Hz), 0.9-1.0 (m, 6H). **LCMS** (Formic, 2 min) Rt = 1.08 min, MH+ = 274 (100% a/a).

**Compound 4**

**Methyl *trans*-1-(3-cyano-6-methoxypyridin-2-yl)-4-isopropylpyrrolidine-3-carboxylate (25)**

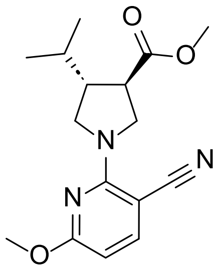

2-Chloro-6-methoxynicotinonitrile (42.0 mg, 0.25 mmol) was added to a solution of *trans*-methyl 4-isopropylpyrrolidine-3-carboxylate (51.0 mg, 0.300 mmol) and DIPEA (0.109 mL, 0.625 mmol) in DMSO (0.7 ml) and heated in a microwave at 120 °C for 1 h. The reaction mixture was directly purified by MDAP (high pH) then the appropriate fractions were combined and concentrated under a stream of nitrogen to give the required product (27.0 mg, 34% yield) as a brown oil.

**$^{1}$H NMR** (400 MHz, Methanol-$d_4$) δ = 7.62 (d, *J* = 8.4 Hz, 1H), 6.08 (d, *J* = 8.4 Hz, 1H), 4.12 (dd, *J* = 8.1, 11.1 Hz, 1H), 4.03 (dd, *J* = 7.9, 10.8 Hz, 1H), 3.89 (s, 3H), 3.84 (dd, *J* = 8.6, 11.1 Hz, 1H), 3.74 (s, 3H), 3.64 - 3.54 (m, 1H), 3.02 (q, *J* = 8.5 Hz, 1H), 2.50 - 2.37 (m, 1H), 1.86 - 1.68 (m, 1H), 0.99 (m, 6H). **LCMS** (High pH, 2 min) Rt = 1.31 min, MH+= 304 (96% a/a).

***trans*-1-(3-Cyano-6-methoxypyridin-2-yl)-4-isopropylpyrrolidine-3-carboxylic acid (4)**

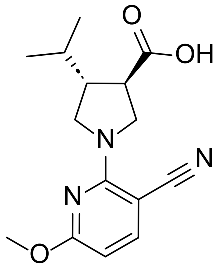

To a solution of methyl *trans*-1-(3-cyano-6-methoxypyridin-2-yl)-4-isopropylpyrrolidine-3-carboxylate (27 mg, 0.089 mmol) in THF (0.4 mL) was added a solution of lithium hydroxide monohydrate (15.0 mg, 0.357 mmol) in water (0.4 mL), then the mixture was stirred at rt for 2 h. Further lithium hydroxide monohydrate (15.0 mg, 0.357 mmol) was added and the mixture stirred at rt for 1 h. The reaction mixture was treated with HCl (4M in 1,4-dioxane, 0.2 mL, 0.8 mmol) with stirring, then the mixture was concentrated under a stream of nitrogen. The residue was dissolved in DMSO:water (1:1, 1.8 mL), filtered and purified by MDAP (formic). The appropriate fractions were combined and evaporated in vacuo to give the required product (19.2 mg, 75% yield) as a white solid.

**$^1$H NMR** (400 MHz, DMSO-d$_6$) δ = 7.75 (d, $J$ = 8.4 Hz, 1H), 6.12 (d, $J$ = 8.4 Hz, 1H), 4.02 (dd, $J$ = 7.9, 10.8 Hz, 1H), 3.91 (dd, $J$ = 7.9, 10.8 Hz, 1H), 3.84 (s, 3H), 3.75 (dd, $J$ = 8.4, 10.8 Hz, 1H), 3.49 (dd, $J$ = 8.4, 10.8 Hz, 1H), 2.89 (q, $J$ = 8.7 Hz, 1H), 2.41 - 2.28 (m, 1H), 1.81 - 1.69 (m, 1H), 0.93 (m, 6H). Carboxylic acid proton not observed. **LCMS** (Formic, 2 min) Rt = 1.14 min, MH+ = 290 (99% a/a).

**<u>Compound 5</u>**

**Methyl *trans*-1-(6-amino-3-cyanopyridin-2-yl)-4-isopropylpyrrolidine-3-carboxylate (26)**

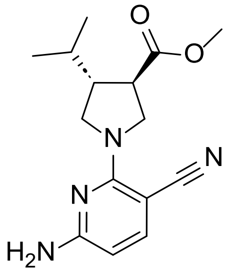

To a stirred solution of 6-amino-2-chloronicotinonitrile (286 mg, 1.86 mmol) in NMP (15 mL) was added *trans*-methyl 4-isopropylpyrrolidine-3-carboxylate (351 mg, 2.049 mmol), then the mixture stirred at 120 °C for 22 h. Further *trans*-methyl 4-isopropylpyrrolidine-3-carboxylate (159 mg, 0.931 mmol) was added, and the mixture stirred for a further 32 h at 120 °C. The mixture was diluted with LiCl (5% aq., 300 mL) and EtOAc (150 mL), then the phases were separated. The aqueous layer was back-extracted with EtOAc (150 mL), then the organic layers were combined, washed with citric acid (5% aq., 300 mL) and LiCl (5% aq., 150 mL), dried using a hydrophobic frit and evaporated in vacuo. The sample was purified by flash chromatography (Si, 40 g) using a 0-50% EtOAc-cyclohexane gradient over 18CV. The appropriate fractions were combined and evaporated in vacuo to give the required product (462 mg, 86% yield) as a pale-yellow solid.

**$^1$H NMR** (400 MHz, DMSO-d$_6$) δ 7.38 (d, J = 8.3 Hz, 1H), 6.58 (br s, 2H), 5.83 (d, J = 8.6 Hz, 1H), 3.93 (dd, J = 8.3, 10.8 Hz, 1H), 3.82 (dd, J = 7.9, 10.6 Hz, 1H), 3.72 - 3.63 (m, 4H), 3.40 (dd, J = 9.2, 10.6 Hz, 1H), 2.97 (q, J = 8.6 Hz, 1H), 2.36 - 2.25 (m, 1H), 1.73 (qd, J = 6.8, 13.7 Hz, 1H), 0.89 (m, 6H). **LCMS** (HpH, 2 min) rt = 1.08 mins, MH+ = 289 (100% a/a).

**_trans_-1-(6-Amino-3-cyanopyridin-2-yl)-4-isopropylpyrrolidine-3-carboxylic acid (5)**

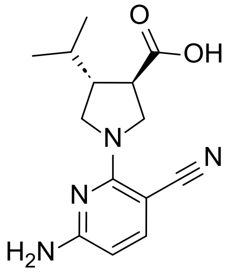

To a stirred solution of methyl *trans*-1-(6-amino-3-cyanopyridin-2-yl)-4-isopropylpyrrolidine-3-carboxylate (452 mg, 1.57 mmol) in THF (10 mL) was added a solution of lithium hydroxide hydrate (132 mg, 3.14 mmol) in water (2 mL), then the mixture was stirred at rt for 4 h. The mixture was diluted with $NaHCO_3$ (sat. aq., 20 mL), water (20 mL) and EtOAc (50 mL), then the phases were separated. The organic layer was back-extracted with $NaHCO_3$ (sat. aq., 50 mL), then the aqueous layers were combined and extracted with further EtOAc (50 mL). The pH of the aqueous layer was adjusted to ~3 by dropwise addition of HCl (25% aq.), then the solution was extracted with EtOAc (3 x 100 mL). These organic extracts were combined and concentrated in vacuo. The sample was purified by chromatography using an XBridge Phenyl column (30x150 mm, 5 μM) using a 15-65% acetonitrile-water (+0.1% formic acid modifier) gradient over 20 min. The appropriate fractions were combined and concentrated under a stream of nitrogen to remove organics. The aqueous solution extracted with EtOAc (3 x 500 mL), then the organic layers were combined, dried over sodium sulfate, filtered and evaporated in vacuo to give the required product (366 mg, 85% yield) as an off-white solid.

**$^1$H NMR** (400 MHz, DMSO-$d_6$) δ = 12.56 (br s, 1H), 7.39 (d, *J* = 8.6 Hz, 1H), 6.59 (br s, 2H), 5.83 (d, *J* = 8.3 Hz, 1H), 3.92 (dd, *J* = 8.2, 10.6 Hz, 1H), 3.81 (dd, *J* = 8.1, 10.8 Hz, 1H), 3.70 (dd, *J* = 8.7, 10.6 Hz, 1H), 3.40 (dd, *J* = 8.8, 10.5 Hz, 1H), 2.86 (q, *J* = 8.4 Hz, 1H), 2.33 - 2.26 (m, 1H), 1.80 - 1.67 (m, 1H), 0.93 (m, 6H). **LCMS** (High pH, 2 min) Rt = 0.56 min, MH+ = 275 (96% a/a).

**<u>Compound 6</u>**

**2-Chloro-6-(isopropylamino)nicotinonitrile (27)**

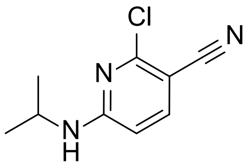

Isopropylamine (0.065 ml, 0.757 mmol) was added to a solution of 2,6-dichloronicotinonitrile (120 mg, 0.694 mmol) and DIPEA (0.424 ml, 2.43 mmol) in DMF (2 mL), then the mixture was heated at 80 °C for 16 h. The reaction mixture was directly purified by MDAP (high pH), then the appropriate fractions were combined and concentrated under a stream of nitrogen to give the required product (116 mg, 85% yield) as a yellow solid.

**$^1$H NMR** (400 MHz, $CDCl_3$) δ = 7.55 (d, J = 8.4 Hz, 1H), 6.27 (d, J = 8.9 Hz, 1H), 5.05 (br s, 1H), 4.08 - 3.88 (m, 1H), 1.26 (d, J = 6.4 Hz, 6H). **LCMS** (High pH, 2 min) Rt = 1.07 min, MH+ = 196 (100% a/a).

***trans*-1-(3-Cyano-6-(isopropylamino)pyridin-2-yl)-4-isopropylpyrrolidine-3-carboxylate (6)**

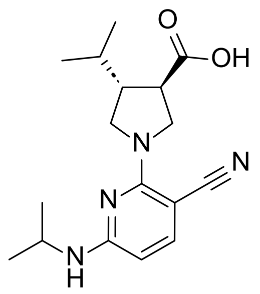

A vial containing 2-chloro-6-(isopropylamino)nicotinonitrile (100 mg, 0.511 mmol), methyl *trans*-4-isopropylpyrrolidine-3-carboxylate (105 mg, 0.613 mmol) and DIPEA (0.268 mL, 1.53 mmol) in NMP (1 mL) was heated at 120 °C for 1 h, then at 180 °C for a further 1 h. The mixture was cooled to rt, then treated with NaOH (2M aq., 1 mL, 2.0 mmol) and the mixture stirred at 70 °C for 1 h. The reaction mixture was directly purified by MDAP (high pH), then the appropriate fractions were combined, diluted with EtOAc (10 mL) and water (10 mL), the pH adjusted to 2 with HCl (2M aq.), then the phases were separated. The aqueous layer was back-extracted with EtOAc (2 x 10 mL), then the organic layers were combined and dried under a stream of nitrogen to give the required product (115 mg, 71% yield) as a white solid.

**$^1$H NMR** (400 MHz, DMSO-d6) δ = 7.32 (d, J = 8.6 Hz, 1H), 7.05 (td, J = 1.0, 1.8 Hz, 1H), 5.82 (d, J = 8.6 Hz, 1H), 4.08 - 3.89 (m, 2H), 3.83 (dd, J = 7.9, 10.6 Hz, 1H), 3.68 (dd, J = 8.3, 10.8 Hz, 1H), 3.43 (dd, J = 8.7, 10.6 Hz, 1H), 2.85 (q, J = 8.5 Hz, 1H), 2.37 - 2.24 (m, 1H), 1.72 (qd, J = 6.9, 13.7 Hz, 1H), 1.14 (dd, J = 1.6, 6.5 Hz, 6H), 0.92 (m, 6H). Carboxylic acid proton not observed. **LCMS** (High pH, 2 min) Rt = 0.75 mins, MH+ = 317 (100% a/a).

**Compound 7**

**2-Chloro-6-(cyclopropylamino)nicotinonitrile (28)**

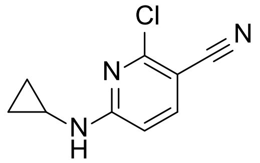

Cyclopropylamine (50.0 µL, 0.722 mmol) was added to a solution of 2,6-dichloronicotinonitrile (120 mg, 0.694 mmol) and DIPEA (0.424 mL, 2.43 mmol) in DMF (2 mL), then the mixture was heated at 80 °C for 16 h. The reaction mixture was directly purified by MDAP (high pH), then the appropriate fractions were combined and concentrated under a stream of nitrogen to give the required product (69 mg, 51% yield) as a white solid.

**$^1$H NMR** (400 MHz, $CDCl_3$) δ = 7.68 (d, *J* = 8.4 Hz, 1H), 6.68 (d, *J* = 8.4 Hz, 1H), 5.57 (br s, 1H), 2.64 - 2.54 (m, 1H), 0.95 - 0.87 (m, 2H), 0.66 - 0.59 (m, 2H). **LCMS** (HpH, 2 min) Rt = 0.97 min, MH+ = 194 (100% a/a).

**Methyl *trans*-1-(3-cyano-6-(cyclopropylamino)pyridin-2-yl)-4-isopropylpyrrolidine-3-carboxylate (29)**

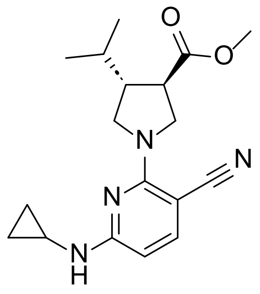

To a solution of 2-chloro-6-(cyclopropylamino)nicotinonitrile (68 mg, 0.351 mmol) and DIPEA (0.153 mL, 0.878 mmol) in DMSO (0.7 mL) was added methyl trans-4-isopropylpyrrolidine-3-carboxylate (60 mg, 0.350 mmol) then the mixture was heated in a microwave at 120 °C for 1 h. The reaction mixture was directly purified by MDAP (formic), then the appropriate fractions were combined and dried under a stream of nitrogen to give the required product (44 mg, 38% yield) as a colourless oil.

**$^1$H NMR** (400 MHz, $CDCl_3$) δ = 7.49 (d, *J* = 8.9 Hz, 1H), 6.09 (d, *J* = 8.9 Hz, 1H), 5.11 (s, 1H), 4.07 (dd, *J* = 8.4, 10.8 Hz, 1H), 3.96 (dd, *J* = 8.4, 10.8 Hz, 1H), 3.82 (dd, *J* = 9.1, 11.1 Hz, 1H), 3.73 (s, 3H), 3.50 (dd, *J* = 8.9, 10.8 Hz, 1H), 2.88 (q, *J* = 8.9 Hz, 1H), 2.58 - 2.50 (m, 1H), 2.48 - 2.36 (m, 1H), 1.82 - 1.65 (m, 1H), 0.96 (m, 6H), 0.84 - 0.75 (m, 2H), 0.60 - 0.52 (m, 2H). **LCMS** (High pH, 2 min) Rt = 1.31 min, MH+ = 329 (98% a/a).

***trans*-1-(3-Cyano-6-(cyclopropylamino)pyridin-2-yl)-4-isopropylpyrrolidine-3-carboxylic acid (7)**

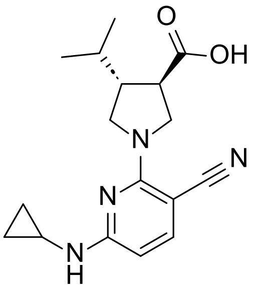

To a solution of methyl *trans*-1-(3-cyano-6-(cyclopropylamino)pyridin-2-yl)-4-isopropylpyrrolidine-3-carboxylate (22 mg, 0.067 mmol) in THF (0.4 mL) was added a solution of lithium hydroxide monohydrate (11.0 mg, 0.262 mmol) in water (0.4 mL) at room temperature with stirring. The reaction mixture was left to stir at room temperature for 1 h. The reaction mixture was directly purified by MDAP (formic), then the appropriate fractions were combined and concentrated under a stream of nitrogen to give the required product (19.5 mg, 93% yield) as a colourless oil.

**$^1$H NMR** (400 MHz, $CDCl_3$) δ = 7.50 (d, *J* = 8.6 Hz, 1H), 6.11 (d, *J* = 8.6 Hz, 1H), 5.30 (br s, 1H), 4.08 (dd, *J* = 8.2, 11.1 Hz, 1H), 3.98 (dd, *J* = 7.9, 10.9 Hz, 1H), 3.88 (dd, *J* = 8.4, 11.1 Hz, 1H), 3.55 (dd, *J* = 8.8, 10.8 Hz, 1H), 2.99 - 2.85 (m, 1H), 2.57 - 2.50 (m, 1H), 2.50 - 2.41 (m, 1H), 1.86 - 1.72 (m, 1H), 1.00 (m, 6H), 0.87 - 0.73 (m, 2H), 0.60 - 0.54 (m, 2H). Carboxylic acid proton not observed. **LCMS** (Formic, 2 min) Rt = 1.10 min, MH+ = 315 (100% a/a).

**Compound 8**

**(*R*,*E*)-3-(4-Methylpent-2-enoyl)-4-phenyloxazolidin-2-one (30)**

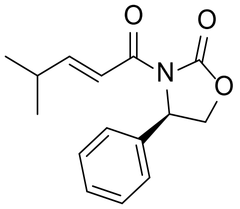

To a stirred solution of (*E*)-4-methylpent-2-enoic acid (10.0 g, 88.0 mmol) DCM (150 mL) was added oxalyl chloride (16.1 mL, 184 mmol) at 0 °C followed by DMF (0.678 mL, 8.76 mmol). The reaction mixture was stirred at room temperature for 2 h. The reaction mixture was concentrated in vacuo then the residue was dissolved in DCM (100 mL).

In a separate flask, sodium hydride (60% in mineral oil, 2.45 g, 61.3 mmol) was suspended in THF (150 mL) then (*R*)-4-phenyloxazolidin-2-one (10.0 g, 61.3 mmol) added in portions over 15 min. After stirring for a further 5 min, the solution of (*E*)-4-methylpent-2-enoyl chloride in DCM (100 mL) was added then the mixture was stirred at rt for 2 h. The reaction mixture was quenched with 2 M HCl (2M aq., 100 mL) then diluted with water (100 mL) and DCM (100 mL). The phases were separated and the aqueous layer back-extracted with DCM (4 x 100 mL). The organic layers were combined, washed with $NaHCO_3$ (sat. aq., 2 x 200 mL) dried using a hydrophobic frit and concentrated in vacuo. The sample was purified by flash chromatography (Si, 330g) using a 0-20% EtOAc-cyclohexane gradient over 20 CV. The appropriate fractions were combined and concentrated in vacuo to give (R,E)-3-(4-methylpent-2-enoyl)-4-phenyloxazolidin-2-one (12.8 g, 80% yield) as a white solid.

**$^1$H NMR** (400 MHz, DMSO-$d_6$) δ = 7.41 - 7.35 (m, 2H), 7.34 - 7.27 (m, 3H), 7.19 - 7.08 (m, 1H), 6.91 (dd, *J* = 6.4, 15.7 Hz, 1H), 5.51 (dd, *J* = 3.7, 8.6 Hz, 1H), 4.76 (t, *J* = 8.6 Hz, 1H), 4.17 (dd, *J* = 3.7, 8.6 Hz, 1H), 1.03 (d, *J* = 6.4 Hz, 6H). One additional multiplet coincides with DMSO solvent peak. **LCMS** (Formic, 2 min) Rt = 1.16 min, MH+ = 260 (95% a/a).

**(*R*)-3-((3*S*,4*S*)-1-Benzyl-4-isopropylpyrrolidine-3-carbonyl)-4-phenyloxazolidin-2-one (31) and (*R*)-3-((3*R*,4*R*)-1-Benzyl-4-isopropylpyrrolidine-3-carbonyl)-4-phenyloxazolidin-2-one (32)**

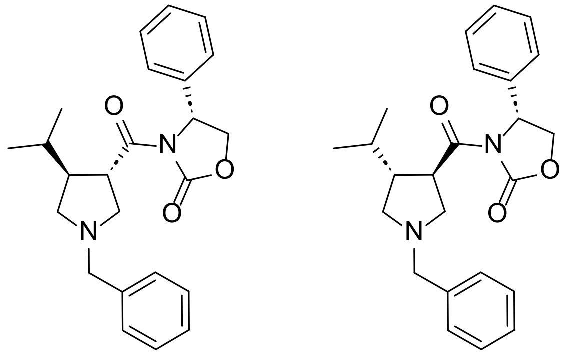

A solution of (*R*,*E*)-3-(4-methylpent-2-enoyl)-4-phenyloxazolidin-2-one (12.8 g, 49.3 mmol) in DCM (100 mL) was cooled to 0 °C then *N*-benzyl-1-methoxy-*N*-((trimethylsilyl)methyl)methanamine (18.9 mL, 73.9 mmol) and TFA (0.759 mL, 9.86 mmol) added. The reaction mixture was stirred at 0 °C for 15 min, then at rt for 4 h. Further *N*-benzyl-1-methoxy-*N*-((trimethylsilyl)methyl)methanamine (6.30 mL, 24.6 mmol) was added and the reaction mixture stirred at rt for 1 h. The reaction mixture was diluted with DCM (200 mL), then washed with $NaHCO_3$ (sat. aq., 200 mL). The aqueous layer was back-extracted with DCM (100 mL), then organic layers were combined, dried using a hydrophobic frit and evaporated in vacuo. The sample was purified by flash chromatography (Si, 330g) using a 0-50% EtOAc-cyclohexane over 20 CV. The appropriate fractions were combined and concentrated in vacuo to give the required products:

(*R*)-3-((3*S*,4*S*)-1-Benzyl-4-isopropylpyrrolidine-3-carbonyl)-4-phenyloxazolidin-2-one (13.67 g, 64% yield) as a white solid.

**$^1$H NMR** (400 MHz, DMSO-$d_6$) δ = 7.41 - 7.17 (m, 10H), 5.47 (dd, *J* = 4.2, 8.6 Hz, 1H), 4.76 (t, *J* = 8.8 Hz, 1H), 4.15 - 4.09 (m, 1H), 3.73 - 3.66 (m, 1H), 3.64 - 3.56 (m, 2H), 2.82 - 2.73 (m, 2H), 2.62 - 2.56 (m, 1H), 2.49 - 2.42 (m, 1H), 2.16 - 2.10 (m, 1H), 1.56 - 1.45 (m, 1H), 0.81 (m, 6H). **LCMS** (Formic, 2 min) Rt = 0.71 min, MH+ = 393 (100% a/a).

(*R*)-3-((3*R*,4*R*)-1-benzyl-4-isopropylpyrrolidine-3-carbonyl)-4-phenyloxazolidin-2-one (6.10 g, 25% yield) as an off-white solid.

**$^1$H NMR** (400 MHz, DMSO-$d_6$) δ = 7.41 - 7.21 (m, 10H), 5.46 (dd, *J* = 3.9, 8.8 Hz, 1H), 4.73 (t, *J* = 8.8 Hz, 1H), 4.16 - 4.10 (m, 1H), 3.87 - 3.78 (m, 1H), 3.58 - 3.46 (m, 2H), 2.82 - 2.72 (m, 2H), 2.69 - 2.63 (m, 1H), 2.35 - 2.24 (m, 1H), 2.22 - 2.17 (m, 1H), 1.51 - 1.41 (m, 1H), 0.74 (d, *J* = 6.4 Hz, 3H), 0.62 (d, *J* = 6.4 Hz, 3H). **LCMS** (Formic, 2 min) Rt = 0.72 min, MH+ = 393 (94% a/a).

Absolute stereochemistry of compounds **31** and **32** confirmed by VCD, see later section.

**Methyl (3*R*,4*R*)-1-benzyl-4-isopropylpyrrolidine-3-carboxylate (33)**

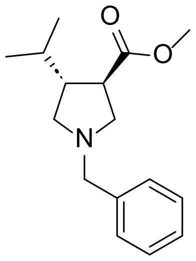

To a stirred solution of (*R*)-3-((3*R*,4*R*)-1-benzyl-4-isopropylpyrrolidine-3-carbonyl)-4-phenyloxazolidin-2-one (6.10 g, 15.5 mmol) in THF (150 mL) was added a mixture of lithium hydroxide (1M aq., 38.9 mL, 38.9 mmol) and hydrogen peroxide (30% aq., 3.17 mL, 31.1 mmol) in water (50 mL) dropwise over 15 min at 0 °C, then the mixture was stirred at 0 °C for 1 h. The reaction mixture was quenched with a solution of sodium metabisulfite (5.91 g, 31.1 mmol) in water (200 mL) then extracted with EtOAc (200 mL). The aqueous layer was acidified to pH 5 using HCl (2M aq.) then extracted with isopropanol:DCM (1:3, 12 x 100 mL). The organic layers were combined, dried using a hydrophobic frit and evaporated *in vacuo* give crude (3*R*,4*R*)-1-benzyl-4-isopropylpyrrolidine-3-carboxylic acid (3.29 g) as a pale orange solid that was used directly in the next step.

In a separate flask, thionyl chloride (4.85 mL, 66.5 mmol) was added dropwise at 0 °C to MeOH (40 mL). A solution of crude (3*R*,4*R*)-1-benzyl-4-isopropylpyrrolidine-3-carboxylic acid (3.29 g) in MeOH (40 mL) was then added in portions at 0 °C. The ice bath was removed, and the reaction mixture was stirred at 40 °C for 1 h. The reaction mixture was cooled to rt then concentrated in vacuo. The sample was purified by flash chromatography (KP-NH, 110g) using a 0-50% EtOAc-cyclohexane over 20 CV. The appropriate fractions were combined and evaporated in vacuo to give the required product (1.33 g, 33% yield over two steps) as a pale-yellow oil.

**$^1$H NMR** (400 MHz, DMSO-d$_6$) δ = 7.35 - 7.19 (m, 5H), 3.60 (s, 3H), 3.58 - 3.44 (m, 2H), 2.75 - 2.56 (m, 4H), 2.23 - 2.12 (m, 2H), 1.64 - 1.47 (m, 1H), 0.83 (m, 6H). **LCMS** (High pH, 2 min) Rt = 1.34 min, MH+ = 262 (99% a/a).

**Methyl (3*R*,4*R*)-4-isopropylpyrrolidine-3-carboxylate (34)**

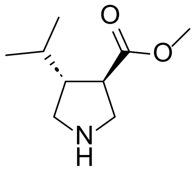

To a flask containing palladium hydroxide on carbon (20% wt, 0.357 g, 0.509 mmol) and ammonium formate (0.963 g, 15.3 mmol) was added a solution of methyl (3*R*,4*R*)-1-benzyl-4-isopropylpyrrolidine-3-carboxylate (1.33 g, 5.09 mmol) in EtOH (25 mL), then the mixture was stirred at 55 °C for 2 h. The reaction mixture was cooled to rt then filtered through celite (10 g cartridge, preconditioned with EtOH), eluting with EtOH. The filtrate was concentrated *in vacuo* then redissolved in MeOH (5 mL) and purified by solid phase extraction (SCX cartridge, 10g) eluting sequentially with MeOH (150 mL) then ammonia in MeOH (4M, 150 mL). The ammonia fractions were combined and concentrated *in vacuo* to give the required product (707 mg, 81% yield) as a pale-yellow oil.

**$^1$H NMR** (400 MHz, Methanol-d$_4$) δ = 3.68 (s, 3H), 3.13 (dd, *J* = 7.8, 11.2 Hz, 1H), 3.08 - 3.02 (m, 2H), 2.73 - 2.64 (m, 1H), 2.55 (dd, *J* = 8.3, 11.7 Hz, 1H), 2.19 - 2.07 (m, 1H), 1.69 - 1.53 (m, 1H), 0.93 (m, 6H). **LCMS** (HpH, 2min) Rt = 0.71 min, MH+ = 172 (92% a/a; expect modest chromophore. NMR indicates >95% purity).

**(3*R*,4*R*)-1-(3-Cyano-6-(cyclopropylamino)pyridin-2-yl)-4-isopropylpyrrolidine-3-carboxylic acid (8)**

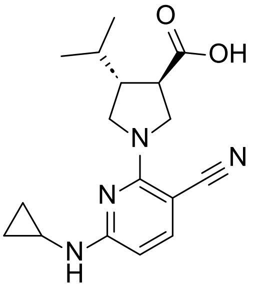

To a solution of 2-chloro-6-(cyclopropylamino)nicotinonitrile (69 mg, 0.356 mmol) and DIPEA (0.187 mL, 1.07 mmol) in DMSO (1.2 mL) was added methyl (3*R*,4*R*)-4-isopropylpyrrolidine-3-carboxylate (65 mg, 0.380 mmol), then the mixture was heated in the microwave at 120 °C for 1 h. The mixture was cooled to rt, then NaOH (10M aq., 0.143 mL, 1.43 mmol) was added and the mixture heated at 40 °C for 1 h. The reaction mixture was diluted with DMF (0.5 mL) and purified by MDAP (formic). The appropriate fractions were combined and dried under a stream of nitrogen to give the required product (58.6 mg, 52% yield) as a white solid.

**$^1$H NMR** (400 MHz, CDCl$_3$) δ = 7.49 (d, *J* = 8.4 Hz, 1H), 6.10 (d, *J* = 8.4 Hz, 1H), 5.26 (br s, 1H), 4.07 (dd, *J* = 8.1, 11.1 Hz, 1H), 3.97 (dd, *J* = 7.9, 10.8 Hz, 1H), 3.88 (dd, *J* = 8.6, 11.1 Hz, 1H), 3.54 (dd, *J* = 8.9, 10.8 Hz, 1H), 2.90 (q, *J* = 8.5 Hz, 1H), 2.56 - 2.49 (m, 1H), 2.49 - 2.39 (m, 1H), 1.85 - 1.71 (m, 1H), 0.99 (m, 6H), 0.82 - 0.75 (m, 2H), 0.60 - 0.52 (m, 2H). Carboxylic acid proton not observed. **LCMS** (Formic, 2 min) Rt = 1.13 min, MH+ = 315 (100% a/a).

__**Compound 9**__

**(3*S*,4*S*)-1-Benzyl-4-isopropylpyrrolidine-3-carboxylic acid (35)**

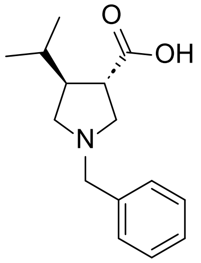

To a solution of (*R*)-3-((3*S*,4*S*)-1-benzyl-4-isopropylpyrrolidine-3-carbonyl)-4-phenyloxazolidin-2-one (1.19 g, 3.03 mmol) in THF (25 ml) was added lithium hydroxide (1M aq., 7.58 mL, 7.58 mmol) followed by hydrogen peroxide (30% aq., 0.619 mL, 6.06 mmol) in water (8.3 mL) portion wise at 0 °C with stirring. The reaction mixture was stirred at 0 °C for 1 h, then was quenched with a solution of sodium metabisulfite (1.15 g, 6.06 mmol) in water (35 mL) and the mixture was extracted with EtOAc. The aqueous phase was adjusted to pH~5 with $K_2HPO_4$ (1.36 g, 10.0 mmol) and HCl (10% aq., ~1 mL). This solution was extracted with isopropanol:DCM (1:3, 3 x 60 ml). The organic phase was combined, dried with hydrophobic frit and evaporated under reduced pressure to give the required product (390 mg, 52% yield) as a pale-yellow solid which was used directly in the next reaction.

**LCMS** (High pH, 2 min) Rt = 0.64 min, MH+ = 248 (87% a/a, expect modest chromophore).

**Methyl (3*S*,4*S*)-1-benzyl-4-isopropylpyrrolidine-3-carboxylate (36)**

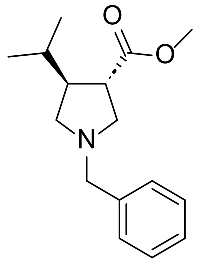

Thionyl chloride (0.575 mL, 7.88 mmol) was added dropwise at 0 °C to MeOH (6 mL). (3*S*,4*S*)-1-benzyl-4-isopropylpyrrolidine-3-carboxylic acid (390 mg, 1.58 mmol) was then added portion wise at 0 °C. The ice bath was removed, then the mixture was heated at 40 °C for 1 h. The mixture was concentrated in vacuo, then the residue was purified by MDAP (high pH) then the appropriate fractions were combined and evaporated in vacuo to give the required product (244 mg, 59% yield) as a colourless oil.

**$^1$H NMR** (400 MHz, $CDCl_3$) δ = 7.38 - 7.21 (m, 5H), 3.70 (s, 3H), 3.69 - 3.64 (m, 1H), 3.58 - 3.53 (m, 1H), 2.87 - 2.77 (m, 2H), 2.77 - 2.69 (m, 2H), 2.38 - 2.29 (m, 2H), 1.67 - 1.58 (m, 1H), 0.91 (m, 6H). **LCMS** (High pH, 2 min) Rt = 1.33 min, MH+ = 262 (100% a/a).

**Methyl (3*S*,4*S*)-4-isopropylpyrrolidine-3-carboxylate (37)**

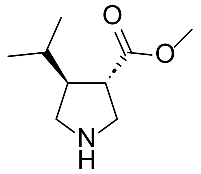

To a stirred solution of methyl (3*S*,4*S*)-1-benzyl-4-isopropylpyrrolidine-3-carboxylate (210 mg, 0.803 mmol) in MeOH (2 mL) was added ammonium formate (152 mg, 2.41 mmol) and Pd/C (5% wt, 86 mg, 0.040 mmol) then the mixture was stirred at 60 °C for 2 h. The reaction mixture was cooled to rt, passed through a pad of Celite and the filtrate concentrated *in vacuo* to give the required product (97 mg, 71% yield) as a colourless oil.

**$^1$H NMR** (400 MHz, $CDCl_3$) δ = 3.69 (s, 3H), 3.25 - 3.10 (m, 2H), 3.07 - 2.99 (m, 1H), 2.66 - 2.57 (m, 1H), 2.53 (dd, *J* = 8.4, 11.3 Hz, 1H), 2.16 - 2.04 (m, 1H), 1.62 - 1.53 (m, 2H), 0.92 (m, 6H). **LCMS** (High pH, 2 min) m/z not observed, not detected by UV.

**(3*S*,4*S*)-1-(3-Cyano-6-(cyclopropylamino)pyridin-2-yl)-4-isopropylpyrrolidine-3-carboxylic acid (9)**

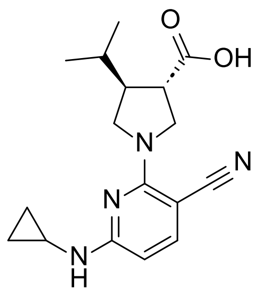

A solution of 2-chloro-6-(cyclopropylamino)nicotinonitrile (34.3 mg, 0.177 mmol), methyl (3*S*,4*S*)-4-isopropylpyrrolidine-3-carboxylate (30 mg, 0.175 mmol) and DIPEA (0.061 mL, 0.350 mmol) in NMP (0.4 mL) was heated at 150 °C for 2 h. The sample was cooled to room temperature, then treated with a solution of lithium hydroxide hydrate (14.7 mg, 0.350 mmol) in water (0.4 mL) and the mixture stirred at rt for 2 h. The mixture was partitioned between EtOAc (5 mL) and HCl (2M aq., 10 mL), then the phases were separated. The aqueous layer was back-extracted with EtOAc (2 x 5 mL), then the organic layers were combined and dried under a stream of nitrogen. The sample was loaded in DMSO and purified by reverse phase chromatography (C18, 12g) using a 30-60% acetonitrile-water (+0.1% formic acid modifier) gradient over 10CV. The appropriate fractions were combined and evaporated in vacuo to give the required product (23 mg, 42% yield) as a pale brown solid.

**$^1$H NMR** (400 MHz, DMSO-$d_6$) δ = 12.58 (br s, 1H), 7.44 (br d, *J* = 7.4 Hz, 1H), 7.30 (br s, 1H), 5.95 (br s, 1H), 3.94 (dd, *J* = 8.4, 10.3 Hz, 1H), 3.84 (dd, *J* = 7.9, 10.8 Hz, 1H), 3.70 (dd, *J* = 8.4, 10.8 Hz, 1H), 3.46 - 3.38 (m, 1H), 2.85 (q, *J* = 8.4 Hz, 1H), 2.65 - 2.56 (m, 1H), 2.35 - 2.24 (m, 1H), 1.79 - 1.66 (m, 1H), 0.92 (m, 6H), 0.72 - 0.64 (m, 2H), 0.47 - 0.39 (m, 2H). **LCMS** (Formic, 2 min) Rt = 1.10 min, MH+ = 315 (100% a/a).

**Compound 10**

**2-Chloro-6-(cyclopentylamino)nicotinonitrile (38)**

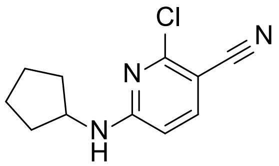

A solution of 2,6-dichloronicotinonitrile (80.0 mg, 0.462 mmol), cyclopentylamine (45.7 µL, 0.462 mmol) and DIPEA (161 µL, 0.925 mmol) in DMF (0.5 mL) was heated at 80 °C for 16 h. The reaction mixture was directly purified by MDAP (high pH), then the appropriate fractions were combined and concentrated under a stream of nitrogen to give the required product (73.2 mg, 71% yield) as an off-white solid.

**$^1$H NMR** (400 MHz, $CDCl_3$) δ = 7.56 (d, *J* = 8.6 Hz, 1H), 6.29 (d, *J* = 8.6 Hz, 1H), 5.21 (br s, 1H), 4.03 (br s, 1H), 2.15 - 1.99 (m, 2H), 1.82 - 1.60 (m, 4H), 1.54 - 1.43 (m, 2H). **LCMS** (High pH, 2 min) Rt = 1.20 min, MH+ = 222 (100% a/a).

***trans*-1-(3-Cyano-6-(cyclopentylamino)pyridin-2-yl)-4-isopropylpyrrolidine-3-carboxylic acid (10)**

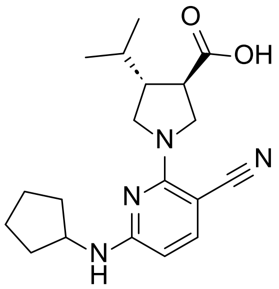

A solution of 2-chloro-6-(cyclopentylamino)nicotinonitrile (44 mg, 0.198 mmol), methyl *trans*-4-isopropylpyrrolidine-3-carboxylate (37.4 mg, 0.218 mmol) and DIPEA (0.069 mL, 0.397 mmol) in NMP (0.4 mL) was heated in the microwave at 180 °C for 1 h. The mixture was cooled to rt, then was treated with NaOH (2M aq., 0.4 mL, 0.8 mmol) and heated to 70 °C for 1 h. The reaction mixture was directly purified by MDAP (formic)

then the appropriate fractions were combined and concentrated under a stream of nitrogen to give the required product (19.3 mg, 28% yield) as an off-white solid.

**$^1$H NMR** (400 MHz, Methanol-$d_4$) δ = 7.29 (d, *J* = 8.6 Hz, 1H), 5.84 (d, *J* = 8.6 Hz, 1H), 4.26 - 4.14 (m, 1H), 4.12 - 4.03 (m, 1H), 4.02 - 3.92 (m, 1H), 3.86 - 3.77 (m, 1H), 3.59 - 3.50 (m, 1H), 2.97 - 2.86 (m, 1H), 2.47 - 2.35 (m, 1H), 2.08 - 1.95 (m, 2H), 1.85 - 1.48 (m, 8H), 1.06 - 0.99 (m, 6H). **LCMS** (High pH, 2 min) Rt = 0.81 mins, MH+ = 343 (99% a/a).

**Compound 11**

**2-Chloro-6-(cyclohexylamino)nicotinonitrile (39)**

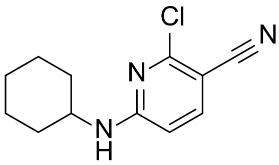

To a solution of 2,6-dichloronicotinonitrile (50 mg, 0.289 mmol) and DIPEA (0.151 mL, 0.867 mmol) in DMF (0.6 mL) was added cyclohexylamine (33.0 µL, 0.289 mmol), was heated in the microwave at 80 °C for 1 h. The reaction mixture was directly purified by MDAP (high pH), then the appropriate fractions were combined and dried under a stream of nitrogen to give the required product (40.4 mg, 59% yield) as a white solid.

**$^1$H NMR** (400 MHz, $CDCl_3$) δ = 7.55 (d, *J* = 8.9 Hz, 1H), 6.27 (d, *J* = 8.9 Hz, 1H), 5.10 (br s, 1H), 3.76 - 3.47 (m, 1H), 2.07 - 1.95 (m, 2H), 1.85 - 1.72 (m, 2H), 1.72 - 1.62 (m, 1H), 1.49 - 1.35 (m, 2H), 1.33 - 1.18 (m, 3H). **LCMS** (High pH, 2 min) Rt = 1.26 min, MH+ = 236 (100% a/a).

***trans*-1-(3-Cyano-6-(cyclohexylamino)pyridin-2-yl)-4-isopropylpyrrolidine-3-carboxylic acid (11)**

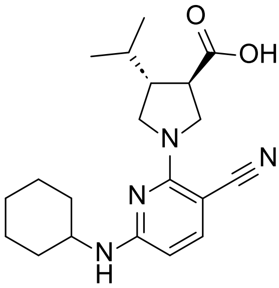

A solution of methyl *trans*-4-isopropylpyrrolidine-3-carboxylate (20 mg, 0.117 mmol), 2-chloro-6-(cyclohexylamino)nicotinonitrile (20 mg, 0.085 mmol) and DIPEA (0.037 mL, 0.212 mmol) in DMSO (0.5 mL) was heated in the microwave to 120 °C for 1 h. The reaction mixture was directly purified by MDAP (high pH), then the appropriate fractions were combined and concentrated under a stream of nitrogen. The residue was directly dissolved in THF (1 mL) and treated with NaOH (2M aq., 0.5 mL, 1.0 mmol) and the mixture stirred at rt for 60 h. The mixture was diluted with EtOAc (5 mL) and water (5 mL), the pH adjusted to ~2 with HCl (2M aq.), then the phases were separated. The aqueous layer was back-extracted with EtOAc (5 mL), then the organic layers were combined and dried under a stream of nitrogen to give the required product (21.3 mg, 70% yield) as a colourless solid.

**$^1$H NMR** (400 MHz, $CDCl_3$) δ = 7.42 (br d, *J* = 6.4 Hz, 1H), 5.76 (br d, *J* = 7.9 Hz, 1H), 4.23 - 3.87 (m, 3H), 3.72 - 3.44 (m, 2H), 3.03 - 2.88 (m, 1H), 2.48 (quin, *J* = 8.0 Hz, 1H), 2.05 - 1.94 (m, 2H), 1.89 - 1.73 (m, 3H), 1.68 - 1.59 (m, 1H), 1.41 - 1.17 (m, 6H), 1.06 - 0.96 (m, 6H). Carboxylic acid proton not observed. **LCMS** (High pH, 2 min) Rt = 0.87 min, MH- = 355.4 (98% a/a)

#### Compound 12

**(*R*)-2-Chloro-6-((tetrahydrofuran-3-yl)amino)nicotinonitrile (40)**

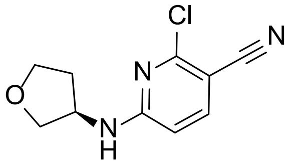

A solution of 2,6-dichloronicotinonitrile (30.0 mg, 0.173 mmol) in DMF (1 mL) was added to (*R*)-tetrahydrofuran-3-amine hydrochloride (21.0 mg, 0.173 mmol). DIPEA (91 µL, 0.52 mmol) was added then the mixture was heated at 80 °C for 16 h. The reaction mixture was directly purified by MDAP (high pH), then the appropriate fractions were combined and concentrated under a stream of nitrogen to give the required product (21.8 mg, 56% yield) which was used directly in the next reaction.

**LCMS** (High pH, 2 min) Rt = 0.78 min, MH+ = 224 (100% a/a).

***trans*-1-(3-Cyano-6-(((*R*)-tetrahydrofuran-3-yl)amino)pyridin-2-yl)-4-isopropylpyrrolidine-3-carboxylic acid (12)**

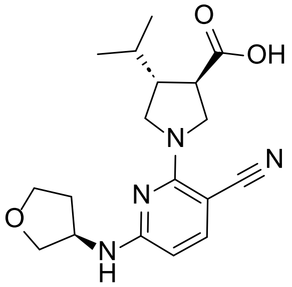

A mixture of (*R*)-2-chloro-6-((tetrahydrofuran-3-yl)amino)nicotinonitrile (21.8 mg 0.097 mmol), methyl *trans*-4-isopropylpyrrolidine-3-carboxylate (26 mg, 0.15 mmol), and DIPEA (79 µL, 0.45 mmol) dissolved in DMSO (0.5 mL) was heated in the microwave at 120 °C for 30 min. After cooling the reaction added additional methyl *trans*-4-isopropylpyrrolidine-3-carboxylate (26 mg, 0.15 mmol) and DIPEA (79 µL, 0.45 mmol) to each reaction mixtures and heated in the microwave at 130 °C for 30 min. Further DIPEA (79 µL, 0.45 mmol) was added to the mixture heated in the microwave at 140 °C for 30 min. Further DIPEA (79 µL, 0.45 mmol) was added to the mixture heated in the microwave at 150 °C for 30 min. The mixture was cooled to rt then NaOH (10 M aq., 100 µL) was added, then the mixture heated at 40 °C for 1 h. The reaction mixture was diluted with DMSO (0.4 mL), filtered and purified by MDAP (high pH). The appropriate fractions were combined and concentrated under a stream of nitrogen to give the required product (11.5 mg, 34% yield) a pale brown gum.

**$^1$H NMR** (400 MHz, Methanol-$d_4$) δ = 7.41 (d, *J* = 8.6 Hz, 1H), 5.84 (d, *J* = 8.8 Hz, 1H), 4.53 - 4.44 (m, 1H), 4.11 - 4.02 (m, 1H), 3.99 - 3.91 (m, 1H), 3.83 - 3.74 (m, 1H), 3.63 - 3.44 (m, 5H), 2.84 (q, *J* = 8.9 Hz, 1H), 2.48 - 2.31 (m, 1H), 2.17 - 1.95 (m, 2H), 1.83 - 1.69 (m, 1H), 1.04 - 0.96 (m, 6H). **LCMS** (Formic, 2 min) Rt = 0.96 min, MH+ = 345 (100% a/a).

#### Compound 13

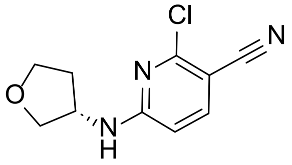

**(*S*)-2-Chloro-6-((tetrahydrofuran-3-yl)amino)nicotinonitrile (41)**

A solution of 2,6-dichloronicotinonitrile (30.0 mg, 0.173 mmol) in DMF (1 mL) was added to (*S*)-tetrahydrofuran-3-amine hydrochloride (21.0 mg, 0.173 mmol). DIPEA (91 µL, 0.52 mmol) was added then the mixture was heated at 80 °C for 16 h. The reaction mixture was directly purified by MDAP (high pH), then the appropriate fractions were combined and concentrated under a stream of nitrogen to give the required product (18.5 mg, 48% yield) which was used directly in the next reaction.

**LCMS** (High pH, 2 min) Rt = 0.86 min, MH+ = 224 (100% a/a)

***trans*-1-(3-Cyano-6-(((*S*)-tetrahydrofuran-3-yl)amino)pyridin-2-yl)-4-isopropylpyrrolidine-3-carboxylic acid (13)**

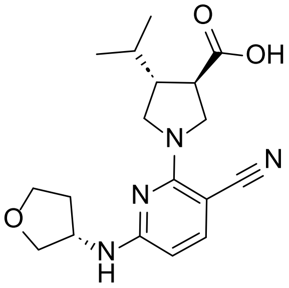

A mixture of (*S*)-2-chloro-6-((tetrahydrofuran-3-yl)amino)nicotinonitrile (18.5 mg, 0.083 mmol), methyl *trans*-4-isopropylpyrrolidine-3-carboxylate (0.026 g, 0.150 mmol), and DIPEA (79 µL, 0.45 mmol) dissolved in DMSO (0.5 mL) was heated in the microwave at 120 °C for 30 min. After cooling the reaction added additional methyl *trans*-4-isopropylpyrrolidine-3-carboxylate (26 mg, 0.15 mmol) and DIPEA (79 µL, 0.45 mmol) to each reaction mixtures and heated in the microwave at 130 °C for 30 min. Further DIPEA (79 µL, 0.45 mmol) was added to the mixture heated in the microwave at 140 °C for 30 min. Further DIPEA (79 µL, 0.45 mmol) was added to the mixture heated in the microwave at 150 °C for 30 min. The mixture was cooled to rt then NaOH (10 M aq., 100 µL) was added, then the mixture heated at 40 °C for 1 h. The reaction mixture was diluted with DMSO (0.4 mL), filtered and purified by MDAP (high pH). The appropriate fractions were combined and concentrated under a stream of nitrogen to give the required product (13.1 mg, 46% yield) as a white solid.

**$^1$H NMR** (400 MHz, Methanol-$d_4$) δ = 7.31 (d, *J* = 8.6 Hz, 1H), 5.86 (d, *J* = 8.6 Hz, 1H), 4.56 - 4.40 (m, 1H), 4.11 - 4.01 (m, 1H), 4.01 - 3.87 (m, 3H), 3.86 - 3.75 (m, 2H), 3.69 - 3.61 (m, 1H), 3.59 - 3.49 (m, 1H), 2.91 (q, *J* = 8.8 Hz, 1H), 2.46 - 2.34 (m, 1H), 2.32 - 2.19 (m, 1H), 1.95 - 1.85 (m, 1H), 1.84 - 1.70 (m, 1H), 1.00 (m, 6H). **LCMS** (Formic, 2 min) Rt = 1.00 min, MH+ = 345 (100% a/a).

**Compound 14**

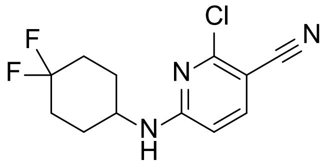

**2-Chloro-6-((4,4-difluorocyclohexyl)amino)nicotinonitrile (42)**

A solution of 2,6-dichloronicotinonitrile (60 mg, 0.347 mmol), 4,4-difluorocyclohexylamine hydrochloride (59.5 mg, 0.347 mmol) and DIPEA (0.182 mL, 1.04 mmol) in DMF (0.4 mL) was heated in a microwave at 80 °C for 1 h. The reaction mixture was directly purified by MDAP (high pH), then the appropriate fractions were combined and concentrated under a stream of nitrogen to give the required product (48 mg, 51% yield) as a white solid.

**$^1$H NMR** (400 MHz, $CDCl_3$) δ = 7.57 (d, *J* = 8.6 Hz, 1H), 6.30 (d, *J* = 8.8 Hz, 1H), 5.05 - 4.85 (m, 1H), 4.01 - 3.82 (m, 1H), 2.23 - 2.06 (m, 4H), 2.04 - 1.83 (m, 2H), 1.70 - 1.56 (m, 2H). **LCMS** (High pH, 2 min) Rt = 1.17 mins, MH- = 270 (100% a/a).

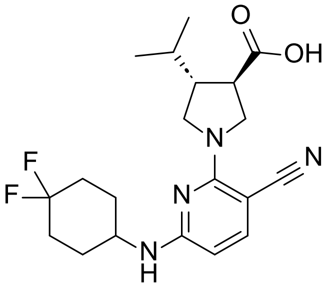

***trans*-1-(3-Cyano-6-((4,4-difluorocyclohexyl)amino)pyridin-2-yl)-4-isopropylpyrrolidine-3-carboxylic acid (14)**

A solution of 2-chloro-6-((4,4-difluorocyclohexyl)amino)nicotinonitrile (48.0 mg, 0.177 mmol), methyl *trans*-4-isopropylpyrrolidine-3-carboxylate (33.3 mg, 0.194 mmol) and DIPEA (0.062 mL, 0.353 mmol) in NMP (0.4 mL) was heated in a microwave at 180 °C for 1 h. The mixture was cooled to rt, then treated with NaOH (2M aq.,

0.4 mL, 0.8 mmol), then the mixture was stirred at 70 °C for 1 h. The reaction mixture was directly purified by MDAP (formic), then the appropriate fractions were combined and concentrated under a stream of nitrogen to give the required product (31 mg, 45% yield) as a brown solid.

**$^1$H NMR** (400 MHz, DMSO-$d_6$) δ = 12.61 (br s, 1H), 7.36 (d, *J* = 8.4 Hz, 1H), 7.19 (br d, *J* = 6.4 Hz, 1H), 5.87 (d, *J* = 8.9 Hz, 1H), 4.00 - 3.80 (m, 3H), 3.70 (dd, *J* = 7.9, 10.8 Hz, 1H), 3.43 (dd, *J* = 8.6, 10.6 Hz, 1H), 3.30 (br s, 3H), 2.86 (q, *J* = 8.4 Hz, 1H), 2.36 - 2.24 (m, 1H), 2.13 - 1.82 (m, 6H), 1.79 - 1.66 (m, 1H), 1.63 - 1.48 (m, 2H), 0.93 (m, 6H). **LCMS** (High pH, 2 min) Rt = 0.81 min, MH+ = 393 (98% a/a).

**Compound 15**

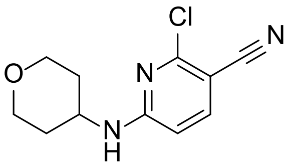

**2-Chloro-6-((tetrahydro-2H-pyran-4-yl)amino)nicotinonitrile (43)**

A solution of 2,6-dichloronicotinonitrile (60 mg, 0.347 mmol), tetrahydro-2*H*-pyran-4-amine (0.036 mL, 0.347 mmol) and DIPEA (0.121 mL, 0.694 mmol) in DMF (0.5 mL) was heated in a microwave at 80 °C for 1 h. The mixture was directly purified by MDAP (high pH), then the appropriate fractions were combined, diluted with DCM (50 mL) and water (50 mL), then the phases were separated using a hydrophobic frit. The organic layer was evaporated in vacuo to give the required product (38 mg, 46% yield) as a white solid.

**$^1$H NMR** (400 MHz, $CDCl_3$) δ = 7.58 (d, *J* = 8.4 Hz, 1H), 6.32 (d, *J* = 8.9 Hz, 1H), 5.02 (br s, 1H), 4.09 - 3.90 (m, 3H), 3.63 - 3.51 (m, 2H), 2.12 - 1.98 (m, 2H), 1.63 - 1.50 (m, 2H). **LCMS** (High pH, 2 min) Rt = 0.94 min, MH- = 236 (100% a/a).

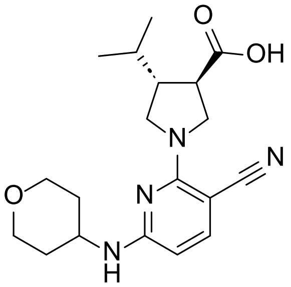

***trans*-1-(3-Cyano-6-((tetrahydro-2H-pyran-4-yl)amino)pyridin-2-yl)-4-isopropylpyrrolidine-3-carboxylic acid (15)**

A solution of 2-chloro-6-((tetrahydro-2*H*-pyran-4-yl)amino)nicotinonitrile (40.0 mg, 0.168 mmol), methyl *trans*-4-isopropylpyrrolidine-3-carboxylate (34.6 mg, 0.202 mmol) and DIPEA (0.059 mL, 0.337 mmol) in NMP (0.4 mL) was heated in a microwave at 180 °C for 30 min. The mixture was directly treated with NaOH (2M aq., 0.2 mL, 0.4 mmol), then the mixture was heated at 70 °C for 1 h. The reaction mixture was directly purified by MDAP (formic) then the appropriate fractions were combined and concentrated under a stream of nitrogen to give the required product (12.1 mg, 20% yield) as an off-white solid.

**$^1$H NMR** (400 MHz, MeCN-$d_3$) δ = 7.33 (d, *J* = 8.3 Hz, 1H), 5.80 (d, *J* = 8.6 Hz, 1H), 5.57 (br d, *J* = 7.1 Hz, 1H), 4.07 - 3.84 (m, 5H), 3.80 - 3.73 (m, 1H), 3.54 - 3.39 (m, 4H), 2.96 - 2.83 (m, 1H), 2.42 - 2.31 (m, 1H), 1.92 - 1.89 (m, 1H), 1.81 - 1.71 (m, 1H), 1.53 - 1.38 (m, 2H), 1.00 - 0.92 (m, 6H). Carboxylic acid proton not observed. **LCMS** (High pH, 2 min) Rt = 0.66 min, MH+ = 359 (99% a/a).

**<u>Compound 16</u>**

**2-Chloro-6-((cyclopropylmethyl)amino)nicotinonitrile (44)**

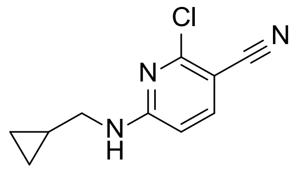

A solution of 2,6-dichloronicotinonitrile (80.0 mg, 0.462 mmol), cylopropylmethylamine (39.9 µL, 0.462 mmol) and DIPEA (161 µL, 0.925 mmol) in DMF (0.5 mL) was heated at 80 °C for 16 h. The reaction mixture was directly purified by MDAP (high pH), then the appropriate fractions were combined and concentrated under a stream of nitrogen to give the required product (70.5 mg, 74% yield) as an off-white solid.

**$^{1}$H NMR** (400 MHz, $CDCl_3$) δ = 7.55 (d, *J* = 8.6 Hz, 1H), 6.28 (d, *J* = 8.6 Hz, 1H), 5.28 (br s, 1H), 3.19 (br t, *J* = 5.7 Hz, 2H), 1.13 - 1.01 (m, 1H), 0.63 - 0.55 (m, 2H), 0.31 - 0.24 (m, 2H). **LCMS** (High pH, 2 min) Rt = 1.10 min, MH+ = 208 (100% a/a).

***trans*-1-(3-Cyano-6-((cyclopropylmethyl)amino)pyridin-2-yl)-4-isopropylpyrrolidine-3-carboxylic acid (16)**

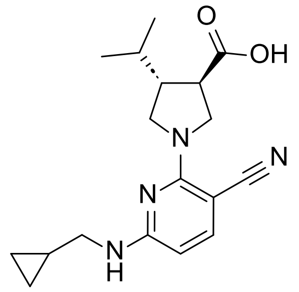

A solution of methyl *trans*-4-isopropylpyrrolidine-3-carboxylate (114 mg, 665 µmol) and DIPEA (174 µL, 997 µmol) in NMP (0.5 mL) was added to 2-chloro-6-((cyclopropylmethyl)amino)nicotinonitrile (69.0 mg, 332 µmol) then the mixture was heated at 150 °C for 2 h, then heated in the microwave at 180 °C for 1 h. The mixture was cooled to rt, then treated with NaOH (10M aq., 166 µL, 1.66 mmol) and heated at 70 °C for 1 h. The reaction mixture was directly purified by MDAP (high pH) then the appropriate fractions were combined and concentrated under a stream of nitrogen. The residue was dissolved in DCM (5 mL) and citric acid (10% aq., 5 mL), then the phases were separated and the organic layer dried under a stream of nitrogen to give the required product (18.7 mg, 17% yield) as an off-white solid.

**$^{1}$H NMR** (400 MHz, Methanol-$d_4$) δ = 7.30 (d, *J* = 8.6 Hz, 1H), 5.85 (d, *J* = 8.6 Hz, 1H), 4.07 (dd, *J* = 8.2, 10.9 Hz, 1H), 3.96 (dd, *J* = 7.9, 10.6 Hz, 1H), 3.81 (dd, *J* = 8.9, 10.9 Hz, 1H), 3.59 - 3.50 (m, 1H), 3.22 (d, *J* = 6.8 Hz, 2H), 2.95 - 2.85 (m, 1H), 2.47 - 2.35 (m, 1H), 1.85 - 1.72 (m, 1H), 1.14 - 1.06 (m, 1H), 1.05 - 0.99 (m, 6H), 0.55 - 0.45 (m, 2H), 0.28 - 0.19 (m, 2H). **LCMS** (High pH, 2 min) Rt = 0.75 min, MH+ = 329 (100% a/a).

**Compound 17**

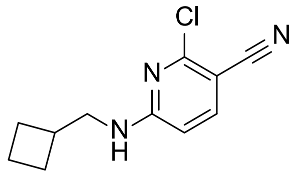

**2-Chloro-6-((cyclobutylmethyl)amino)nicotinonitrile (45)**

A solution of 2,6-dichloronicotinonitrile (30.0 mg, 0.173 mmol), cyclobutylmethylamine (15.0 mg, 0.173 mmol) and DIPEA (91.0 µL, 0.520 mmol) in DMF (1 mL) was heated at 80 °C for 16 h. The reaction mixture was directly purified by MDAP (high pH), then the appropriate fractions were combined and concentrated under a stream of nitrogen to give the required product (23.4 mg, 61% yield) which was used directly in the next reaction.

**LCMS** (High pH, 2 min) Rt = 1.19 min, MH+ = 222 (100% a/a).

***trans*-1-(3-Cyano-6-((cyclobutylmethyl)amino)pyridin-2-yl)-4-isopropylpyrrolidine-3-carboxylic acid (17)**

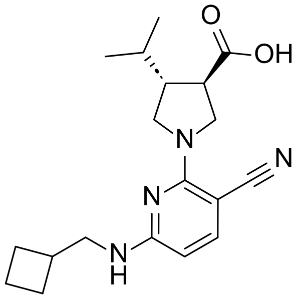

A solution of 2-chloro-6-((cyclobutylmethyl)amino)nicotinonitrile (23.4 mg 0.106 mmol), methyl *trans*-4-isopropylpyrrolidine-3-carboxylate (26 mg, 0.15 mmol), and DIPEA (79 µL, 0.45 mmol) dissolved in DMSO (0.5 mL) was heated in the microwave for at 120 °C for 30 min. Further methyl *trans*-4-isopropylpyrrolidine-3-carboxylate (26 mg, 0.15 mmol) and DIPEA (79 µL, 0.45 mmol) was added then the mixture heated in the microwave at 130 °C for 30 min. The mixture was cooled to rt then NaOH (10 M aq., 100 µL) was added and the mixture heated at 40 °C for 1 h. The reaction mixture was diluted with DMSO (0.4 mL), filtered and purified by MDAP (high pH). The appropriate fractions were combined and concentrated under a stream of nitrogen to give the required product (14.2 mg, 39% yield) as a pale brown solid.

**$^1$H NMR** (400 MHz, Methanol-d$_4$) δ = 7.30 (d, *J* = 8.6 Hz, 1H), 5.83 (d, *J* = 8.6 Hz, 1H), 4.08 (dd, *J* = 8.2, 10.9 Hz, 1H), 3.97 (dd, *J* = 8.1, 10.8 Hz, 1H), 3.85 - 3.77 (m, 1H), 3.58 - 3.50 (m, 1H), 3.38 (d, *J* = 7.1 Hz, 2H), 2.89 (q, *J* = 8.6 Hz, 1H), 2.66 - 2.55 (m, 1H), 2.47 - 2.36 (m, 1H), 2.13 - 2.03 (m, 2H), 1.98 - 1.85 (m, 2H), 1.84 - 1.71 (m, 3H), 1.06 - 0.98 (m, 6H). **LCMS** (Formic, 2 min) Rt = 1.29 min, MH+ = 343 (100% a/a).

**Compound 18**

**2-Chloro-6-((4,4-difluorocyclohexyl)amino)-4-methylnicotinonitrile (46)**

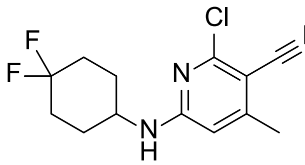

A mixture of 4,4-difluorocyclohexan-1-amine (397 mg, 2.94 mmol), 2,6-dichloro-4-methylnicotinonitrile (500 mg, 2.67 mmol) and DIPEA (1.40 mL, 8.02 mmol) in isopropanol (10 mL) was stirred at 50 °C for 23 h, then at 70 °C for 24 h, then 80 °C for a further 6 h. The reaction mixture was cooled to rt then partitioned between EtOAc (20 mL) and water (20 mL). The organic layer was collected then the aqueous extracted with EtOAc (20 mL). The combined organic phases were washed with brine (25 mL), passed through a hydrophobic frit and concentrated in vacuo. The sample was purified by flash chromatography (Si, 40 g) using a 0-50% EtOAc-cyclohexane gradient over 20 CV. The appropriate fractions were combined and concentrated in vacuo to give the required product (416 mg, 55% yield) as a white solid.

**$^1$H NMR** (400 MHz, DMSO-$d_6$) δ = 7.87 (br d, *J* = 7.3 Hz, 1H), 6.42 (br s, 1H), 2.30 (s, 3H), 2.13 - 1.85 (m, 6H), 1.60 - 1.47 (m, 2H). **LCMS** (High pH, 2 min) Rt = 1.24 min, MH+ = 286/288 (chlorine isotopic distribution, 100% a/a).

***trans*-1-(3-Cyano-6-((4,4-difluorocyclohexyl)amino)-4-methylpyridin-2-yl)-4-isopropylpyrrolidine-3-carboxylic acid (18)**

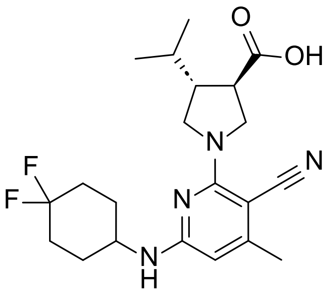

A solution of 2-chloro-6-((4,4-difluorocyclohexyl)amino)-4-methylnicotinonitrile (46 mg, 0.16 mmol) methyl *trans*-4-isopropylpyrrolidine-3-carboxylate (27 mg, 0.160 mmol) and DIPEA (84 µL, 0.48 mmol) in DMSO (0.16 mL). was heated in the microwave 120 °C for 1 h, then at 150 °C for 30 min. The mixture was cooled to rt, NaOH (10 M aq., 0.064 mL, 0.640 mmol) was added then the mixture was heated at 40 °C for 1 h. The mixture was diluted with DMSO (1 mL) and purified by MDAP (high pH), then the appropriate fractions were combined and concentrated under a stream of nitrogen to give the required product (8.6 mg, 12% yield).

**$^1$H NMR** (400 MHz, DMSO-$d_6$) δ = 7.02 (br d, *J* = 5.9 Hz, 1H), 5.79 (s, 1H), 3.95 - 3.78 (m, 3H), 3.71 (dd, *J* = 8.4, 10.8 Hz, 1H), 3.42 (br dd, *J* = 8.6, 11.1 Hz, 1H), 2.83 - 2.73 (m, 1H), 2.30 - 2.22 (m, 1H), 2.15 (s, 3H), 2.13 - 1.79 (m, 6H), 1.75 - 1.64 (m, 1H), 1.61 - 1.47 (m, 2H), 0.91 (m, 6H). Carboxylic acid proton not observed. **LCMS** (Formic, 2 min) Rt = 1.28 min, MH+ = 407 (100% a/a).

**Compound 19**

**Methyl (3*R*,4*R*)-1-(3-cyano-6-((4,4-difluorocyclohexyl)amino)-4-methylpyridin-2-yl)-4-isopropylpyrrolidine-3-carboxylate (47)**

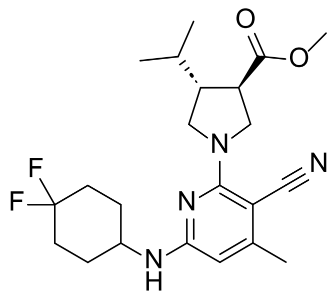

A solution of methyl (3*R*,4*R*)-4-isopropylpyrrolidine-3-carboxylate (264 mg, 1.540 mmol), 2-chloro-6-((4,4-difluorocyclohexyl)amino)-4-methylnicotinonitrile (400 mg, 1.40 mmol) and DIPEA (0.611 mL, 3.50 mmol) in DMSO (10 mL) was heated at 120 °C in the microwave for 8 h. The reaction mixture was partitioned between EtOAc (50 mL) and water (50 mL). The phases were separated then the aqueous was back-extracted with EtOAc (50 mL). The organic layers were combined, washed with brine (50 mL), dried using a hydrophobic frit and concentrated in vacuo. The sample was purified by flash chromatography (Si, 40g) using a 0-100% EtOAc in cyclohexane over 20 CV. The appropriate fractions were combined and evaporated in vacuo to give the required product (458 mg, 78% yield) as a pale-yellow solid.

**$^{1}$H NMR** (400 MHz, DMSO-$d_6$) δ = 7.05 (br d, *J* = 5.9 Hz, 1H), 5.80 (s, 1H), 4.01 - 3.92 (m, 1H), 3.92 - 3.81 (m, 2H), 3.71 (dd, *J* = 8.3, 10.8 Hz, 1H), 3.45 (dd, *J* = 8.8, 10.8 Hz, 1H), 3.29 (s, 3H), 2.97 (q, *J* = 8.6 Hz, 1H), 2.35 - 2.23 (m, 1H), 2.16 (s, 3H), 2.11 - 1.82 (m, 6H), 1.78 - 1.65 (m, 1H), 1.61 - 1.46 (m, 2H), 0.89 (m, 6H). **LCMS** (High pH, 2 min) Rt = 1.45 min, MH+ = 421 (99% a/a).

**(3*R*,4*R*)-1-(3-Cyano-6-((4,4-difluorocyclohexyl)amino)-4-methylpyridin-2-yl)-4-isopropylpyrrolidine-3-carboxylic acid (19)**

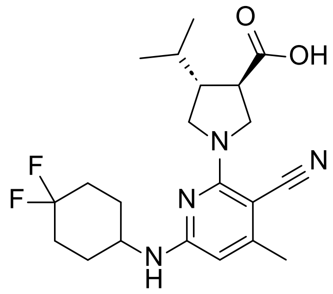

To a stirred solution of methyl (3*R*,4*R*)-1-(3-cyano-6-((4,4-difluorocyclohexyl)amino)-4-methylpyridin-2-yl)-4-isopropylpyrrolidine-3-carboxylate (458 mg, 1.09 mmol) in MeOH (10 mL) was added NaOH (2M aq., 3.0 mL, 6.0 mmol) and the reaction mixture stirred at room temperature for 16 h. Further MeOH (10 mL) and NaOH (2 M aq., 3.0 mL, 6.0 mmol) were added and the reaction mixture stirred at 50 °C for 1 h. The reaction mixture was concentrated in vacuo then dissolved in minimal DMSO and purified by MDAP (high pH), then the appropriate fractions were combined and concentrated under a stream of nitrogen to give the required product (81 mg, 18% yield) as a white solid.

**$^{1}$H NMR** (400 MHz, DMSO-$d_6$) δ = 7.04 (br d, *J* = 6.4 Hz, 1H), 5.79 (s, 1H), 3.98 - 3.78 (m, 3H), 3.71 (dd, *J* = 8.3, 10.8 Hz, 1H), 3.44 (dd, *J* = 8.8, 10.8 Hz, 1H), 2.82 (q, *J* = 8.6 Hz, 1H), 2.34 - 2.21 (m, 1H), 2.16 (s, 3H), 2.11 - 1.80 (m, 6H), 1.77 - 1.64 (m, 1H), 1.62 - 1.46 (m, 2H), 0.92 (m, 6H). Carboxylic acid proton not observed. **LCMS** (High pH, 2 min) Rt = 0.86 min, MH+ = 407 (100% a/a).

**Compound 20**

**Methyl (3*S*,4*S*)-1-(3-cyano-6-((4,4-difluorocyclohexyl)amino)-4-methylpyridin-2-yl)-4-isopropylpyrrolidine-3-carboxylate (48)**

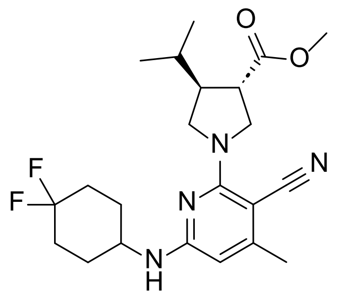

To a solution of 2-chloro-6-((4,4-difluorocyclohexyl)amino)-4-methylnicotinonitrile (100 mg, 0.350 mmol) and methyl (3*S*,4*S*)-4-isopropylpyrrolidine-3-carboxylate (120 mg, 0.700 mmol) in DMSO (1.75 mL) was added DIPEA (0.183 mL, 1.05 mmol), then the mixture was heated in the microwave at 120 °C for 2 h. The reaction mixture was diluted with EtOAc (25 mL) and washed with brine (10 × 25 mL). The organic phase was dried over sodium sulfate, filtered and concentrated *in vacuo*. The sample was purified by flash chromatography (Si, 10g) using a 0-30% EtOAc-cyclohexane gradient. The appropriate fractions were combined and evaporated in vacuo to give the required product (78.7 mg, 50% yield) as a white solid.

**$^1$H NMR** (400 MHz, $CDCl_3$) δ = 5.62 (s, 1H), 4.42 (br d, *J* = 7.3 Hz, 1H), 4.07 (dd, *J* = 8.1, 11.0 Hz, 1H), 3.97 (dd, *J* = 8.1, 11.0 Hz, 1H), 3.85 (dd, *J* = 8.8, 10.8 Hz, 2H), 3.73 (s, 3H), 3.56 (dd, *J* = 9.3, 10.8 Hz, 1H), 2.88 (q, *J* = 8.8 Hz, 1H), 2.47 - 2.36 (m, 1H), 2.29 (s, 3H), 2.18 - 2.02 (m, 4H), 1.97 - 1.79 (m, 2H), 1.79 - 1.69 (m, 1H), 1.66 - 1.54 (m, 2H), 1.00 - 0.92 (m, 6H). **LCMS** (Formic, 2 min) Rt = 1.41 min, MH+ = 421 (100% a/a).

**(3*S*,4*S*)-1-(3-Cyano-6-((4,4-difluorocyclohexyl)amino)-4-methylpyridin-2-yl)-4-isopropylpyrrolidine-3-carboxylic acid (20)**

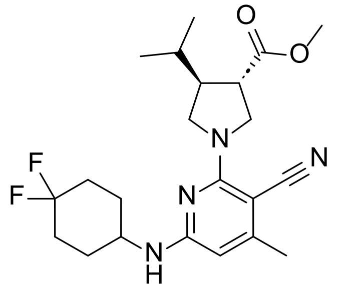

NaOH (2M aq., 0.417 mL, 0.835 mmol) was added to a solution of methyl (3*S*,4*S*)-1-(3-cyano-6-((4,4-difluorocyclohexyl)amino)-4-methylpyridin-2-yl)-4-isopropylpyrrolidine-3-carboxylate (65 mg, 0.155 mmol) in MeOH (0.77 mL), then the mixture was stirred at rt for 16 h. Further NaOH (2M aq., 0.155 mL, 0.309 mmol) was added, then the mixture was stirred for 1 h. The reaction mixture was diluted with EtOAc (20 mL) and the solution washed with brine (3 x 20 mL). The combined aqueous phases were acidified with HCl (2M aq.) to pH = 2, then extracted with EtOAc (20 mL). The organic phases were combined, dried over sodium sulfate and concentrated *in vacuo*. The sample was purified by MDAP (formic), then the appropriate fractions were combined and concentrated in vacuo to give the required product (38.6 mg, 61% yield) as a white solid.

**$^1$H NMR** (400 MHz, Methanol-$d_4$) δ = 5.78 (s, 1H), 4.05 (dd, *J* = 8.3, 10.8 Hz, 1H), 3.98 - 3.89 (m, 2H), 3.82 (dd, *J* = 8.6, 11.0 Hz, 1H), 3.61 - 3.50 (m, 1H), 2.89 (q, *J* = 8.5 Hz, 1H), 2.45 - 2.32 (m, 1H), 2.22 (s, 3H), 2.12 - 1.71 (m, 7H), 1.69 - 1.54 (m, 2H), 1.00 (m, 6H). Exchangeable protons not observed. **LCMS** (Formic, 2 min) Rt = 1.26 min, MH+ = 407 (100% a/a).

**Compound 21**

**2-Chloro-4-methyl-6-((4-methyltetrahydro-2*H*-pyran-4-yl)amino)nicotinonitrile (49)**

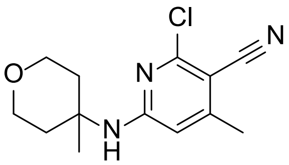

To a solution of 2,6-dichloro-4-methylnicotinonitrile (16 g, 86 mmol) in propylene carbonate (75 mL) at and 2,4,6-trimethylpyridine (12 mL, 90 mmol) was added 4-methyltetrahydro-2*H*-pyran-4-amine (22.2 mL, 174 mmol) were added, then the mixture was heated to 150 °C for 16 h. The temperature was lowered to 70 °C and water (20 mL) was added dropwise, then the mixture was slowly cooled over 2 h to rt, then cooled in an ice bath for 1 h. The resulting precipitate was isolated by filtration and washed with water (50 mL) and triturated with diethyl ether (50 mL) to give the required product (10.0 g, 44% yield) as a red solid.

**$^1$H NMR** (400 MHz, $CDCl_3$) δ = 6.19 (d, *J* = 0.7 Hz, 1H), 4.81 (br s, 1H), 3.81 - 3.57 (m, 4H), 2.37 (d, *J* = 0.7 Hz, 3H), 2.13 - 2.01 (m, 2H), 1.88 - 1.77 (m, 2H), 1.52 (s, 3H). **LCMS** (Formic, 2 min) Rt = 1.09 min, MH+ = 266.

**Methyl (3*R*,4*R*)-1-(3-cyano-4-methyl-6-((4-methyltetrahydro-2H-pyran-4-yl)amino)pyridin-2-yl)-4-isopropylpyrrolidine-3-carboxylate (50)**

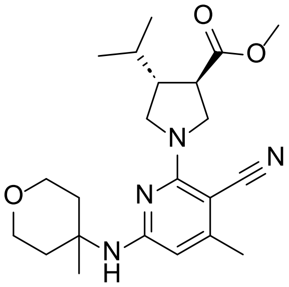

A mixture of 2-chloro-4-methyl-6-((4-methyltetrahydro-2H-pyran-4-yl)amino)nicotinonitrile (4.90 g, 18.4 mmol), methyl (3*R*,4*R*)-4-isopropylpyrrolidine-3-carboxylate hydrochloride (4.60 g, 22.1 mmol) and potassium phosphate, dibasic (12.85 g, 73.80 mmol) in DMSO (25 mL) was heated at 90 °C for 24 h. The mixture was diluted with water (100 mL) and extracted with EtOAc (2 x 100 mL). The organic layers were combined, washed with water (2 x 100 mL) and evaporated in vacuo. The sample was purified by flash chromatography (Si, 120g) using a 0-50% EtOAc/cyclohexane gradient. The appropriate fractions were combined and evaporated in vacuo to give the required product (6.10 g, 83% yield) as a colourless solid.

**$^1$H NMR** (400 MHz, $CDCl_3$) δ = 5.67 (d, *J* = 0.7 Hz, 1H), 4.33 (s, 1H), 4.07 (dd, *J* = 8.2, 10.9 Hz, 1H), 3.98 (dd, *J* = 8.1, 10.8 Hz, 1H), 3.85 (dd, *J* = 8.8, 11.0 Hz, 1H), 3.75 (s, 3H), 3.72 (br d, *J* = 4.9 Hz, 4H), 3.58 (dd, *J* = 8.9, 10.9 Hz, 1H), 2.91 (q, *J* = 8.8 Hz, 1H), 2.50 - 2.37 (m, 1H), 2.28 (s, 3H), 2.23 - 2.10 (m, 2H), 1.83 - 1.71 (m, 3H), 1.52 (s, 3H), 0.98 (m, 6H). **LCMS** (Formic, 2 min) Rt = 1.31 min, MH+ = 401 (100% a/a).

**(3*R*,4*R*)-1-(3-Cyano-4-methyl-6-((4-methyltetrahydro-2H-pyran-4-yl)amino)pyridin-2-yl)-4-isopropylpyrrolidine-3-carboxylic acid (21)**

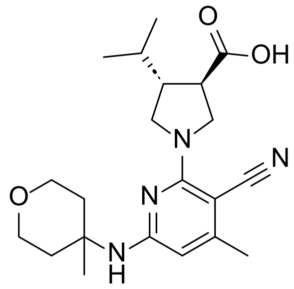

Methyl (3*R*,4*R*)-1-(3-cyano-4-methyl-6-((4-methyltetrahydro-2H-pyran-4-yl)amino)pyridin-2-yl)-4-isopropylpyrrolidine-3-carboxylate (6.10 g, 15.2 mmol) was dissolved in MeOH (90 mL) and NaOH (2M aq., 38 mL, 76 mmol) was added, then the mixture was heated to 50 °C for 2 h. The mixture was cooled and organic solvent removed in vacuo, then the mixture was diluted with water (100 mL) and washed with diethyl ether. The aqueous layer was acidified with HCl (2M aq.) to pH ~4, then extracted with EtOAc (2 x 100 mL) and the combined organics washed with brine, dried and evaporated in vacuo to give the required product (5.80 g, 99% yield) as a colourless solid.

**LCMS** (TFA, 10 min) Rt = 3.93 min, MH+ = 387 (99% a/a).

The compound was mixed with another isolated batch of (3*R*,4*R*)-1-(3-cyano-4-methyl-6-((4-methyltetrahydro-2H-pyran-4-yl)amino)pyridin-2-yl)-4-isopropylpyrrolidine-3-carboxylic acid (13.4 g) then EtOAc (250 mL) was added. The mixture was heated to 70 °C until dissolved, then the mixture was evaporated in vacuo to about half the original volume, resulting in a dense suspension. This was cooled in an ice bath and stirred for 1 h, then the resulting solid was collected by filtration and dried in a vacuum oven at 40 °C to give the required product (15.1 g, 79% yield) as a crystalline solid.

**$^1$H NMR** (600 MHz, DMSO-$d_6$) δ = 12.56 (br s, 1H), 6.70 (s, 1H), 5.91 (s, 1H), 3.87 - 3.92 (m, 1H), 3.82 - 3.76 (m, 1H), 3.71 - 3.65 (m, 1H), 3.59 - 3.55 (m, 2H), 3.55 - 3.49 (m, 2H), 3.42 (dd, *J* = 10.4, 8.9 Hz), 2.85 (q, *J* = 8.4 Hz, 1H), 2.28 (quin, *J* = 8.2 Hz, 1H), 2.21 - 2.15 (m, 2H), 2.14 (s, 3H), 1.71 (dq, *J* = 13.7, 6.8 Hz, 1H), 1.59 - 1.51 (m, 2H), 1.41 (s, 3H), 0.92 (d, *J* = 6.8 Hz, 3H), 0.90 (d, *J* = 6.8 Hz, 3H). **$^{13}$C NMR** (151 MHz; DMSO-$d_6$): δ = 174.9, 158.2, 157.4, 150.5, 120.0, 100.6, 76.0, 63.0, 52.2, 51.7, 50.4, 47.7, 46.0, 36.2 - 36.9 (m, 2C), 29.9, 26.0, 20.6, 20.3, 19.8. **LCMS** (TFA, 10 min) Rt = 3.95 min, MH+ = 387 (100% a/a).

### Analytical Data for compound 21 (GSK235)

#### $^{1}$H NMR spectrum

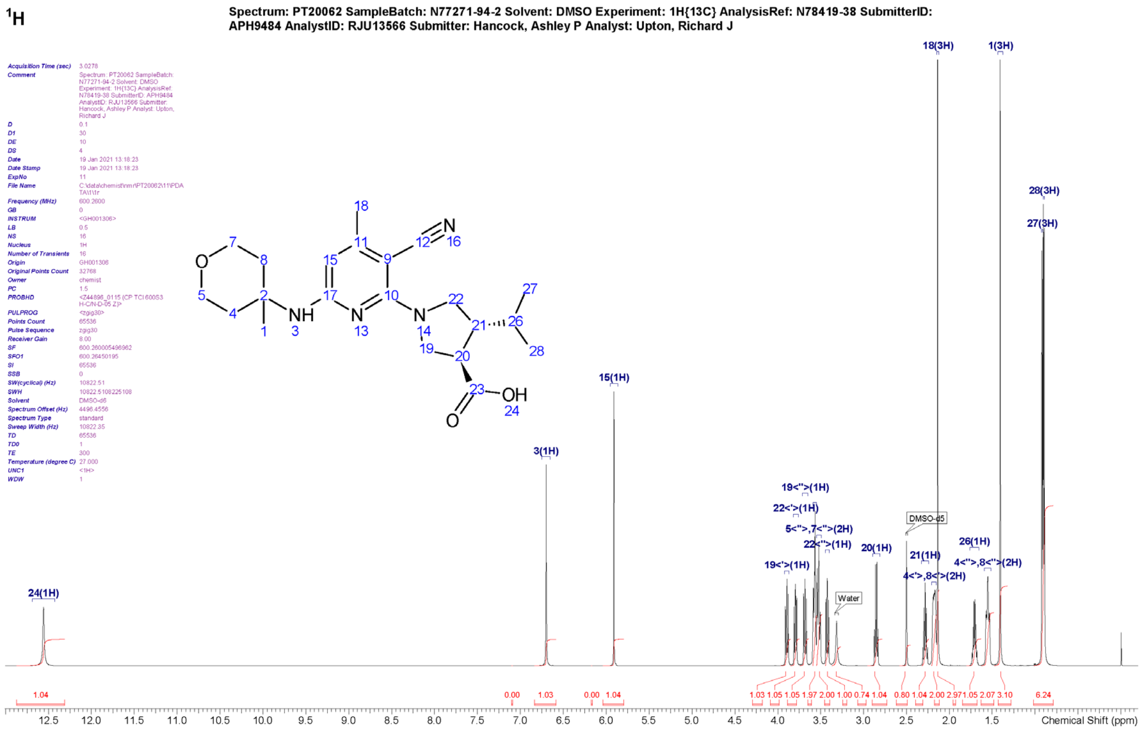

$^{1}$H NMR (DMSO-$d_6$ ,600MHz): δ (ppm) 12.56 (br s, 1H, H-24), 6.70 (s, 1H, H-3), 5.91 (s, 1H, H-15), 3.87 - 3.92 (m, 1H, H-19<'>), 3.76 - 3.82 (m, 1H, H-22<'>), 3.65 - 3.71 (m, 1H, H-19<">), 3.55 - 3.59 (m, 2H, H-5<'>, 7<'>), 3.49 - 3.55 (m, 2H, H-5<">, 7<">), 3.42 (dd, $J$=10.4, 8.9 Hz, 1H, H-22<">), 2.85 (q, $J$=8.4 Hz, 1H, H-20), 2.28 (quin, $J$=8.2 Hz, 1H, H-21), 2.15 - 2.21 (m, 2H, H-4<'>, 8<'>), 2.14 (s, 3H, H-18), 1.71 (dq, $J$=13.7, 6.8 Hz, 1H, H-26), 1.51 - 1.59 (m, 2H, H-4<">, 8<">), 1.41 (s, 3H, H-1), 0.92 (d, $J$=6.8 Hz, 3H, H-27), 0.90 (d, $J$=6.8 Hz, 3H, H-28)

#### $^{1}$H NMR spectrum (magnified)

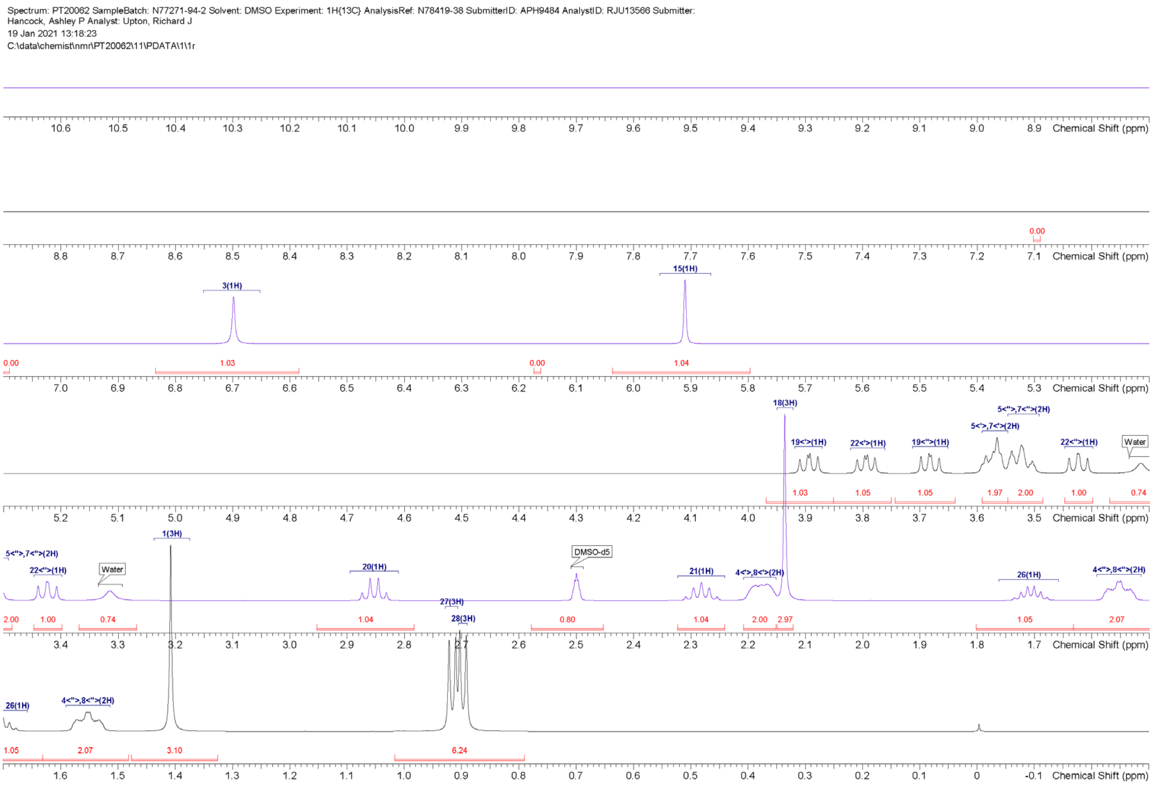

#### $^{13}C$ NMR spectrum

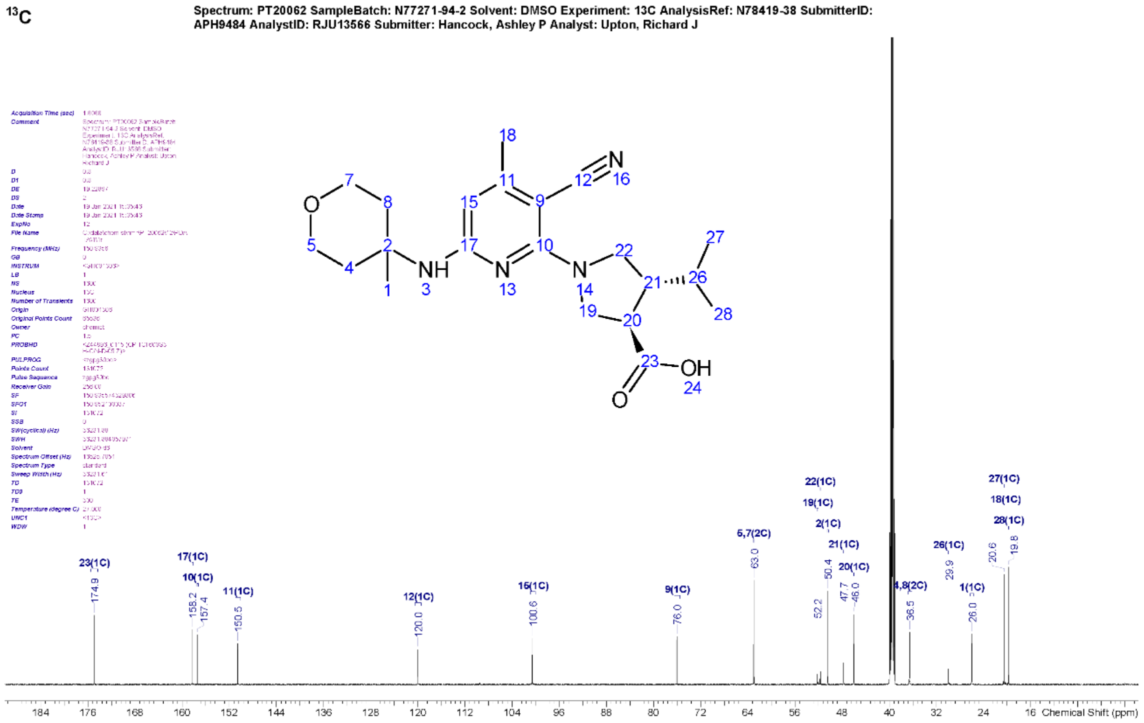

$^{13}C$ NMR (DMSO-$d_6$, 151MHz): δ (ppm) 174.9 (s, C-23), 158.2 (s, C-17), 157.4 (s, C-10), 150.5 (s, C-11), 120.0 (s, C-12), 100.6 (s, C-15), 76.0 (s, C-9), 63.0 (s, C-5, 7), 52.2 (s, C-19), 51.7 (s, C-22), 50.4 (s, C-2), 47.7 (s, C-21), 46.0 (s, C-20), 36.2 - 36.9 (m, C-4, 8), 29.9 (s, C-26), 26.0 (s, C-1), 20.6 (s, C-27), 20.3 (s, C-18), 19.8 (s, C-28)

#### $^{13}C$ NMR spectrum (magnified)

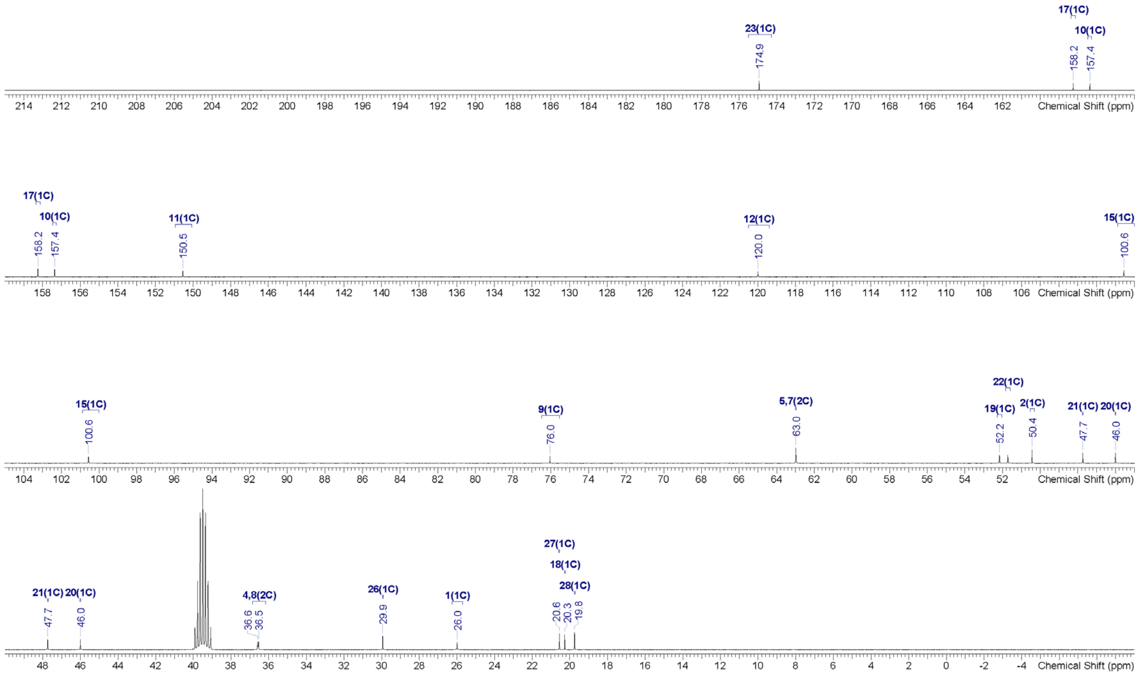

### ROESY NMR spectrum

**ROESY**

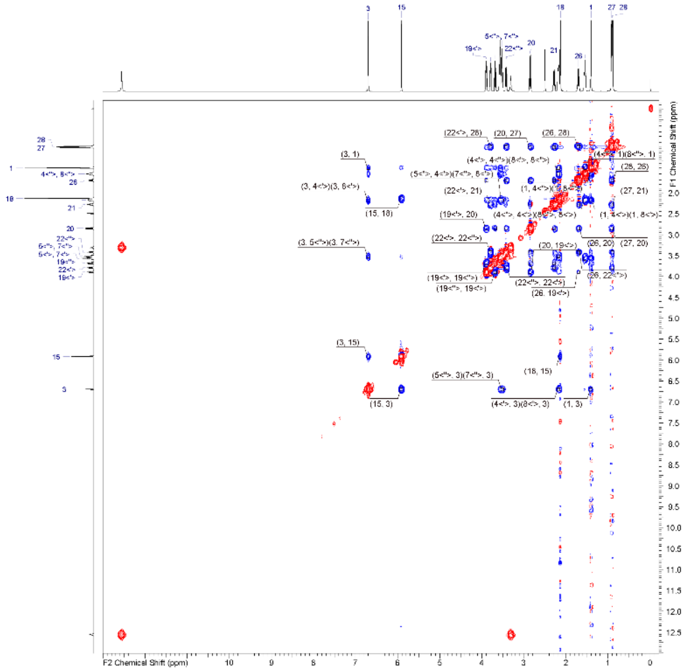

Spectrum: PT20062 SampleBatch: N77271-94-2 Solvent: DMSO Experiment: ROESY AnalysisRef: N78419-38 SubmitterID: APH9484 AnalystID: RJU13566 Submitter: Hancock, Ashley P Analyst: Upton, Richard J

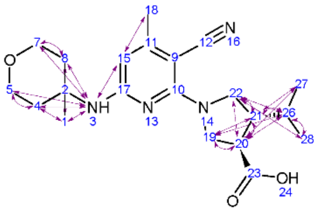

### ROESY NMR spectrum (magnified)

**ROESY**

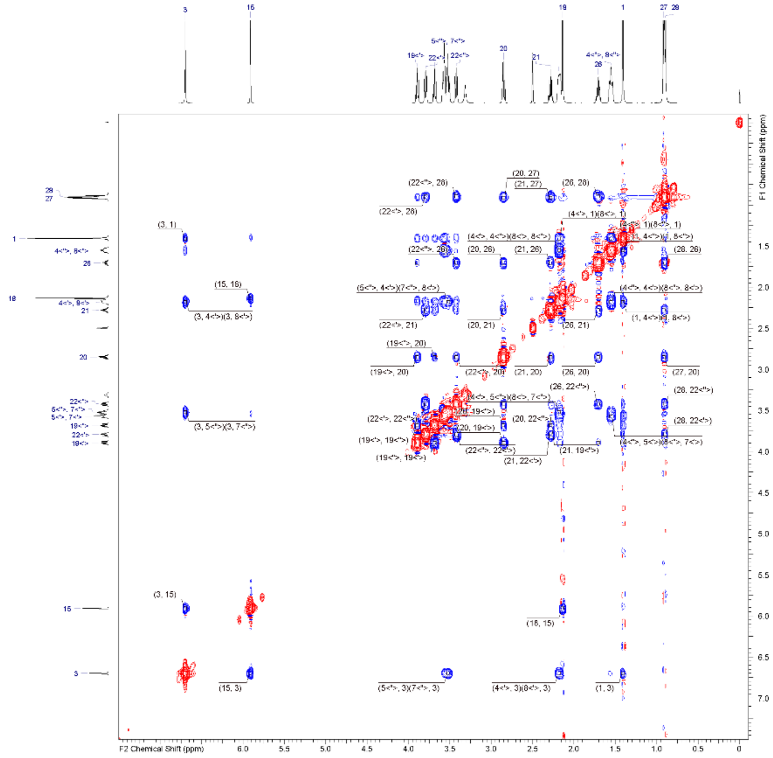

Spectrum: PT20062 SampleBatch: N77271-94-2 Solvent: DMSO Experiment: ROESY AnalysisRef: N78419-38 SubmitterID: APH9484 AnalystID: RJU13566 Submitter: Hancock, Ashley P Analyst: Upton, Richard J

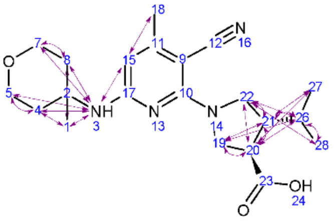

<u>LCMS spectrum (TFA, 10 min)</u>

**Sample: 1**

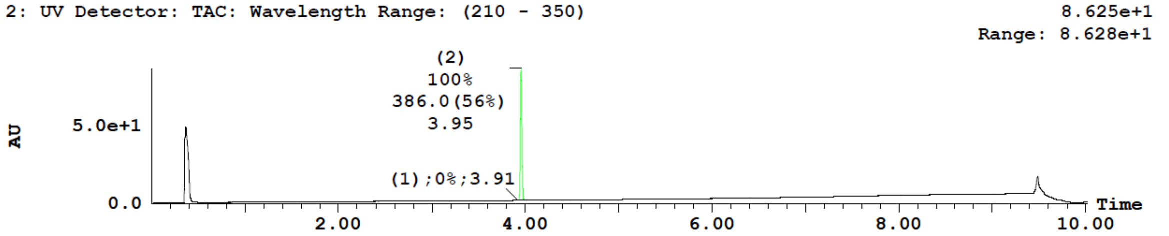

| Peak Number | Time | Area %Total | Mass Found |
|---|---|---|---|
| 1 | 3.91 | 0.14 | Not Found |
| 2 | 3.95 | 99.86 | 386.0000 |

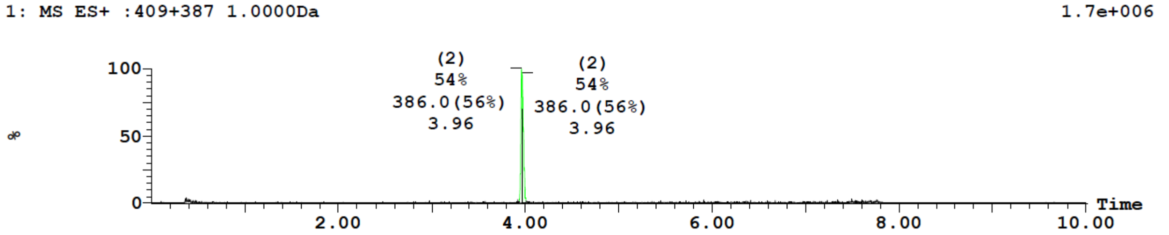

| Peak Number | Time | Mass Found |
|---|---|---|
| 2 | 3.96 | 386.0000 |
| 2 | 3.97 | 386.0000 |

| Peak | Time |
|---|---|
| 1 | 3.91 |

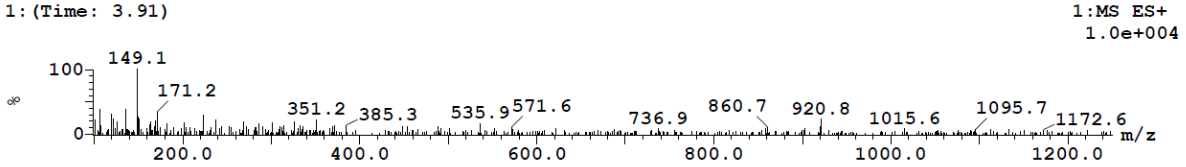

| Peak | Time |
|---|---|
| 1 | 3.91 |

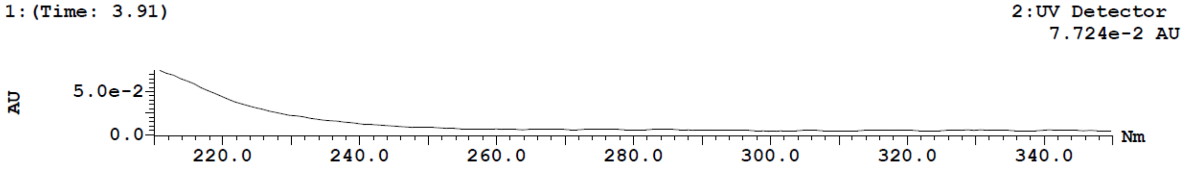

| Peak | Time |
|---|---|
| 2 | 3.96 |

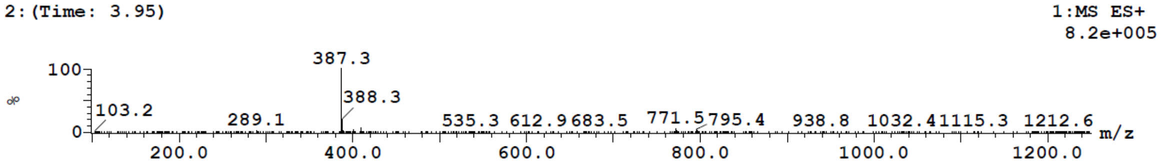

| Peak | Time |
|---|---|
| 2 | 3.96 |

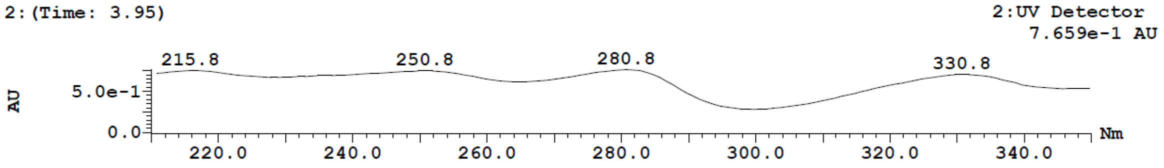

#### ***Ab initio* VCD Analysis of compounds 31 and 32**

Determination of absolute configurations of compounds **31** and **32**

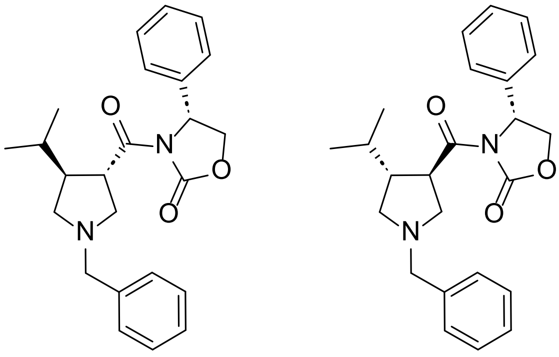

Theoretical Calculation

A conformational search on each of the following structures (fs1rss and fs2rrr) was carried out using MOE at LowMode using MMFF94x forcefield with Born solvation, dielectric constant set at 10 and exterior dielectric constant set at 5.

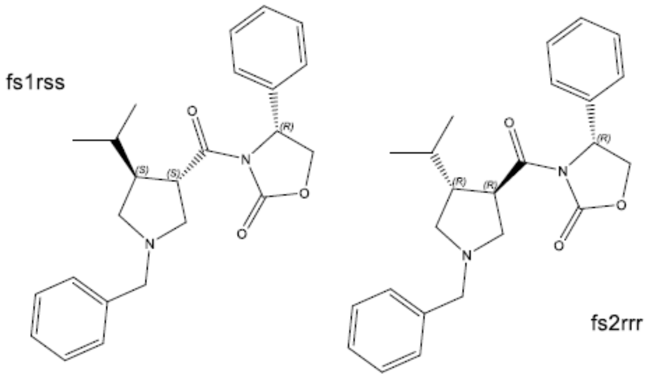

The geometry optimization, frequency, and IR and VCD intensity calculations of the 30 lowest-energy conformers resulted from the conformational search for each structure modelled were carried out at the DFT level (b3lyp/6-31G(d) scrf=(solvent=chloroform)) with Gaussian 16. The Gaussian output files were converted to VCD and IR spectra using BLAIR. The calculated frequencies were scaled by 0.973 and the IR and VCD intensities were converted to Lorentzian bands with 6 $cm^{-1}$ half-width for comparison to experimental spectra. The 18 lowest energy conformers resulting from Gaussian calculations were selected to generate the Boltzmann summed IR and VCD spectra for each modelled structure.

Results: Analysis of Experimental and Calculated Data

Compounds **31** and **32** are diastereomers with two chiral centres. The chiral centre on the oxazolidinone ring is known as (*R*)- in both compounds and the two chiral centres on the pyrrolidine ring are known as relative *trans*- in each compound. The two possible structures were built for the calculations of VCD and IR spectra to be compared with the experimental spectra.

The experimental VCD spectra of compounds **31** and **32** are compared in Figure V1. The major difference in VCD spectra of the two compounds is in the region of 1400-950 $cm^{-1}$. The VCD difference between the two compounds comes from the unknown chiral centre that is opposite in the two isomers. The VCD features that are the same in the two VCD spectra are from the chiral centre that is known as (*R*)- in both compounds.

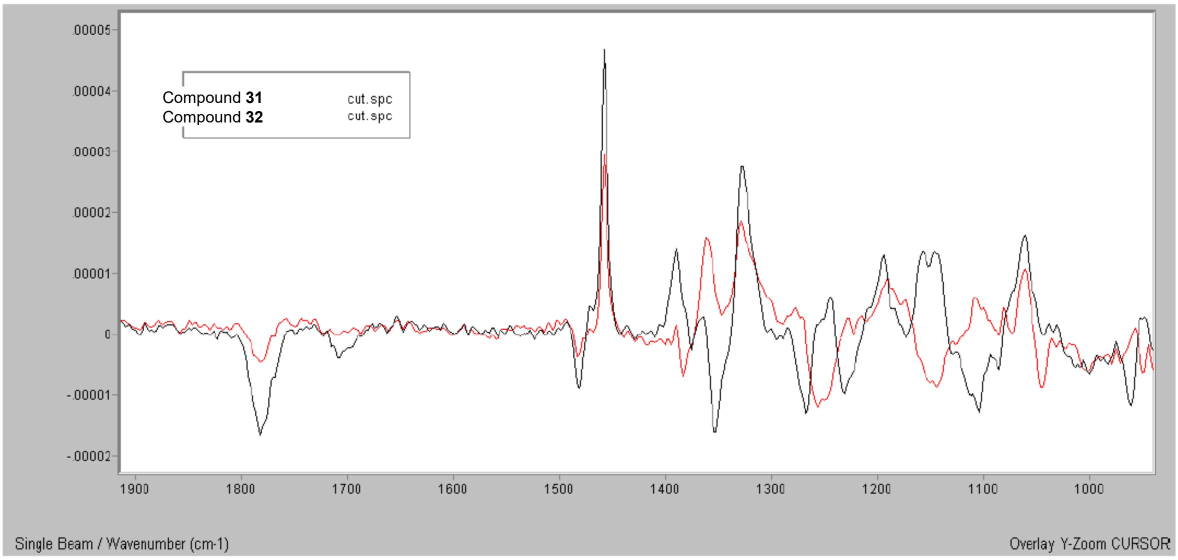

**Figure S1.** Observed baseline corrected VCD spectra of compounds **31** and **32**.

The VCD difference spectrum, obtained by subtracting the VCD spectrum of compound **32** from that of compound **31**, was compared with the VCD difference spectrum of the calculated VCD of fs1rss minus that of fs2rrr in Figure V2. The comparison indicates that the two chiral centres on the pyrrolidine ring are (3*S*,4*S*)- in compound **31** and (3*R*,4*R*)- in compound **32**. The IR spectra for compound **31** and compound **32** are compared in Figure V3 with the calculated IR spectra of the two models. Both the modelled spectra are in good qualitative agreement with experimental, confirming the overall structure of these samples (i.e. its molecular connectivity) and providing additional support for satisfactory coverage of their solution phase conformational space by the computational analysis.

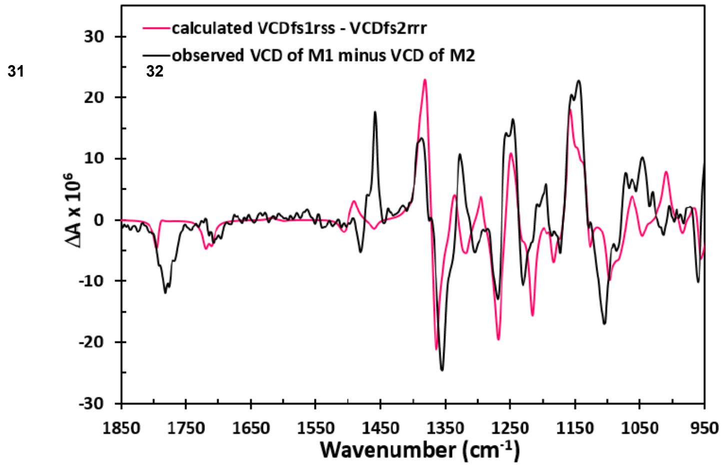

**Figure S2.** Observed VCD difference spectrum between compound **31** and compound **32** compared with the calculated VCD difference spectrum between the fs1rss and fs2rrr configurations.

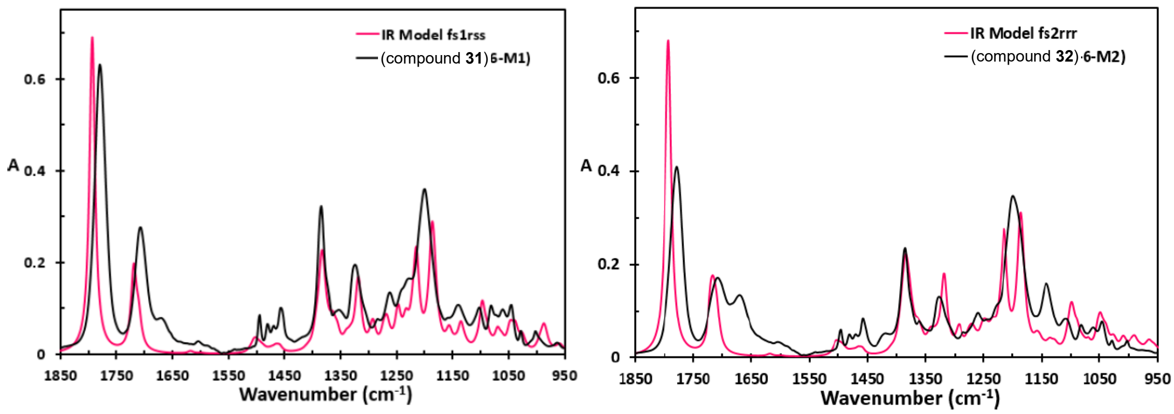

**Figure S3.** IR spectra observed for compound **31** (left) and compound **32** (right) compared with the calculated IR spectra of the fs1rss (left) and fs2rrr (right) configurations
